# Mechanistic Insights into Magnesium Pyrophosphate Formation in the Presence of Gold Nanoclusters Enable Genetic Analysis via Co-Aggregation-Induced Fluorescence Enhancement

**DOI:** 10.64898/2026.08.12.744482

**Authors:** Stylianos Grammatikos, Konstantina Alexaki, Electra Gizeli

## Abstract

The formation of magnesium pyrophosphate (Mg_2_P_2_O_7_) in nucleic acid amplification and cell-free transcription systems has attracted considerable attention, since Mg_2_P_2_O_7_ serves as a reliable indicator of reaction efficiency. However, real-time monitoring of Mg_2_P_2_O_7_ remains challenging, relying largely on time-consuming analytical techniques or end-point detection methods. Here, we report a Mg_2_P_2_O_7_-driven co-aggregation mechanism involving glutathione-capped gold nanoclusters (GSH-AuNCs) that induces fluorescence enhancement, enabling real-time crystal formation monitoring. The mechanism was first investigated in simplified mixtures containing pyrophosphate (P_2_O_7_^4-^) and magnesium (Mg^2+^) ions. Real-time fluorescence profiles revealed that the GSH-AuNCs/Mg_2_P_2_O_7_ co-aggregation can be correlated with crystal formation/growth/solubilization and solution turbidity, while distinct kinetic patterns can be indicative of the crystal size at the end of the reaction. As a next level of complexity, we examined the effects of common components in an enzymatic amplification reaction, *i.e.*, dithiothreitol (DTT), ammonium sulfate ((NH_4_)_2_SO_4_), deoxynucleotides (dNTPs) and Bst polymerase, on Mg_2_P_2_O_7_ formation through real-time GSH-AuNCs fluorescence variations. Guided by the above results, we studied and selected the experimental parameters for the design of an optimized qualitative (end-point) or quantitative (real-time) genetic test. Finally, the loop-mediated isothermal amplification (LAMP) was used as a platform to demonstrate the quantification of Influenza A RNA within the range of 10^2^-10^8^ copies/reaction. The resulting one-tube, contamination-free assay was shown to have a response time of <25 min even in a crude saliva sample. Beyond diagnostics, this crystallization-activated fluorescence strategy may also support real-time investigation of Mg_2_P_2_O_7_ formation in other biotechnological processes, including *in vitro* transcription and Mg_2_P_2_O_7_-bioorganic composites synthesis.

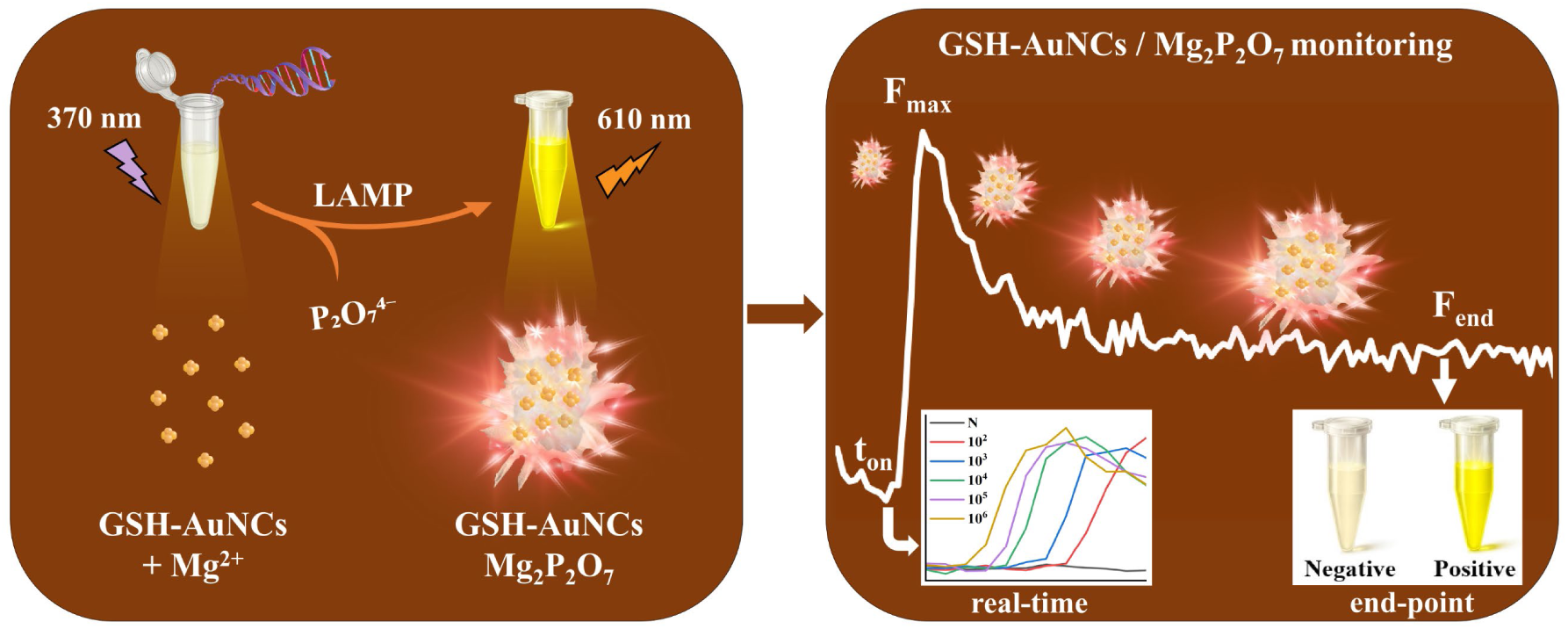

## Introduction

Magnesium pyrophosphate (Mg_2_P_2_O_7_) is a fundamental compound in diverse biochemical and molecular biology processes as well as biotechnological applications of major significance.^1,2^ Its importance is intrinsically linked to the pyrophosphate anion (P_2_O_7_^4-^), an essential by-product of numerous biosynthetic reactions, and magnesium cation (Mg^2+^), a crucial co-factor for the catalytic activity of various enzymes.^3,4^ Mg_2_P_2_O_7_ serves as the substrate of pyrophosphatases, -phosphorylases and -kinases, in processes related to energy metabolism.^5–7^ During nucleic acid synthesis, Mg_2_P_2_O_7_ is formed when released P_2_O_7_^4-^ binds to Mg^2+^ producing an insoluble by-product.^2^ Moreover, Mg_2_P_2_O_7_ crystals have been explored for the formation of hybrid Mg_2_P_2_O_7_-DNA/enzymes composites for therapeutic purposes and efficient delivery of bioactive compounds.^8–13^

In the context of nucleic acid synthesis and *in vitro* transcription (IVT), Mg_2_P_2_O_7_ formation constitutes an important side process, in which an RNA polymerase synthesizes RNA from a DNA template through sequential incorporation of ribonucleotide triphosphates (NTPs), leading to P_2_O_7_^4-^ release.^2,14,15^ This process, used to produce mRNA vaccines or other therapeutic RNAs, is strongly influenced by the formed Mg_2_P_2_O_7_, which often hinders continued RNA production.^7,16^ Therefore, a better understanding of the Mg_2_P_2_O_7_ crystallization mechanism could facilitate the optimization of RNA production processes. Mg_2_P_2_O_7_ formation during IVT has conventionally been investigated using discontinuous and sample-intensive approaches, including turbidity measurements, time-point sampling followed by centrifugation and separate assays of soluble P_2_O_7_^4-^ and Mg^2+^, and microscopic or solid-state characterization of the resulting precipitate.^2,14,16^ More recent methods have enabled rapid at-line monitoring of free Mg^2+7^ or time-lapse imaging of optically detectable crystals in IVT-mimicking microbatch systems.^16^ Nevertheless, these approaches infer crystallization indirectly from changes in solution composition, requiring sample withdrawal and processing, or detect Mg_2_P_2_O_7_ crystals directly after sufficient growth. Consequently, the earliest nucleation events and continuous crystallization dynamics during active IVT remain insufficiently resolved.

Mg_2_P_2_O_7_ is also a key by-product of DNA polymerase-driven amplification, namely the polymerase chain reaction (PCR),^9^ rolling circle amplification (RCA),^17^ loop-mediated isothermal amplification (LAMP),^1^ as well as variants of these methods (*e.g.*, combined with reverse transcription (RT)). In a process resembling IVT, a DNA polymerase incorporates a deoxynucleotide triphosphate (dNTP) into a growing DNA strand, releasing P_2_O_7_^4-^ which interacts with Mg^2+^ to form Mg_2_P_2_O_7_. High DNA amplicon yields produced in RCA^8,18^ and LAMP^1,19^ reactions can lead to the formation of Mg_2_P_2_O_7_ crystal aggregates and a turbid solution. Therefore, turbidity can serve as a sensitive marker for the presence of a genetic target *via* eye observation or monitoring using a turbidimeter, both methods exploited extensively in LAMP reactions.^1,19^ However, while turbidity-based detection of Mg_2_P_2_O_7_ provides a simple label-free readout, it lacks sensitivity and robustness, limiting its applicability primarily to end-point measurements. Mg_2_P_2_O_7_ formation can also be monitored indirectly by using metal ion indicators (hydroxynaphthol blue (HNB), calcein) that interact with Mg^2+^, also enabling visual inspection and low-cost molecular diagnostics.^20,21^ However, such approaches are limited by interference from divalent cations in crude samples, as well as the need for manganese ions (Mn^2+^) in the case of calcein, which may inhibit enzymatic performance.^22^

Recently, ultrasmall nanostructures, such as quantum dots (QDs)^23^, carbon dots (CDs)^24,25^ and gold nanoclusters (AuNCs)^26,27^, have been explored for the fluorescent detection of LAMP-produced Mg_2_P_2_O_7_ (or P_2_O_7_^4-^) based on the use of primarily positively charged nanoprobes. Carbon dots (<10 nm) and ultrasmall (<3 nm) AuNCs have been explored as fluorescent probes due to their unique optical properties, where long fluorescence lifetime is combined with excellent photo/chemical stability and biocompatibility.^28,29^ For example, CDs have been reported to work as probes for the detection of P_2_O_7_^4-^ by-product in a LAMP reaction.^24^ In another study, BSA-AuNCs quenched with copper ions (Cu^2+^) recovered their fluorescence in a positive reaction due to the higher affinity of Cu^2+^ for P_2_O_7_^4-^ released during amplification.^27^ Moreover, cationic polymer polyethylenimine (PEI) coated CDs^25^ or AuNCs^26^ were used for fluorescent detection, through electrostatic interaction with Mg_2_P_2_O_7_. In all the above cases, the particles were inserted inside the LAMP reaction post-amplification, avoiding incompatibility issues with the LAMP reaction. Despite the above advances made for the direct or indirect detection of Mg_2_P_2_O_7_, these approaches are primarily limited to end-point measurements, reducing considerably the general applicability of the techniques within and beyond diagnostics.

An interesting extension of fluorescent metal NCs is that they exhibit aggregation-induced-emission (AIE) enhancement characteristics, by restriction of intramolecular vibration and intermolecular rotation of the surface motifs of NCs, minimizing nonradiative decay.^30–32^ Several studies have demonstrated this phenomenon, such as the transition of non-fluorescent water-soluble glutathione (GSH)-capped AuNCs (GSH-AuNCs) to highly luminescent aggregates in the presence of ethanol,^33^ or the formation of peptide-stabilized AuNCs with a pronounced AIE enhancement at pH 5.^34^ However, until today, such a method has not been described in the context of Mg_2_P_2_O_7_ co-aggregation.

To address the challenge of monitoring the formation of Mg_2_P_2_O_7_, we explored the use of negatively charged green-synthesized GSH-AuNCs which are compatible with a nucleic acid enzymatic reaction producing Mg_2_P_2_O_7_. GSH, a naturally occurring tripeptide, exhibits exceptional metal coordination ability and compatibility with *in vivo* and *in vitro* experiments.^35,36^ Importantly, GSH-AuNCs were selected following our discovery that they co-precipitate with Mg_2_P_2_O_7_ leading to an increase in fluorescence though a mechanism resembling AIE enhancement. Initially, we investigated the formation of Mg_2_P_2_O_7_ in reactions containing only the two inorganic substances (Mg^2+^ and P_2_O_7_^4-^), and then in the presence of key components of an enzymatic amplification reaction, *i.e.*, dithiothreitol (DTT), ammonium sulfate ((NH_4_)_2_SO_4_) and dNTPs. These studies provided valuable insight into the Mg_2_P_2_O_7_ crystallization/solubilization process as a function of the [Mg^2+^]/[P_2_O_7_^4-^] ratio, in the absence or presence of additives, while SEM and absorbance measurements provided complementary information on the crystals’ size and solution turbidity, respectively. The above mechanistic insights were further used to design an optimized genetic analysis assay based on the LAMP method, demonstrated to detect down to 100 copies per reaction within 25 minutes, even in crude saliva samples. This work provides the first real-time monitoring of Mg_2_P_2_O_7_ formation in an enzymatic amplification reaction based on co-aggregation-induced fluorescence enhancement of GSH-AuNCs, while it sets the foundation for its application in other biotechnological processes leveraging Mg_2_P_2_O_7_ formation.

## Results and Discussion

### GSH-AuNCs synthesis and fluorescence emission enhancement induced by co-aggregation with Mg_2_P_2_O_7_

Our work involved first the synthesis of GSH-AuNCs *via* a one-step procedure, as shown in **Scheme 1**. The ultraviolet-visible (UV-Vis) spectrum showed a weak absorption band at ∼400 nm, while fluorescence spectroscopy (FS) measurements revealed a distinct emission peak at 610 nm (upon excitation at 370 nm) (**Fig. 1a**), both consistent with size-dependent electronic transitions within the small-sized AuNCs.^37^ GSH molecules, known to act simultaneously as a stabilizing ligand and a mild reducing agent,^38^ can strongly coordinate with Au atoms through thiol groups, forming stable Au-S covalent bonds. The carboxyl and amine groups of the GSH further participate in the electronic interactions with the gold core, influencing the optical properties and enhancing fluorescence.^39^ Further characterization of GSH-AuNCs using attenuated total reflectance-Fourier transform infrared spectroscopy (ATR-FTIR), zeta potential (ZP) measurements and transmission electron microscopy (TEM) imaging confirmed the successful formation of spherical, negatively charged AuNCs capped with GSH, with a calculated mean diameter of 2.2±0.4 nm (**Fig. S1a-d**).

**Scheme 1:**
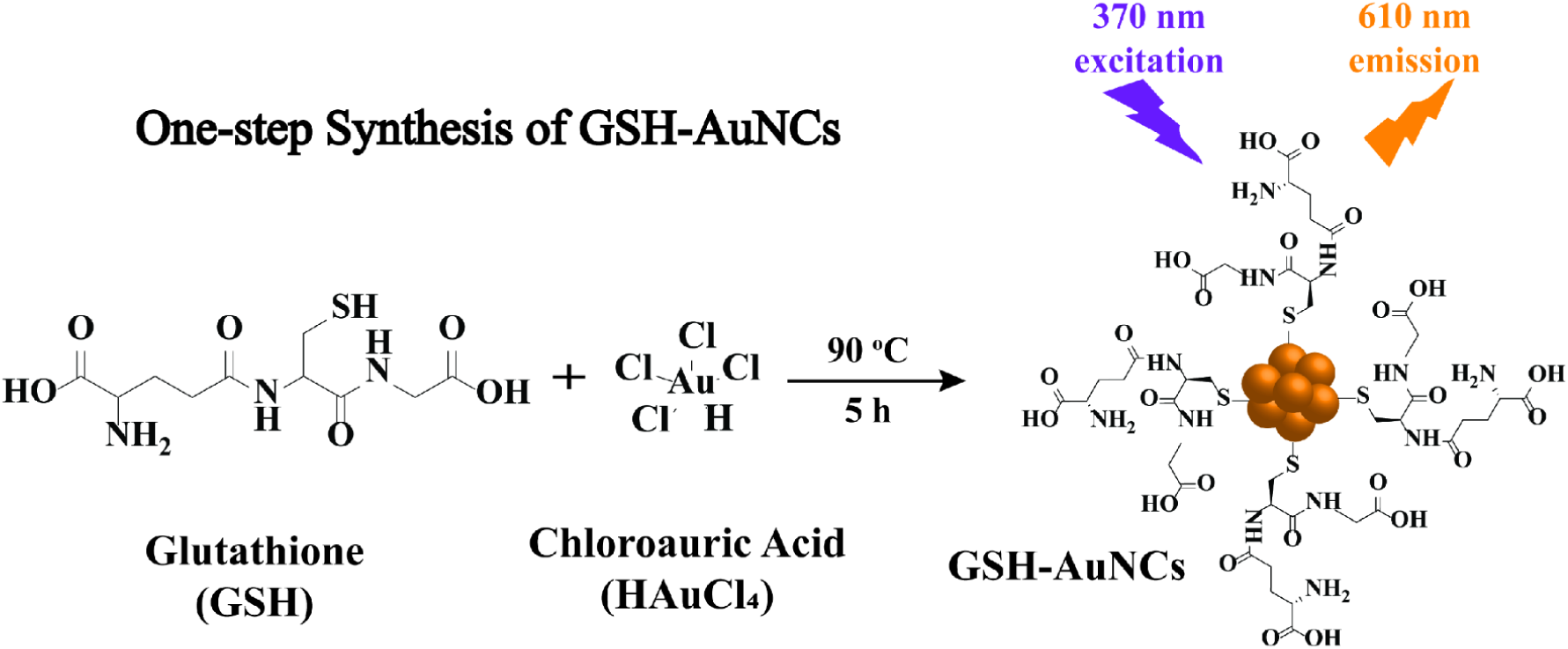
Schematic illustration of the one-step synthesis of GSH-AuNCs.

In a follow-up step, we investigated the interaction of GSH-AuNCs with Mg^2+^ (8 mM), P_2_O_7_^4-^ (4 mM), and both of them (8 mM Mg^2+^, 4 mM P_2_O_7_^4-^) in Tris buffer (**Fig. 1b**). Our results show that only the combination of Mg^2+^ with P_2_O_7_^4-^ induces a significant fluorescence increase, a result also confirmed by visual inspection of the reaction solutions under UV light. Further characterization of the formed crystalline precipitate using ATR-FTIR and energy-dispersive X-ray spectroscopy (EDX) confirmed the presence of GSH-AuNCs in the crystals (**Fig. S2a-c**). This led us to assume that the co-aggregation of GSH-AuNCs with Mg_2_P_2_O_7_ induces fluorescence enhancement, through an AIE-like enhancement mechanism.^40–43^ In a previous study, the AIE fluorescence increase of GSH-AuNCs complexes induced by multivalent cations such as cadmium ions (Cd^2+^) was attributed to electrostatic and coordination interactions between the cation and the monovalent carboxylic anions of GSH to form inter- and/or intra-complex cross-links.^44^ A similar AIE-like increase in the fluorescence of GSH-AuNCs has also been reported when they are complexed with aluminium ions (Al^3+^).^45–47^ In our case, the presence of 8 mM Mg^2+^ alone was not enough to induce significant fluorescence enhancement. However, in the presence of both Mg^2+^ and P_2_O_7_^4-^, it appears that Mg^2+^ can stabilize the coordination of the GSH-AuNCs with the P_2_O_7_^4-^ in the formed Mg_2_P_2_O_7_ crystals. This effect may allow the GSH-AuNCs to come into close proximity in a geometry that induces ligand-to-metal and ligand-to-metal-metal charge transfer (LMCT and LMMCT).^30,44^ Notably, the fluorescence-enhanced solution exhibited a blue-shift of the emission maximum from 610 nm to 604 nm, together with a narrowing of the full width at half maximum (FWHM) from 116.3±1.2 nm to 111.4±1.7 nm. Such spectral changes suggest a more rigid environment around the NCs, typical of AIE systems, where restriction of motion leads to higher-energy emission.^48^ In contrast, when GSH-AuNCs were embedded in NaCl crystals, a small red-shift (∼4 nm) was reported, attributed to the change of dielectric constant in the surrounding medium.^49^ In that case, NaCl confinement enhanced the radiative decay rate while suppressing non-radiative losses, leading to strong luminescence enhancement. Taken together, these findings suggest that Mg_2_P_2_O_7_ enhances GSH-AuNCs fluorescence primarily through restriction of motion within a rigid crystalline environment, aligning closely with an AIE-type mechanism. X-ray diffraction (XRD) spectroscopy was also employed to study the crystalline phase of the formed Mg_2_P_2_O_7_, revealing a hydrate phase (Mg_2_P_2_O_7_ ⋅ 3.5H_2_O) (**Fig. S2d**) as already reported in the literature.^16^

We further investigated the dependence of GSH-AuNCs fluorescence enhancement on the formed Mg_2_P_2_O_7_ under varying [P_2_O_7_^4-^], by measuring end-point fluorescence. As-synthesized GSH-AuNCs were incubated with Mg^2+^ (8 mM) and various [P_2_O_7_^4-^] (0-10 mM) in Tris buffer. As shown in **Fig. 1c**, increasing [P_2_O_7_^4-^] up to 4 mM significantly enhanced the fluorescence signal relative to the 0 mM (control). However, at higher concentrations (5-10 mM), fluorescence gradually decreased, revealing a concentration-dependent inverse response. The dependence of the fluorescence enhancement on the [P_2_O_7_^4-^] was further corroborated by visual observation under UV illumination. The addition of 1-4 mM P_2_O_7_^4-^ transformed the weakly colored GSH-AuNCs solution into a brightly colored one under UV light (**Fig. 1d-top**), consistent with enhanced fluorescence. The formation of a colored precipitate with a clear supernatant after a spin-down step supports the co-aggregation of GSH-AuNCs with the Mg_2_P_2_O_7_ crystals. In contrast, solutions incubated with >4 mM [P_2_O_7_^4-^] gradually lost fluorescence enhancement, reverting to their original weak color under UV light, while the amount of Mg_2_P_2_O_7_ precipitate decreased. In this [P_2_O_7_^4-^] range, the supernatant produced after spin-down was not clear, indicating release of GSH-AuNCs from the Mg_2_P_2_O_7_ aggregates. A similar trend was observed under visible light, with the precipitate (**Fig. 1d-bottom**) significantly dissolved at higher [P_2_O_7_^4-^] (> 8 mM). Plotting the relative fluorescence intensity (F/F_0_) (from **Fig. 1c**) as a function of [P_2_O_7_^4-^] revealed a 4.7-fold increase at a [Mg^2+^]/[P_2_O_7_^4-^] ratio equal to 2 (**Fig. 1e**), corresponding to the stoichiometric ratio of the two inorganic components for maximum pellet formation, according to: 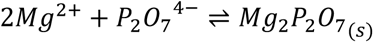

The turbidity of the solutions was also recorded at the end of the reaction by measuring absorption at visible wavelengths,^50^ where turbidity can indicate the amount of the formed precipitate.^1^ As shown in **Fig. 1f**, turbidity increased with [P_2_O_7_^4-^] up to 4 mM, indicating enhanced Mg_2_P_2_O_7_ formation, and then decreased at higher concentrations due to gradual solubilization of the precipitate (Mg_2_P_2_O_7_), with almost complete solubilization occurring at >8 mM P_2_O_7_^4-^. Comparison of F/F_0_ (**Fig. 1e**) with turbidity (**Fig. 1f**) reveals overall similar, but not identical trends.

To rationalize the non-monotonic dependence of Mg_2_P_2_O_7_ precipitation on [P_2_O_7_^4-^], we considered the following equations depicting the formation of Mg_2_P_2_O_7._ Free Mg^2+^ and P_2_O_7_^4-^ first chelate to form the more stable 1:1 MgP_2_O_7_^2-^ complex, followed by the formation of the neutral 2:1 Mg_2_P_2_O_7_, according to:

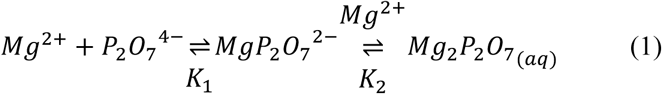

with reported stability constants K_1_=10^5.41^ M^-1^ and K_2_=10^2.34^ M^-1^, respectively, under alkaline conditions.^51^ Similar constants have also been reported in a recent study.^16^ In addition, precipitation of Mg_2_P_2_O_7_ occurs once the ion product:

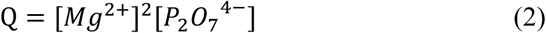

exceeds the solubility product constant (K_sp_), where its apparent value for Mg_2_P_2_O_7_ ⋅ 3.5H_2_O, consistent with the phase identified by XRD (**Fig. S2d**), can be taken as K_sp_ = 1.63⋅10^-13^ M^3^ at 25 °C.^16^ Based on the above equations, once the maximum production of Mg_2_P_2_O_7_ crystals is reached at a ratio of [Mg^2+^]/[P_2_O_7_^4-^]=2, any further excess of free P_2_O_7_^4-^ will compete for the crystal-bound Mg^2+^, leading gradually to the solubilization of the crystals and formation of the stable MgP_2_O_7_^2-^ soluble complex, in agreement with our end-point fluorescence (**Fig. 1e**) and turbidity (**Fig. 1f**) measurements. Note that this behavior, *i.e.*, reverse solubility of Mg_2_P_2_O_7_ as a function of the [P_2_O_7_^4-^], has been reported before.^14,16^

Further insights on the interaction of GSH-AuNCs with Mg_2_P_2_O_7_ crystals were obtained using confocal microscopy imaging (**Fig. S3a**). The same amount of GSH-AuNCs was added either after the interaction of 8 mM Mg^2+^ with 4 mM P_2_O_7_^4-^ was completed (*i.e.*, post Mg_2_P_2_O_7_ formation), or *in situ* during the Mg_2_P_2_O_7_ formation. In the latter case, more GSH-AuNCs seem to be bound into the crystals, resulting in higher end-point fluorescence enhancement. This is also indicated by photographs of the solutions under UV light **(Fig. S3b**), where no fluorescence in the supernatant was observed after spin-down during the *in situ* formation, contrary to the preformed Mg_2_P_2_O_7_ case. This result implies that GSH-AuNCs may be entrapped inside Mg_2_P_2_O_7_ crystals, while in the preformed Mg_2_P_2_O_7_ case, they may only be bound onto the surface of the formed crystals.^23^

**Figure 1:**
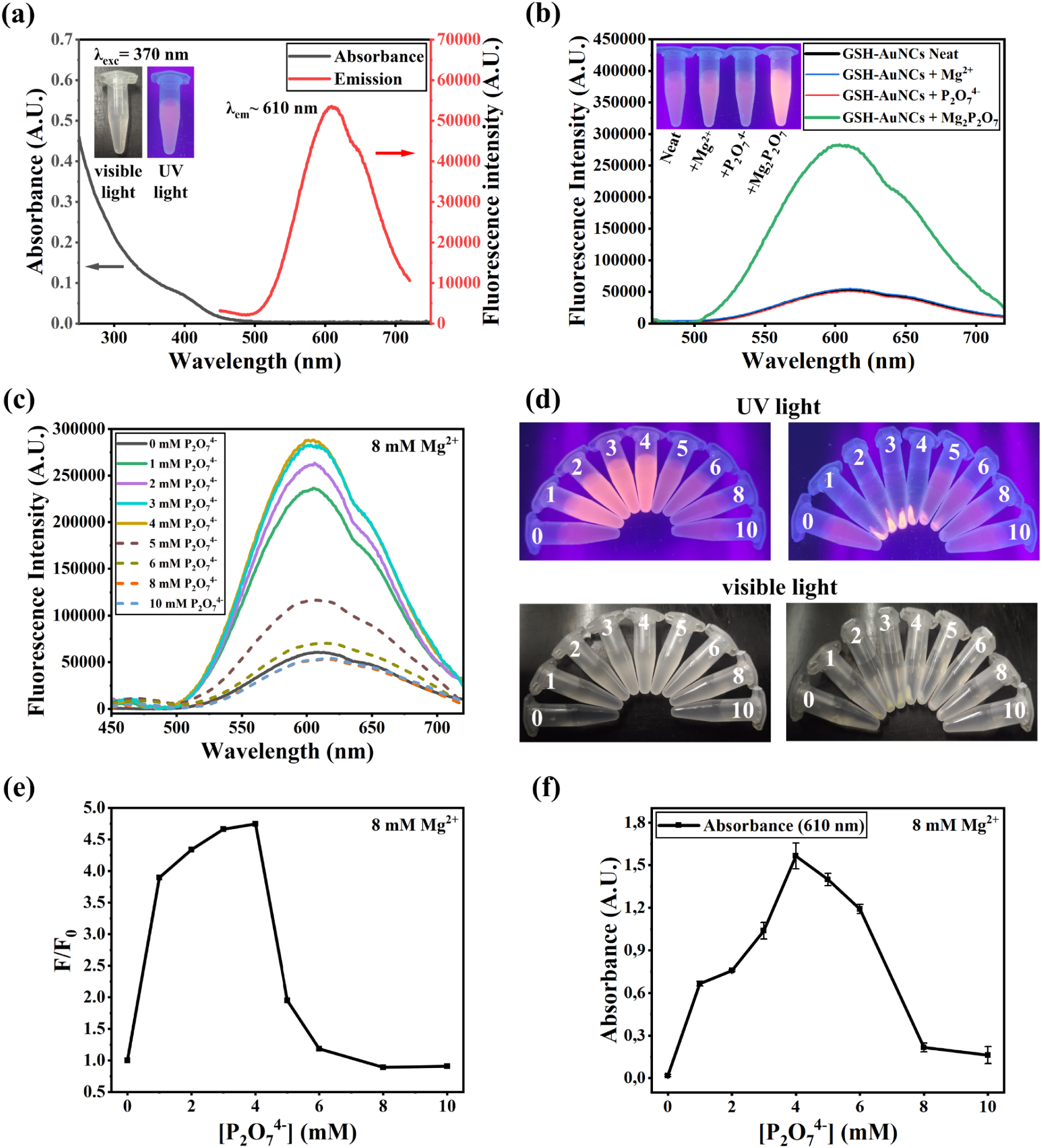
(a) UV-Vis absorption and FS emission spectra of GSH-AuNCs. Inset: Photographs of the GSH-AuNCs solution under visible light (left) and UV light (right). (b) End-point FS measurements of GSH-AuNCs in the presence of neat Mg^2+^ (8 mM), P_2_O_7_^4-^ (4 mM), or both. Inset: Photograph of the corresponding solutions under UV light. (c) FS measurements of GSH-AuNCs solutions prepared with 8 mM Mg^2+^ and varying concentrations of P_2_O_7_^4-^ (0-10 mM). (d) Photographs of the solutions from (c) under UV light (top) and visible light (bottom), before (left) and after (right) spin-down. (e) Relative fluorescence intensity (F/F_0_) of the spectra in (c), as a function of [P_2_O_7_^4-^]. (f) Absorbance measurements at 610 nm for the solutions in (c), a measure of the solution turbidity as a function of [P_2_O_7_^4-^]. All solutions were incubated at 65 °C for 60 min in Tris (20 mM, pH 8), in the presence of 25-fold diluted GSH-AuNCs.

### Real-time study of Mg_2_P_2_O_7_ formation in the presence of GSH-AuNCs

The incorporation of GSH-AuNCs in the Mg_2_P_2_O_7_, resulting in fluorescence enhancement, opens the opportunity for the *in situ* study of Mg_2_P_2_O_7_ crystal formation. To investigate this process and assess the robustness of GSH-AuNCs as a sensitive fluorescence probe, we tested the simplest case where various [P_2_O_7_^4-^] (0-10 mM) were incubated with 4, 8 or 12 mM Mg^2+^ in Tris buffer, together with 1 μL of the synthesized GSH-AuNCs in 25 μL total volume reactions. The three selected [Mg^2+^] were intended to cover the range present in IVT^2,14,16^ or isothermal amplification reactions (*e.g.*, LAMP and RCA),^1,18^ while for better simulation of an enzymatic amplification process, experiments were performed at 65 °C (simulated LAMP).^52,53^

**Fig. 2a-c** provide the combined response of the solutions’ fluorescence as a function of the [P_2_O_7_^4-^] and reaction time. Inspection of the above graphs reveals as points of interest the onset of Mg_2_P_2_O_7_ production (t_on_) and a sharp fluorescence peak observed in some cases. To enable better comparison of Mg_2_P_2_O_7_ formation under the various experimental conditions, we also plotted on the same graph the fluorescence response obtained for each [P_2_O_7_^4-^] when incubated with the three different [Mg^2+^] (**Fig. S4**). **Fig. 2d** depicts four representative real-time responses: at low [P_2_O_7_^4-^] (0.5 mM), fluorescence begins to increase rather late (t_on_ ∼ 31 min) and gradually reaches a plateau. In contrast, higher P_2_O_7_^4-^ concentrations produce traces in which fluorescence initially rises to a F_max_ and then decreases before stabilization occurs (F_end_). This response can be smooth and fast (*e.g.*, 1 mM P_2_O_7_^4-^, t_on_ ∼ 14 min) or more pronounced and even faster (*e.g.*, 2 mM P_2_O_7_^4-^, t_on_ ∼ 5 min), with both cases exhibiting a distinct “hat-shaped” curve. At even higher [P_2_O_7_^4-^] (*e.g.*, 10 mM), the fluorescence trace mostly overlaps with the 0 mM control, consistent with MgP_2_O_7_^2-^ becoming the dominant species in the solution.

#### F_max_ and F_end_ as a function of [P_2_O_7_^4-^]

The above results reveal F_max_ and F_end_ as the main parameters of interest in real-time curves. When we plotted the F_max_/F_0_ against [P_2_O_7_^4-^] for each one of the curves depicted in **Fig. 2a-c**, we obtained the highest response at a [P_2_O_7_^4-^] that corresponds to half of the [Mg^2+^] in solution (**Fig. 2e**), or at a [Mg^2+^]/[P_2_O_7_^4-^] ratio that equals 2 (**Fig. S5a)**. Closer observation of **Fig. 2e** reveals that for 8 mM Mg^2+^, the response follows the absorbance measurements at end-point, as this is depicted in **Fig. 1f**. Assuming that this is true for all the [Mg^2+^] tested here, we can propose that the F_max_ values recorded during the course of the interaction of the two ions in the presence of GSH-AuNCs follow the formation of Mg_2_P_2_O_7_ aggregates. This is supported by the (calibrated) 14 times higher fluorescence enhancement observed during the incubation of 12 mM Mg^2+^ with 6 mM P_2_O_7_^4-^, compared to the 10 and 4 times enhancement in the presence of 8 mM Mg^2+^/ 4 mM P_2_O_7_^4-^ and 4 mM Mg^2+^/ 2 mM P_2_O_7_^4-^, respectively, in agreement with the production of a higher amount of crystal aggregates as the concentration of the initial reactants increases. Moreover, since the process is monitored through the Mg_2_P_2_O_7_ co-aggregation with GSH-AuNCs, **Fig. 2e** suggests that fluorescence enhancement is proportional to the amount of produced Mg_2_P_2_O_7_ crystal aggregates. However, with the exception of the lowest [P_2_O_7_^4-^] (0.5 mM), the F_max_ does not plateau; instead, a transient optical response is recorded in all cases where solid Mg_2_P_2_O_7_ is formed. As a result, the F_end_/F_0_ against [P_2_O_7_^4-^] graph (**Fig. 2f**) is distinctly different from **Fig. 2e**, both qualitatively and quantitatively. In this case, for the 8 mM Mg^2+^ the calibrated F_end_ flattens at a level which is only ∼2.5 times higher than the background fluorescence for the 0.5-4 mM [P_2_O_7_^4-^] range. The same trend is depicted with the 12 mM Mg^2+^. A lower F_end_ is also shown for the 4 mM Mg^2+^, although here the F_end_/F_0_ follows a gradual decrease with [P_2_O_7_^4-^] (**Fig. 2f** and **Fig. S5b**). The fluorescence of the solutions shown in **Fig. 2f** under UV light and after 60 min of the reaction is also depicted in **Fig. 2g**.

#### F_max_ and F_end_ indicate stages in the crystallization/aggregation process

To explain the kinetics and end-point fluorescence of the various solutions tested, we looked into the mechanism of Mg_2_P_2_O_7_ formation. According to the literature,^2,14^ Mg_2_P_2_O_7_ formation begins when the monomers (P_2_O_7_^4-^ and Mg^2+^) produce stable clusters (precursors) in the solution. After a critical concentration point, the growth of the generated stable nuclei is induced under supersaturated conditions, leading to Mg_2_P_2_O_7_ crystal growth and increased solution turbidity. In a previous study, it was reported that turbidity can produce a decline in fluorescence intensity during the later stage of an enzymatic amplification reaction (LAMP), also giving a “hat-shaped” real-time curve.^54^ Other studies, reporting that light scattering in a turbid solution (*i.e.*, in the presence of aggregates) leads to fluorescence attenuation, also support the above conclusion.^55^ Based on the above evidence and assuming that GSH-AuNCs co-aggregation follows closely the Mg_2_P_2_O_7_ crystal formation process, we may suggest that the signal increase up to F_max_ traces the first stages of the process up to nucleation and formation of the first aggregates. To explain the sharp fluorescence decrease, we hypothesized that the turbidity increase induced during Mg_2_P_2_O_7_ crystal growth results in light scattering and fluorescence attenuation, in agreement with previous reports. Moreover, to exclude the possibility that the GSH-AuNCs inhibit the crystal growth process, we obtained SEM images at different time intervals of a representative reaction (12 mM Mg^2+^ with 3 mM P_2_O_7_^4-^), in the presence of GSH-AuNCs. This experiment proved the progressive Mg_2_P_2_O_7_ crystal size increase (**Fig. S6**), consistent with crystal formation theory.^56–58^

**Figure 2:**
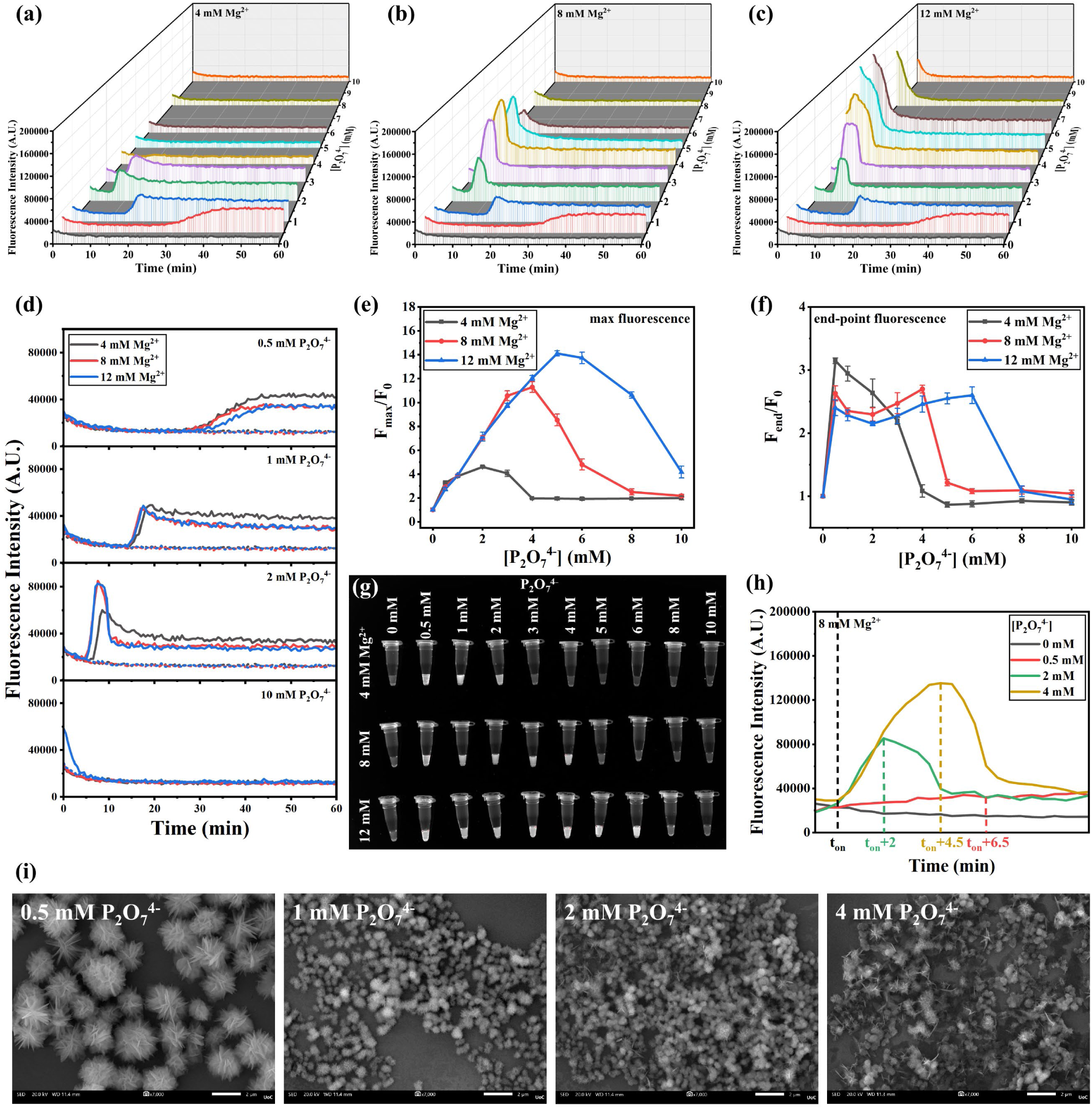
(a-c) Real-time fluorescence “waterfall-type” spectra of GSH-AuNCs recorded during Mg_2_P_2_O_7_ formation, in the presence of 4, 8 or 12 mM Mg^2+^ together with 0-10 mM P_2_O_7_^4-^. (d) Comparison of four representative real-time fluorescence graphs including the incubation of 4, 8 or 12 mM Mg^2+^ with 0.5, 1, 2 and 10 mM P_2_O_7_^4-^; dotted lines represent the fluorescence in the absence of P_2_O_7_^4-^ (0 mM). Relative fluorescence intensity of the (e) maximum fluorescence (F_max_/F_0_) and (f) end-point fluorescence (F_end_/F_0_) against [P_2_O_7_^4-^] of all real-time reactions, where F_0_ is the background fluorescence at 0 mM P_2_O_7_^4-^. (g) Photograph of all solutions tested in (a-d) captured under the GelDoc Go Imaging System. (h) Comparison of four fluorescence curves for 8 mM Mg^2+^, plotted so that the t_on_ values are aligned (total time shown on the x-axis: 12 min). (i) SEM images of the produced aggregates formed after incubating 8 mM Mg^2+^ with 0.5, 1, 2 and 4 mM P_2_O_7_^4-^ in the presence of 1 μL GSH-AuNCs (scale bar: 2 μm). The individual large, round-shaped nanoflakes observed in the presence of 0.5 mM P_2_O_7_^4-^ become smaller crystals in the presence of 1-4 mM P_2_O_7_^4-^.

#### Effect of [Mg^2+^] and [P_2_O_7_^4-^] on the crystallization process

Based on the above rationale, *i.e.*, that fluorescence enhancement recorded up to F_max_ may be a measure of the first stages of the crystal formation in solution, we attributed the differences observed during the interaction of various [Mg^2+^] and [P_2_O_7_^4-^] (**Fig. 2a-c**) to concentration-induced variations of this process. In a close-up observation of the curves obtained within a short window (12 min) from the beginning of the reaction (**Fig. 2h**), we observe that, in the case of *8* mM Mg^2+^ incubated with 4 mM P_2_O_7_^4-^, F_max_ (assumed to reflect the nucleation phase) is reached ∼4.5 min after t_on_. Moreover, the signal transitions sharply to a lower level, consistent with the beginning of growth of crystals and light scattering. In the presence of 2 mM P_2_O_7_^4-^, F_max_ is reached earlier (∼2 min after t_on_) probably due to the formation of fewer precursors/nuclei giving a lower signal than before. In this case, the transition from nucleation to crystal growth and induced fluorescence decrease is more gradual. Finally, when 0.5 mM P_2_O_7_^4-^ is incubated with 8 mM Mg^2+^, no optical transition is recorded (no “hat-shape” peak) and F_max_=F_end_ is reached 6.5 min after the onset of the reaction. This observation may support a slower pre-seeding and nucleation rate than before, leading inevitably to fewer nucleation points growing gradually without a clear separation of nucleation from growth, in agreement with previous observations.^58^

#### Crystal size at F_end_ and proposed GSH-AuNCs/Mg_2_P_2_O_7_ co-aggregation mechanism

The size of the crystals at the end of the reaction was also measured, following reports that Mg_2_P_2_O_7_ crystals exhibit different morphologies/sizes when formed under increasing [P_2_O_7_^4-^].^8^ Other studies also showed that the crystallization process plays an important role in determining the crystal structure and morphology.^13,16^ When we acquired SEM images of the GSH-AuNCs/Mg_2_P_2_O_7_ aggregates produced by incubating various [P_2_O_7_^4-^] with 8 mM Mg^2+^, a correlation between crystal size and [P_2_O_7_^4-^] was also revealed: large crystals (2846.5±131.9 nm) are formed at 0.5 mM P_2_O_7_^4-^, whereas significantly smaller ones were produced at 1, 2 and 4 mM P_2_O_7_^4-^ (651.2±26.7, 546.3±22.6 and 580.7±25.0 nm, respectively) (**Fig. 2i** and **Fig. S7a**).

Taking the SEM images together with the absorption and fluorescence measurements, we propose the following interpretation of the observed real-time kinetics of the GSH-AuNCs/Mg_2_P_2_O_7_ interaction. As suggested before, we associate the initial fluorescence increase in real-time graphs with the beginning and progression of the nucleation of Mg_2_P_2_O_7_. In the case where the [P_2_O_7_^4-^] is low with respect to the available [Mg^2+^], this may result in the formation of few nucleation points leading to large aggregates and later crystals (in the ∼3 μm range) which do not significantly affect fluorescence measurement, hence the signal stabilization (**Fig. 2d**, 0.5 mM P_2_O_7_^4-^). However, in the presence of higher [P_2_O_7_^4-^], the formation of a larger number of nucleation points probably favors the growth of more crystals of smaller size (∼600 nm), leading to light scattering that affects fluorescence enhancement. Overall, it appears that the observed 2-phase (“hat-shaped”) response represents a combined effect of the concentration-dependent fluorescence increase (nucleation points) and subsequent decrease (crystal formation) related to the size and number of GSH-AuNCs/Mg_2_P_2_O_7_ complexes, as well as the solution turbidity. A schematic of the interaction mechanism is presented in **Scheme 2**, where the two processes, *i.e.*, nucleation and crystal formation, appear to take place primarily before and after the observed F_max_. To assess the potential effect of temperature on the observed responses, we repeated selected experiments without heating (**Fig. S8a, b**). Our results indicate that similar response curves are obtained, although the various fluorescence signal phases occur at later times and with different F_end_ values, owing to slower reaction kinetics in the absence of heating.^58^

**Scheme 2:**
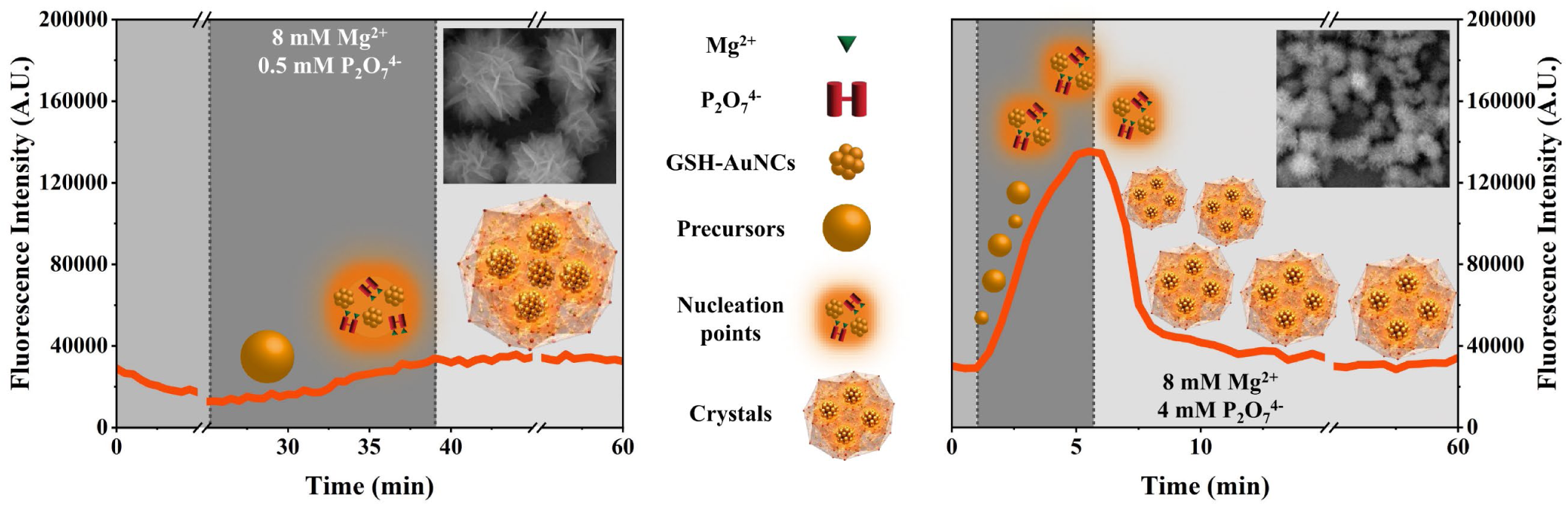
Illustration of the proposed mechanism of fluorescence emission enhancement of GSH-AuNCs induced by co-aggregation with Mg_2_P_2_O_7_. The dark grey area represents the initial stages of the Mg_2_P_2_O_7_ crystal-formation process that leads to nucleation (drawings are not to scale).

#### Effect of [GSH-AuNCs]

As a last parameter, we considered the effect of the amount of GSH-AuNCs on the co-aggregation and crystallization process. Increasing the GSH-AuNCs amount to 8 μL/reaction (**Fig. S9a-e)** provided a F_end_/F_0_ trend (**Fig. S9f**) that aligns closely with the trend of the Mg_2_P_2_O_7_ solid formation as a function of the [P_2_O_7_^4-^] based on absorbance measurements (**Fig. 1f**), although the signals are still lower than the corresponding F_max_. This result suggests that the saturation of F_end_ we observed before (**Fig. 2f**) is also due to the limited amount of available GSH-AuNCs (1 μL/reaction). Overall, the convergence of F_end_ signals to similar values in **Fig. 2f** for a certain [P_2_O_7_^4-^] range can be attributed to several parameters including, in addition to the number of GSH-AuNCs entrapped per crystal, the amount and size of formed crystals and induced light scattering. As a last note, it should be mentioned that the GSH-AuNCs amount also affects the Mg_2_P_2_O_7_ crystal formation especially at low [P_2_O_7_^4-^]. In the case of 0.5 mM P_2_O_7_^4-^, increasing amounts of GSH-AuNCs result in smaller crystals, while in the case of 2 and 4 mM P_2_O_7_^4-^, Mg_2_P_2_O_7_ crystals of similar size were produced (**Fig. S10a, b**). This result reveals that the GSH-AuNCs probe at higher concentrations may be an active component in the reaction, which can perturb the system under specific experimental conditions.

#### Theoretical calculations

The above conclusions are further strengthened by simulated calculations of the theoretically predicted values of solid [Mg_2_P_2_O_7_], free [Mg^2+^], free [P_2_O_7_^4-^], [MgP_2_O_7_^2-^], and saturation (S_initial_ and S_final_) profiles using Eq. (1) and (2) (**Fig. S11a-f**). These simulations illustrate the solid Mg_2_P_2_O_7_ formation, the redistribution of Mg^2+^ into soluble MgP_2_O_7_^2-^ complexes at excess P_2_O_7_^4-^, and the resulting decrease in supersaturation/precipitation tendency. Comparison of the simulations with our experimentally derived fluorescence measurements shows that the F_max_/F_0_ trend (**Fig. 2e**) aligns closely with the theoretically calculated amount of solid Mg_2_P_2_O_7_ at equilibrium. However, the subsequent fluorescence decrease toward F_end_ implies a transient optical state reflecting early nucleation/formation and AIE confinement before redistribution of GSH-AuNCs to the growing crystals. Therefore, S_initial_, which is the initial/no solid supersaturation ratio, is a better fit for qualitative correlation with the F_max_/F_0_ trend. Finally, the F_end_/F_0_ trend should match the solid Mg_2_P_2_O_7_ at equilibrium, provided there is an adequate amount of GSH-AuNCs present in the solution (**Fig. S9f**), as well as similarly sized crystals across the different experimental conditions (*i.e.*, qualitative correlation may not hold at low [P_2_O_7_^4-^] due to size differences affecting fluorescence).

### Fluorescence enhancement of GSH-AuNCs/Mg_2_P_2_O_7_ in the presence of additives encountered in genetic analysis

The formation of Mg_2_P_2_O_7_ from the interaction of the two inorganic reagents in the presence of GSH-AuNCs provides a model system to explore increasingly complex reactions approaching conditions that closely mimic a real enzymatic amplification reaction. Encouraged by the above results, we tested the fluorescence enhancement of GSH-AuNCs co-aggregation with Mg_2_P_2_O_7_ in the presence of components typically encountered in an enzymatic amplification reaction mix, such as those used for LAMP.^53^ We used 8 mM Mg^2+^ with 1 mM P_2_O_7_^4-^ for Mg_2_P_2_O_7_ formation, while modifying the concentration of relevant components in a LAMP-compatible mM reaction range (simulated LAMP reactions).

#### Interference of DTT in GSH-AuNCs/Mg_2_P_2_O_7_ co-aggregation

We first tested the effect of dithiothreitol (DTT), a small molecule used as a reducing agent for protein disulfide bonds and a common reagent in molecular biology buffers (typical concentration 0.04 mM in a LAMP reaction) for maintaining enzyme activity over time.^59^ Since DTT contains two terminal thiol (-SH) groups in its reduced form, we considered its potential interaction with the Au core of the GSH-AuNCs.^60^ According to **Fig. 3a**, the presence of DTT does reduce the overall fluorescence signal, possibly due to the bridging of the GSH-AuNCs with DTT, resulting in insufficient free GSH-AuNCs for co-aggregation with Mg_2_P_2_O_7_ and fluorescence enhancement.^60^ Increasing the amount of GSH-AuNCs in the solution from 1 to 3 μL partly restored the fluorescence enhancement during Mg_2_P_2_O_7_ production (**Fig. 3b**), indicating that excess GSH-AuNCs can both saturate reduced DTT and remain available to interact with Mg_2_P_2_O_7_. Moreover, no significant differences were observed, either at the time point at which the fluorescence signal increased, or in the “hat-shaped” effect; only the intensity changed, indicating a lower number of fluorescent GSH-AuNCs present in the solution when DTT is added. Notably, the oxidized form of DTT, which exists as a stable, six-membered ring with a disulfide bond (-S-S-) formed by the internal oxidation of its two -SH groups (**Fig. 3a**, inset), may not directly interact with the AuNC core, thereby influencing their stability. Additionally, DTT may induce thiol-disulfide exchange with oxidized GSH in the form of glutathione disulfide if present in the solution, potentially altering the redox balance and further destabilizing the AuNCs.^61^

#### Effect of (NH_4_)_2_SO_4_ on Mg_2_P_2_O_7_ formation

Ammonium sulfate (NH_4_)_2_SO_4_ is a common component of isothermal amplification buffers, which enhances primer stringency and enzyme performance by modulating ionic strength and stabilizing nucleic acid duplex formation.^52^ The salt dissociates into ammonium (NH_4_^+^) and sulfate (SO_4_^2-^) ions, where NH_4_^+^ help reduce nonspecific primer annealing and SO_4_^2-^ contribute to ionic strength and enzyme stabilization through Hofmeister-type effects that influence hydration shells and intermolecular interactions.^62^ Although moderate ionic strength improves amplification specificity, elevated sulfate concentrations can perturb precipitation equilibria and colloidal stability in NCs-based systems.^9^ To evaluate the compatibility of (NH_4_)_2_SO_4_ with the GSH-AuNCs/Mg_2_P_2_O_7_ aggregates, we examined concentrations ranging from 0 to 100 mM, exceeding the ∼10 mM typically used in amplification reactions. At 0-10 mM, no significant changes were observed in t_on_, fluorescence intensity or the characteristic “hat-shaped” profile, indicating that ionic strengths within this range do not disturb Mg_2_P_2_O_7_ formation or GSH-AuNCs co-aggregation (**Fig. 3c**). This observation is consistent with studies on fluorescent AuNCs demonstrating that low to moderate ionic strength produces negligible changes in fluorescence intensity, indicating good resistance to ionic interference.^63^ In contrast, at higher [(NH_4_)_2_SO_4_] (50-100 mM), the characteristic “hat-shaped” fluorescence curve observed in the presence of 1 mM P_2_O_7_^4-^ was no longer evident. Instead, fluorescence increased at a delayed time (t_on_ appearing ∼10 min later than before) and stabilized at a higher intensity. SEM analysis (**Fig. 3d**) revealed size-dependent changes in Mg_2_P_2_O_7_ crystal morphology, with higher (NH_4_)_2_SO_4_ levels producing larger aggregates compared to the 0 mM (NH_4_)_2_SO_4_ (**Fig. 2i**, 1 mM P_2_O_7_^4-^). The average crystal sizes increased from 840.9±26.3 nm at 5 mM and 1067.8±54.3 nm at 10 mM to 4498.7±225.1 nm at 50 mM and 6127.3±223.9 nm at 100 mM (**Fig. S7b**). The formation of larger Mg_2_P_2_O_7_ aggregates at high [(NH_4_)_2_SO_4_] suggests an influence on the nucleation and growth kinetics of Mg_2_P_2_O_7_. This effect may arise from combined electrostatic screening, sulfate-specific hydration effects,^64^ and changes in the effective activities of Mg^2+^ and P_2_O_7_^4-^. The gradual increase of fluorescence and final stabilization at F_max_∼F_end_ is consistent with the formation of larger crystals as discussed in **Fig. 2i**.

#### Effect of dNTPs on Mg_2_P_2_O_7_ formation during fluorescence monitoring via GSH-AuNCs

Nucleoside triphosphates (NTPs or dNTPs) are important building blocks in DNA/RNA synthesis and, for this reason, indispensable components in a nucleic acid amplification mix. Previous studies have shown that nucleic acids can be incorporated inside Mg_2_P_2_O_7_ crystals and co-precipitate to form monodisperse composites^8^ and/or micro/nano particles.^9^ Therefore, we tested the effect of dNTPs on the fluorescence of GSH-AuNCs/Mg_2_P_2_O_7_ complexes, during the interaction of different dNTP concentrations (3.2, 5.6 and 8 mM) (**Fig. 3e**). As shown, the addition of dNTPs at a high concentration (8 mM) completely suppressed fluorescence enhancement. This result led us to assume either that the two negatively charged entities compete, leading to GSH-AuNC displacement, or that a high dNTP concentration inhibits or alters Mg_2_P_2_O_7_ formation by sequestering free Mg^2+^. The latter is consistent with previous studies showing that dNTPs bind^2^ to Mg^2+^ in a nearly 1:1 ratio.^65^ At lower [dNTPs], clear fluorescence enhancement was observed at a delayed time, likely reflecting sufficient free Mg^2+^ available to interact with 1 mM P_2_O_7_^4-^. Notably, the distinct “hat-shaped” response observed was smoothed/abolished in the presence of 3.2 and 5.6 mM dNTPs, respectively, with fluorescence instead stabilizing at a higher fluorescence intensity (F_end_) compared to the absence of dNTPs (0 mM). SEM analysis showed the formation of crystal sizes of 799.6±30.8 nm (3.2 mM dNTPs), 1270.3±102.8 nm (5.6 mM dNTPs) and 2938.4±157.7 nm (8 mM dNTPs), all larger than those formed at 0 mM dNTPs (651.2±26.7) (**Fig. 3f** and **Fig. S7c**), consistent with previous fluorescence signal stabilization after F_max_ in the presence of larger Mg_2_P_2_O_7_ crystals. The production of very large crystals (∼ 3 μm) in the presence of 8 mM dNTPs which exhibit no fluorescence supports the hypothesis that the ∼1:1 [free Mg^2+^]/[P_2_O_7_^4-^] ratio favors the production of the soluble MgP_2_O_7_^2-^, leaving GSH-AuNCs free in the solution. This is also consistent with the production of a small number of large crystals, as indicated by the low turbidity measurement of the solution or the concentration of GSH-AuNCs/Mg_2_P_2_O_7_ aggregates (**Fig. S12**). The overall behavior described above at 1 mM P_2_O_7_^4-^ is maintained at higher [P_2_O_7_^4-^] (see **Fig. S13a-d**).

**Figure 3:**
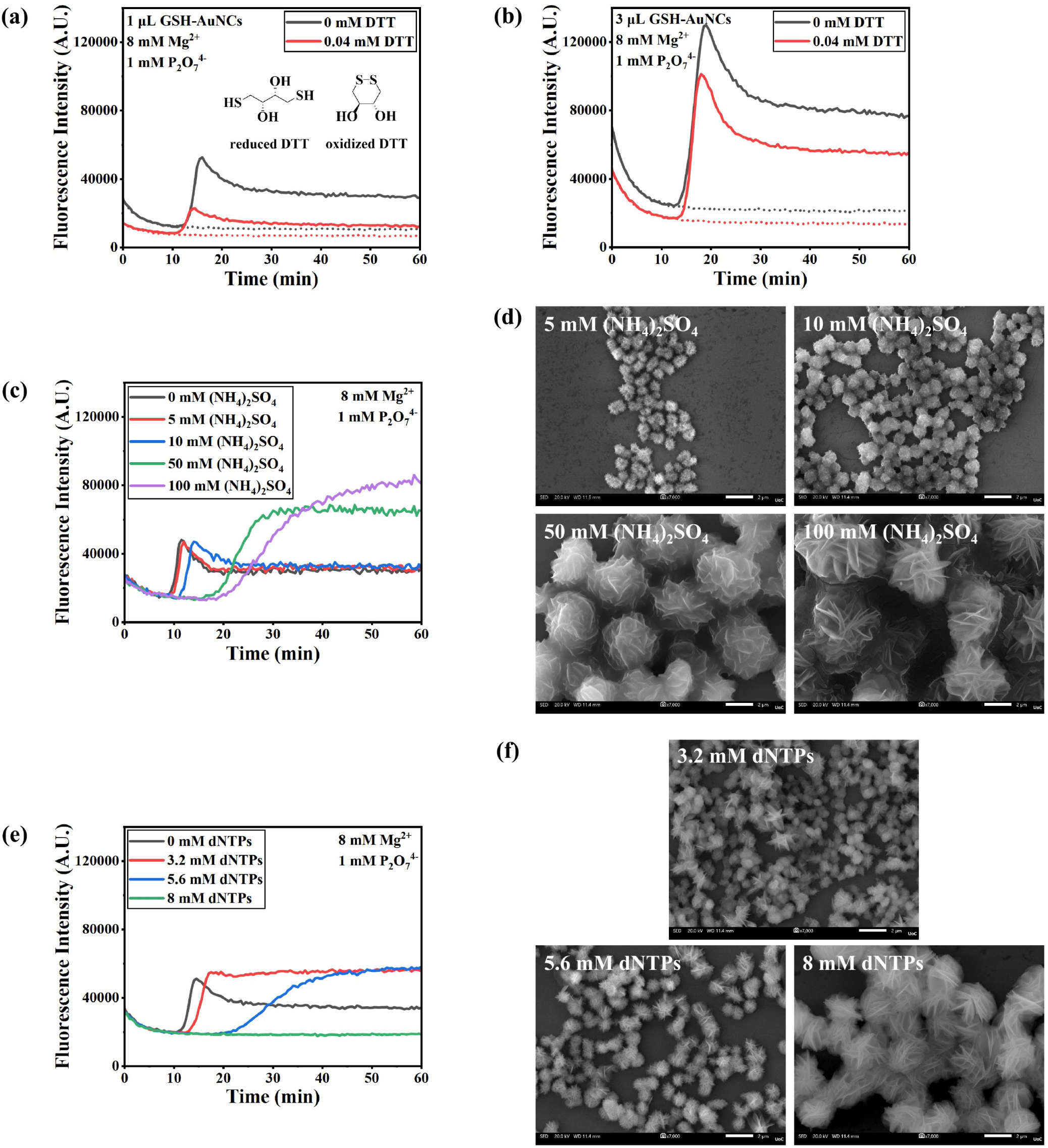
(a, b) Real-time fluorescence monitoring of Mg_2_P_2_O_7_ formation in the presence of 0.04 mM reduced DTT and 1 μL or 3 μL GSH-AuNCs. Control reactions with 0 mM DTT and P_2_O_7_^4-^ are also shown (dotted lines), while chemical structures of reduced and oxidized DTT are presented in the inset of (a). (c) Effect of [(NH_4_)_2_SO_4_] (5, 10, 50 and 100 mM) on the fluorescence response of GSH-AuNCs (1 μL) during the incubation of 8 mM Mg^2+^ with 1 mM P_2_O_7_^4-^. (d) Representative SEM images of Mg_2_P_2_O_7_ structures formed in the absence and presence of [(NH_4_)_2_SO_4_] (scale bar: 2 μm). (e) Effect of [dNTPs] (3.2, 5.6 and 8 mM) on the fluorescence response of GSH-AuNCs (1 μL) during the incubation of 8 mM Mg^2+^ with 1 mM P_2_O_7_^4-^. (f) Representative SEM images of Mg_2_P_2_O_7_ structures formed in the absence and presence of [dNTPs] (scale bar: 2 μm).

### Mechanistic studies of GSH-AuNCs/Mg_2_P_2_O_7_ co-aggregation guide the design of optimized LAMP

Results on the formation of Mg_2_P_2_O_7_ from the two inorganic species provide valuable insight for the design of an efficient amplification reaction. Specifically, the [Mg^2+^]/[P_2_O_7_^4-^] ratio emerges as a useful parameter for defining experimental conditions that yield enhanced Mg_2_P_2_O_7_ production in a genetic analysis assay such as LAMP, including both end-point and real-time detection. For qualitative detection, a visible color change *(e.g.,* under UV illumination) occurs at the end of a simulated reaction (*i.e.*, absence of Bst and other LAMP reagents) when this ratio exceeds 2 **(Fig. 4a and Fig. 2f, g**). In real enzymatic amplification assays (presence of Bst), typical concentrations of dNTPs range from 2-6 mM (0.5-1.5 mM of each dNTP). Assuming 100% amplification efficiency, such that all dNTPs are enzymatically converted to P_2_O_7_^4-^, the maximum dNTP concentration of 6 mM in the presence of 4 and 8 mM Mg^2+^ would give ratios of 0.7 and 1.3, respectively, *i.e.*, below the threshold ratio of 2 for a visible end-point color. This observation suggests that only 12 mM Mg^2+^ would enable naked-eye detection at high dNTP concentrations. In practice, amplification efficiency is typically below 100%, implying the ability to also use the 8 mM Mg^2+^ with 5 or 6 mM dNTPs in a qualitative assay. **Fig. 4b** further shows that the inhibitory effect of DTT (0.04 mM included in a reaction using 8 U Bst) on the co-aggregation of GSH-AuNCs with Mg_2_P_2_O_7_ can be minimized by using an excess of GSH-AuNCs (3 μL instead of 1 μL), enabling end-point fluorescence detection by eye under UV irradiation in a standard DTT-containing amplification mix.

For quantitative genetic analysis of amplified nucleic acids, the key parameter is the time (t_on_) at which fluorescence begins to increase, typically referred to in a diagnostic assay as time-to-positivity (TTP). TTP decreases together with the [Mg^2+^]/[P_2_O_7_^4-^] ratio, reflecting faster Mg_2_P_2_O_7_ formation. **Fig. 4c-(i)** provides a representative set of reference reactions regarding the Mg_2_P_2_O_7_ formation using an excess of GSH-AuNCs, focusing on a 10-minute window around the onset of the fluorescence. Onset values observed in **Fig. 4c-(i)** are very short due to the immediate availability of P_2_O_7_^4-^. However, during an amplification reaction, dNTPs are converted to P_2_O_7_^4-^ enzymatically (essentially, dNTPs are precursors of P_2_O_7_^4-^), which, in turn, interact with the Mg^2+^ to form Mg_2_P_2_O_7_. When we monitored the fluorescence of a GSH-AuNCs-containing reverse transcription LAMP (RT-LAMP) reaction in the presence of Bst (2 U) and 10^5^ Influenza A RNA copies, TTP values were 40.5 and 27.5 min in the presence of 3.2 and 5.6 mM of dNTPs, respectively (**Fig. 4c-(ii)**), *i.e.*, the [dNTPs] that produce the higher amount of P_2_O_7_^4-^ (5.6 mM) results in a shorter TTP. To improve the efficiency of the reaction, we also tested a higher [Bst] (4 and 8 U). As expected, lower TTP values were obtained as the Bst concentration increased. When 8 U Bst was used with 3.2 mM dNTPs, the TTP decreased by 5 min, and the reaction produced a fluorescence profile approaching that of the reference reaction with a [Mg^2+^]/[P_2_O_7_^4-^] ratio near 2.5. Repeating the above experiment using 5.6 mM dNTPs resulted in a TTP that was even shorter, *i.e.*, it occurred 6 min earlier in the presence of 8 U Bst and exhibited a more pronounced “hat-shaped’’ curve (**Fig. 4c-(iii)**).

Based on the above, we conclude that using a low [Bst] (2 U) with a high [dNTPs] (5.6 mM) accelerates the response compared with low [dNTPs] (3.2 mM), but may result in partial dNTPs incorporation into Mg_2_P_2_O_7_, potentially inhibiting amplification.^9,15^ This is reflected in the stabilized fluorescence signal, similar to **Fig. 3e**, suggesting formation of larger Mg_2_P_2_O_7_ crystals incorporating dNTPs due to slower conversion to P_2_O_7_^4-^. In contrast, combining a high [Bst] (8 U) with a high [dNTPs] (5.6 mM) markedly improves amplification efficiency and response time while minimizing interference. Accordingly, we selected 5.6 mM dNTPs with either 4 or 8 U Bst for further evaluation studies, as both conditions yield close TTPs, with the lower Bst volume offering a cost advantage. Finally, it is noted that when we used a higher [dNTPs] (8 mM), it was found to inhibit the formation and/or fluorescence enhancement of the GSH-AuNCs/Mg_2_P_2_O_7_, in agreement with other studies^2,14^ and our previous results (**Fig. 3e**). It should be noted that in a real LAMP reaction, formation of larger crystals and fluorescence stabilization after co-aggregation-induced fluorescence enhancement using GSH-AuNCs can occur due to dNTPs and (NH_4_)_2_SO_4_ acting synergistically, depending on the LAMP isothermal buffer used. A more detailed analysis of the combined influence of dNTPs and Bst concentrations on amplification kinetics and efficiency is presented in **Fig. S14**.

**Figure 4:**
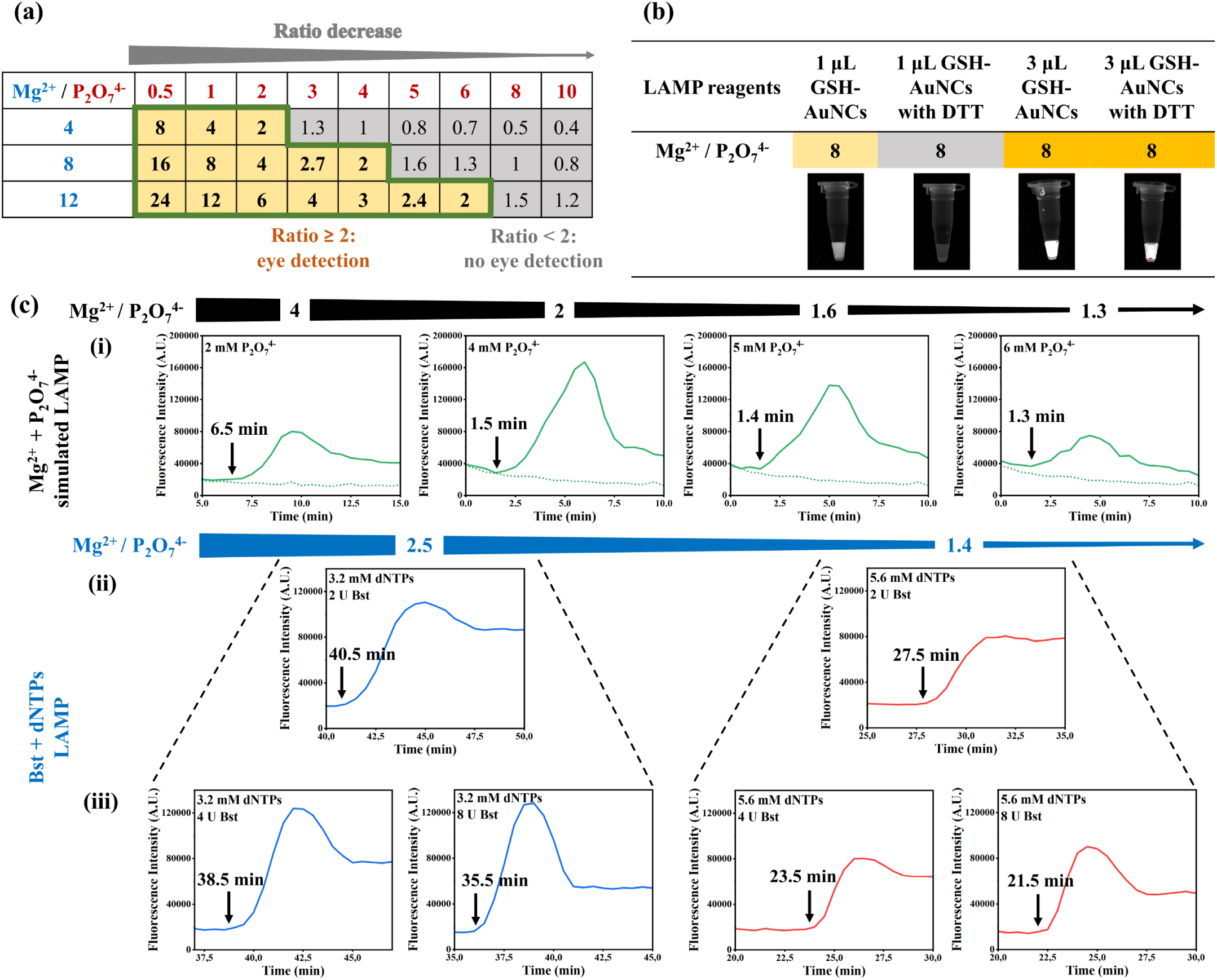
(a) Matrix showing the effect of [Mg^2+^]/[P_2_O_7_^4-^] ratio on the fluorescence of simulated LAMP reactions at end-point, upon incubation of various [Mg^2+^] and [P_2_O_7_^4-^] in the presence of 1 μL of GSH-AuNCs; yellow and grey cells indicate enhanced and weak fluorescence, respectively. (b) Photos of simulated LAMP reactions produced in the presence (0.04 mM) or absence of DTT, combined with 1 or 3 μL GSH-AuNCs. (c) Comparison of real-time fluorescence responses of various [Mg^2+^]/[P_2_O_7_^4-^] ratios during (i) simulated LAMP reactions; (ii) real LAMP reactions, using 3.2 or 5.6 mM dNTPs with 2 U Bst, and (iii) same dNTPs as in (ii) with 4 or 8 U of Bst. In both simulated and real LAMP reactions, 8 mM Mg^2+^ was used. In all the above graphs, the time-to-positivity (TTP), depicted with an arrow, indicates the point at which the Mg_2_P_2_O_7_ production starts, while for the real LAMP reactions, 10^5^ Influenza A RNA copies were used.

### Real-time LAMP detection of Influenza A RNA using GSH-AuNCs

As a final validation, we assessed the sensitivity of GSH-AuNCs as a fluorescent probe during a one-pot real-time LAMP (**Scheme 3**). For these experiments, we used two concentrations of Bst polymerase (4 and 8 U) during simultaneous reverse transcription and isothermal amplification (RT-LAMP) of 10-fold serial dilutions of target RNA (Influenza A). As shown in **Fig. 5a**, when 4 U of Bst was used, the 10^6^ and 10^2^ copies/reaction were detected at 20.2±0.4 and 21.8±0.6 min, respectively. Increasing Bst concentration to 8 U accelerated detection, yielding responses at 16.8±0.2 min for 10^6^ copies and 19.3±0.3 min for 10^2^ copies (**Fig. 5b**).

**Scheme 3:**
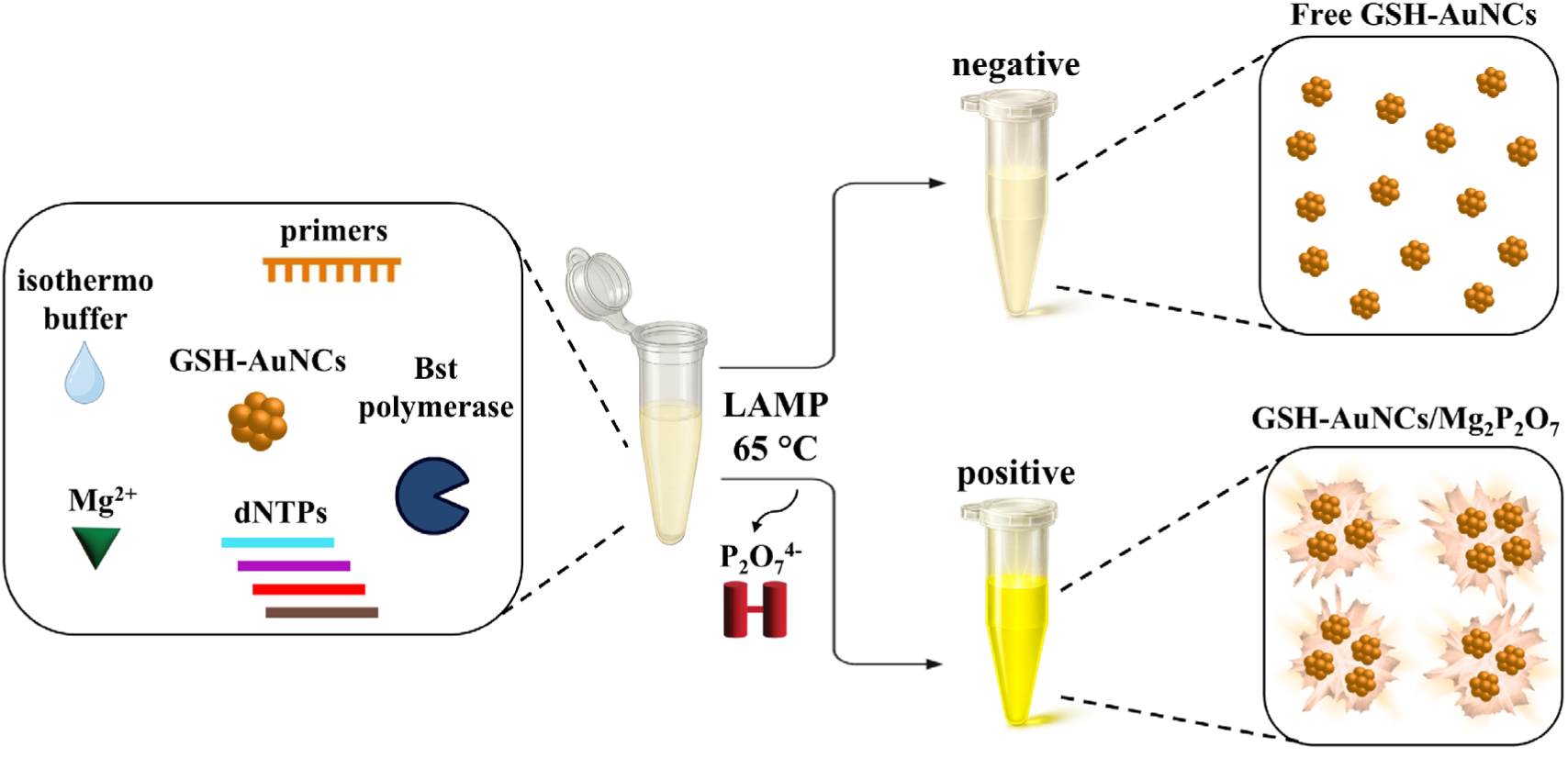
Representation of the developed LAMP assay for real-time detection using GSH-AuNCs and fluorescence enhancement measurement upon co-aggregation with Mg_2_P_2_O_7_.

The relationship between RNA copy number and TTP values is shown in **Fig. 5c** for triplicate measurements for each concentration. A strong linear correlation (R^2^ = 0.99) was observed using 4 U Bst across 6 orders of magnitude (10^2^ to 10^8^ RNA copies/reaction), while a coefficient of R^2^ = 0.92 across 4 orders of magnitude (10^2^ to 10^6^ RNA copies/reaction) was obtained when 8 U Bst was used. Notably, the real-time amplification curves (**Fig. 5a and b**) deviate from the characteristic sigmoidal shape of a conventional LAMP reaction, instead exhibiting a subtle “hat-shaped” profile. This behavior is consistent with our reference reactions, though less pronounced than predicted under conditions containing 5-6 mM P_2_O_7_^4-^ (in 8 mM Mg^2+^). This result suggests amplification efficiency below 100%, potentially with partial interference to the resulting Mg_2_P_2_O_7_ aggregates by other components in the reaction (*e.g.*, excess dNTPs, (NH_4_)_2_SO_4_, or accumulated DNA amplicons). To demonstrate the potential of our method outside a laboratory setting, we performed the GSH-AuNCs-coupled RT-LAMP using simulated crude samples, including 10^2^ and 10^3^ Influenza A RNA copies spiked into 20% lysed saliva. As shown in **Fig. 5d**, both concentrations were successfully amplified and detected using the GSH-AuNCs probe, with a slightly slower TTP in the presence of crude samples. Further comparison with a real-time colorimetric method showed that our newly developed GSH-AuNCs assay is comparable to the commonly used HNB colorimetric metal indicator, when the same assay was run on the Omega plate reader in absorbance mode (650 nm) (**Fig. S15a, c**). When we ran the same assay using HNB in the commercially available Pebble device,^66^ a significant improvement in TTP was observed, probably attributed to the more efficient heating at 65 °C (**Fig. S15b, c**).

The proposed new quantitative LAMP monitoring approach, which relies on GSH-AuNCs and detection of Mg_2_P_2_O_7_ crystals/aggregates, represents a broadly applicable and target-independent detection strategy. In this respect, it is similar to real-time SYBR Green I fluorescent detection, although the latter produces a signal upon intercalation with dsDNA. Both approaches can be affected by nonspecific amplification through primer-dimer formation; therefore, the method does not overcome the limitation of detecting a LAMP by-product and relies on the use of four to six primers to substantially reduce nonspecific amplification. However, it offers the means to design a fully optimized assay not necessarily bound to commercial kits and amplification buffer solutions. Importantly, detailed analysis of the characteristic “hat-shaped” fluorescence profile confirmed that this feature does not interfere with TTP determination, enabling accurate quantitative detection. This contrasts with prior reports describing SYBR Green I-based real-time LAMP, where such effects compromised monitoring ability.^67^

**Figure 5:**
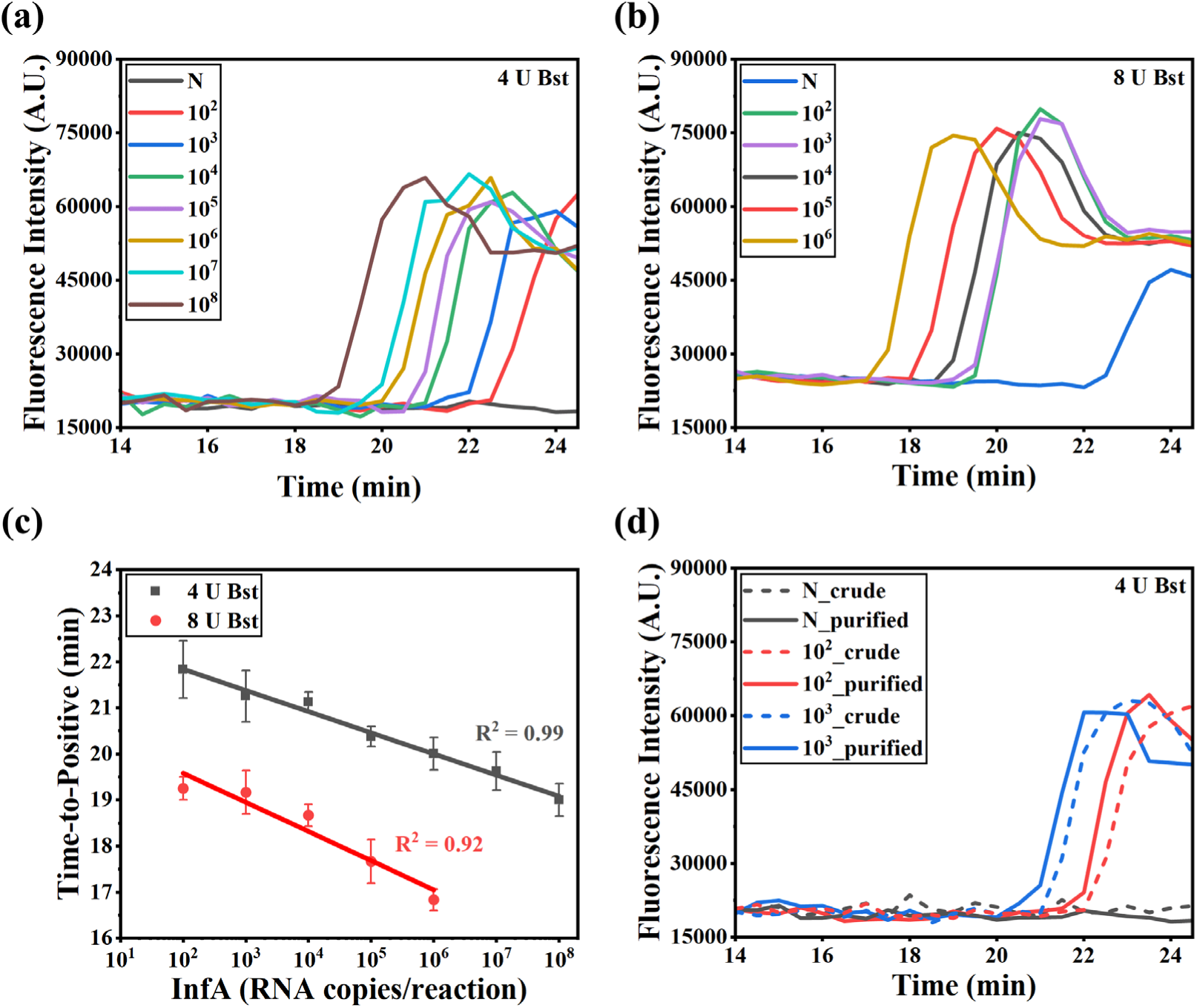
(a, b) Real-time fluorescence signals obtained in Omega Plate Reader using 3 μL GSH-AuNCs and 5.6 mM dNTPs. The 25 μL reactions contained 10^2^ to 10^8^ Influenza A RNA copies/reaction for (a) 4 U Bst and 10^2^ to 10^6^ Influenza A RNA copies/reaction for (b) 8 U Bst. Note that the nonspecific response (primer-dimer) observed in (b) at 22.2 min is well above the TTP for the 10^2^ RNA copies/reaction. (c) Calibration curves derived from the above real-time fluorescence measurements, revealing a linear trend over 6 and 4 orders of magnitude for 4 and 8 U Bst, respectively, during the detection of Influenza A RNA. (d) Real-time fluorescence signals obtained in the presence of 20% crude saliva, revealing comparable TTP to reactions with purified RNA at 10^2^ and 10^3^ Influenza A RNA copies per reaction, using 4 U Bst.

## Conclusions

In this work, we synthesized GSH-AuNCs and showed that in the presence of Mg_2_P_2_O_7_ crystals they co-aggregate, leading to fluorescence enhancement. This discovery allowed the design and testing of a previously unreported class of nanoparticle-based fluorescent probes, enabling for the first time the real-time fluorescent monitoring of Mg_2_P_2_O_7_ formation using NCs. A good correlation between the fluorescence signal and crystal formation theory in solution allowed us to attribute the GSH-AuNCs/Mg_2_P_2_O_7_ co-aggregation response to the crystal formation and growth during the interaction of Mg^2+^ with P_2_O_7_^4-^ as a function of the [Mg^2+^]/[P_2_O_7_^4-^] ratio. Importantly, the real-time fluorescence signal appeared to be a good indicator of the size of the formed crystals, with the presence or absence of a “hat-shaped” response correlated with the formation of smaller (<1.1 μm) or larger (from 1.3 to 6.1 μm) crystals, respectively. The above results were further confirmed when the Mg_2_P_2_O_7_ formation occurred in the presence of GSH-AuNCs and DTT, (NH_4_)_2_SO_4_, dNTPs or Bst, all additives encountered in a LAMP reaction. Guided by the onset of the reaction and kinetics of Mg_2_P_2_O_7_ formation, an optimized LAMP assay was developed and demonstrated for the detection of 100 copies/reaction of Influenza A RNA in less than 25 min, even in the presence of crude specimens. Our RT-LAMP outperformed other QD- and NC-based assays that relied on qualitative, end-point analyses, where the nanostructures were introduced after amplification.^25,26,68^ While preloading the probes inside the reaction tube-cap avoided contamination, such handling is less practical than the single-step integration achieved with our method, which simplifies operation while maintaining quantitative accuracy. Our findings support the use of GSH-AuNC fluorescent probes as a general strategy for developing customized LAMP assays under diverse enzymatic and reaction conditions, thereby supporting the advancement of diagnostics through nanotechnology.^69^ Beyond genetic analysis, we believe that the presented methodology can pave the way for Mg_2_P_2_O_7_ monitoring in biotechnological applications of high significance. Our approach enables the use of GSH-AuNCs for real-time monitoring of Mg_2_P_2_O_7_ formation, opening the possibility for the study of the nucleation and crystallization processes of Mg_2_P_2_O_7_ in the presence of key components such as RNA and DNA. In addition, during IVT, the sequestration of DNA due to crystal formation and degradation/hydrolysis of P_2_O_7_^4-^ to two phosphate ions (PO_4_^3-^) by inorganic pyrophosphatase (PPase) are important processes that could potentially be investigated using the reported methodology with profound implications for IVT engineering.^2^ Finally, the Mg_2_P_2_O_7_-induced fluorescence enhancement represents a broadly applicable platform where more types of similar nanoclusters could be used, expanding its potential impact across diverse bioanalytical applications.

## Materials and Methods

### Reagents and materials

Gold (III) chloride trihydrate (HAuCl_4_ x 3H_2_O, ACS reagent, ≥49% Au basis), l-glutathione reduced (GSH, ≥ 98%), sodium pyrophosphate decahydrate (Na_4_P_2_O_7_ x 10H_2_O, ACS reagent, ≥99%), magnesium sulfate (MgSO_4_) solution (for molecular biology, 1±0.04 M), ammonium sulfate ((NH_4_)_2_SO_4_, for molecular biology, ≥99%) and water for chromatography (LC-MS grade) LiChrosolv^®^ were purchased from Merck (Darmstadt, Germany). Tris buffer (1 M liquid, pH 8.0, molecular biology grade) was purchased from PanReac AppliChem; 2-Hydroxy-1-(2-hydroxy-4-sulfo-1-naphthylazo)-3,6-naphthalenedisulfonic acid, trisodium salt (HNB) was purchased from Dojindo Laboratories; and Dithiothreitol (DTT, 0.1 M) was purchased from Thermo Fisher Scientific/Invitrogen. All chemicals were used as received, without any further purification. For the preparation of LAMP reactions, solutions of 10x Isothermo Buffer (Mg^2+^ free), Bst DNA/RNA Polymerase (8 U/μL), Mg^2+^ (100 mM) and dNTPs mix (dATP, dCTP, dGTP, dTTP of 10 mM each) were purchased from SBS Genetech Co., Ltd. (China). For the preparation of crude sample-containing reactions, normal saliva (pooled human donors) was purchased from Lee Biosolutions (USA). The set of six primers (100 μM) for Influenza A was purchased from Metabion (Germany), where their sequences (5’-3’) were as follows:

Inner FIP: CCAAATGCAATGGGGCTACCATCTTCTGGAAGACAAGCA
Inner BIP: TAACATTGCTGGCTGGATCCTACAATGTAGGACCATGATCT
Outer F3: AACAGTAACACACTCTGTCA
Outer B3: CATTGTCTGAATTAGATGTTTCC
Loop F: CCTCTTAGTTTGCATAGTTTTCCGT Loop B: CCAGAGTGTGAATCACTCTCCAC
Finally, synthesized Influenza A RNA target (25 μg) in nuclease-free water was purchased from Synbio Tech (USA), where the DNA sequence used was as follows: GTAGACACAGTACTAGAAAAGAATGTAACAGTAACACACTCTGTCAATCTTCTGGAAGACAAGCATAACGGAAAACTATGCAAACTAAGAGGGGTAGCCCCATTGCATTTGGGTAAATGTAACATTGCTGGCTGGATCCTGGGAAATCCAGAGTGTGAATCACTCTCCACAGCAAGATCATGGTCCTACATTGTGGAAACATCTAATTCAGACAATGGAACGTGTTACCCAGGAGATTTCATCAATTATGAGGAGCTAAGAGAGCAATTGAGCTCAGTGTCATCATTTGAAAGGTTTGAAATATTCCCCAAGACAAGTTCATGGCCTAATCATGACTCGGACAAAGGTGTAACGGCAGCATGTCCTCACGCTGGAGCAAAAAGCTTCTACAAAAACTTGATATGGCTGGTTAAAAAAGGAAATTCATACCCAAAGCTCAACCAAACCTACAT

### GSH-AuNCs synthesis

GSH-AuNCs were synthesized as described before,^45^ with a few modifications. Briefly, 5 mL of HAuCl_4_ (4 mM) was added to an equal volume of GSH (6 mM), and the solution was stirred for ∼15 min. Subsequently, the temperature was raised to 90 °C, and the 10 mL solution remained under vigorous stirring/heating for ∼5 h. Afterwards, the heating plate was turned off, and the solution was left under stirring until it reached room temperature (25 °C). Finally, the resulting bright yellow GSH-AuNCs solution was stored at 4 °C until further use. Taking into account the specific parameters of our system, *i.e.*, a mean diameter (D) of 2.2±0.4 nm extracted *via* TEM images, and a final reaction volume V=10 mL, a molar concentration of 6.3 μM was calculated for the GSH-AuNCs dispersion.

### Neat GSH-AuNCs & GSH-AuNCs/Mg_2_P_2_O_7_ characterization

#### Spectroscopy measurements

UV-Visible (UV-Vis) absorption spectroscopy (Nanodrop One Spectrophotometer, ThermoFisher, USA) was utilized for optical characterization of GSH-AuNCs. Measurements were performed using 2 μL samples that were placed on a pedestal, while the data were automatically normalized to 10 mm path length equivalent. As a reference, the absorbance of the Tris buffer (20 mM, pH 8) was measured. Fluorescence spectroscopy (FS) measurements were carried out to investigate the emission spectra of the synthesized GSH-AuNCs using a fluorescence spectrofluorometer (Horiba Jobin Yvon, Fluoromax 3, Japan) with excitation from a CW Xenon arc lamp and the following settings: slit excitation at 2 nm, slit emission at 5 nm, t=0.5 s, and excitation wavelength of 370 nm. In this case, samples were measured in a quartz glass cuvette (10 mm optical path length) containing 2 mL sample volumes. For both UV-Vis and FS measurements, the synthesized GSH-AuNCs solutions were diluted 25-fold in Tris buffer (20 mM, pH 8). Attenuated total reflectance/Fourier transform infrared spectroscopy (ATR-FTIR) was employed to obtain the IR spectrum of the neat GSH and GSH-AuNCs over a spectral region of 4000 to 400 cm^-1^ using the VERTEX 70v FT-IR Spectrometer (Bruker, Germany), equipped with a Platinum ATR unit (A225/Q) with a single reflection diamond crystal. To achieve a high signal quality, transmission spectra were collected at a resolution of 4 cm^-1^ and in 8 replicate scans. GSH was measured as a powder, while GSH-AuNCs solution (undiluted) was drop-cast onto a chemically cleaned glass substrate and left to dry at 25 °C. FTIR spectra were also obtained for neat Mg_2_P_2_O_7_ and GSH-AuNCs/Mg_2_P_2_O_7_ complexes, after spin-down (∼30 s) to collect the supernatant, drying (25 °C) and measurement of the resulting powders. The neat Mg_2_P_2_O_7_ and GSH-AuNCs/Mg_2_P_2_O_7_ powders were also investigated *via* X-ray diffraction (XRD) spectroscopy, using a Bruker AXS D8 Advance copper anode diffractometer. XRD patterns were collected over a 2*θ* range of 10° to 40° with a scan rate of 0.02° s^−1^, using a monochromatic Cu K*α* radiation source (*λ* = 1.54056 Å). Elemental analysis was also performed using an energy-dispersive X-ray spectroscopy (EDX) module (DrySD60 detector), mounted on an SM-IT700HR (JEOL Ltd., Japan) microscope. In this case, the solutions were vortexed, 20 μL were then deposited onto chemically cleaned SiO_2_ substrates *via* drop-casting, and finally dried (25 °C) prior to measurement. To ensure adequate sensitivity for GSH-AuNCs detection, an excess of GSH-AuNCs was added to preformed GSH-AuNCs/Mg_2_P_2_O_7_ solutions, prior to final sample preparation. Notably, all complexes investigated using the above techniques were formed using 8 mM Mg^2+^ and 4 mM P_2_O_7_^4-^.

#### Microscopy & imaging

Transmission electron microscopy (TEM) (JEM-2100, JEOL Ltd., Japan) operating at an accelerating voltage of 200 kV (Mag-150 k) was used to obtain images for morphological characterization of the GSH-AuNCs. Specimens for TEM measurements were prepared by drop-casting 10 μL of diluted GSH-AuNCs (1:25 with LC-MS grade water) onto carbon-coated copper TEM grids, and left to dry at 25 °C. Image analysis of the TEM micrographs was performed using the ImageJ software. To calculate the diameter of the synthesized GSH-AuNCs, a total of 250 AuNCs were measured from different areas of the grid. Scanning electron microscopy (SEM) was employed to characterize the morphology of Mg_2_P_2_O_7_ crystals. A JSM-IT700HR (JEOL Ltd., Japan) microscope, operated at an accelerating voltage of 20 kV, was employed to characterize the morphology of crystals produced upon incubation of P_2_O_7_^4-^ (0.5, 1, 2 or 4 mM) with Mg^2+^ (8 mM) in the presence or absence of either (NH_2_)_2_SO_4_ (5, 10, 50 and 100 mM), or dNTPs (3.2, 5.6 and 8 mM), all in the presence of 1 μL GSH-AuNCs. Additionally, a JEOL 7000F microscope (JEOL Ltd., Japan), operated at an accelerating voltage of 15 kV, was further employed to study the effect of the GSH-AuNCs (0, 1 and 3 μL) on the formed Mg_2_P_2_O_7_ crystals (25 μL total volume reactions). Both instruments revealed Mg_2_P_2_O_7_ crystals of similar size, when the same amount of GSH-AuNCs (1 μL) was present in the solution throughout the tested range. Using the JEOL 7000F microscope, SEM images of Mg_2_P_2_O_7_ crystals formed at different time intervals (1, 3, 5, 7, 9, 11 and 13 min) were also obtained. For sample preparation, 20 μL of the final mixtures upon reaction completion and vortexing to redisperse the produced Mg_2_P_2_O_7_ precipitate were deposited onto chemically cleaned SiO_2_ substrates *via* drop-casting, and dried (25 °C) prior to imaging, similar to EDX measurements. Regarding the samples with Mg_2_P_2_O_7_ crystals formed at different time intervals, the drying was performed under vacuum, to minimize continued crystal growth or Ostwald ripening during sample preparation and to ensure that the imaged crystal population more closely reflects the size distribution present at the time of collection. Analysis of the SEM images was performed using the ImageJ software, and used to measure the diameters of the formed crystals. Confocal microscopy (Leica TCS SP8 laser scanning microscope, Leica Microsystems, Germany) was used to acquire images from the fluorescent GSH-AuNCs/Mg_2_P_2_O_7_ complexes, where GSH-AuNCs (3 μL per 25 μL reaction) were incubated either *in situ* with the reagents during Mg_2_P_2_O_7_ formation, or added afterwards to preformed Mg_2_P_2_O_7_ crystals (formed using 8 mM Mg^2+^ and 4 mM P_2_O_7_^4-^). Solutions were drop-cast onto chemically cleaned glass substrates, dried (65 °C), and the resulting dried samples were covered using 0.13-0.17 mm thick cover glass slips. Similar measurement and analysis conditions were ensured for both samples, *i.e.*, excitation laser at 405 nm (3% intensity), emission window of 498-720 nm (177.8 gain, −0.01 offset), pinhole 95.6 μm, scan speed at 400 Hz and 0.3 μm pixel size (z dimension). Representative slices of the samples were chosen and compared, while the image analysis was performed in the visible (brightfield/transmitted) and in the fluorescence (red) channels. For both samples, Mg_2_P_2_O_7_ was formed using 8 mM Mg^2+^ and 4 mM P_2_O_7_^4-^.

#### Zeta potential measurements

Z-potential values of the studied solutions were determined using a Malvern Zetasizer Nano ZS90 (Malvern Instruments Ltd, UK), after diluting GSH-AuNCs (1:25) in Tris buffer (20 mM), pH 8, or in LC-MS grade water at pH 4.

### Preparation and evaluation of simulated LAMP reactions

#### End-point fluorescence measurements

Simulated LAMP reactions were prepared upon incubation of MgSO_4_ (8 mM) with various P_2_O_7_^4-^ concentrations (0-10 mM) in Tris (20 mM), pH 8, in the presence of GSH-AuNCs (25-fold diluted) in a final volume of 1 mL. Upon heating at 65 °C for 60 min in a thermoshaker (MSC-100, Labgene Scientific, Switzerland), the solutions were vortexed to disperse the Mg_2_P_2_O_7_ pellet and their fluorescence was measured as described before (Fluoromax 3). To remove the turbidity effect induced by Mg_2_P_2_O_7_ from the fluorescence emission curves, similarly prepared solutions using the same [Mg^2+^]/[P_2_O_7_^4-^] ratios but without GSH-AuNCs were measured, and their corresponding spectra were subtracted from the fluorescence spectra. Photographs of these solutions at the end of the reaction were taken with a standard smartphone upon illumination with a UV lamp (Herolab UV transilluminator UVT-28 MP, Germany). End-point evaluation of similarly prepared solutions (using 4 and 12 mM Mg^2+^ in addition to 8 mM Mg^2+^) was carried out in 0.2 mL PCR tubes (25 μL total reaction volume). Heating was performed at 65 °C (SimpliAmp Thermal Cycler, ThermoFisher Scientific, USA), while fluorescence was qualitatively evaluated using a GelDoc Go Gel Imaging System (Bio-Rad, USA). The end-point fluorescence reading was done under the “Nucleic Acid Gels” application category (UView option).

#### Real-time monitoring

For the real-time measurements, the FLUOstar Omega filter-based multi-mode microplate reader (BMG Labtech, Germany) was used, with excitation and emission filters at 370 nm and 610 nm, respectively, and with the capability for heating up to 65 °C. Reactions (25 μL) comprising 0-10 mM P_2_O_7_^4-^ together with 4, 8 or 12 mM Mg^2+^ and 1 μL of as-synthesized GSH-AuNCs were set up in 96-well half-area polystyrene microplates (μCLEAR^®^ *black,* Greiner Bio-One, Germany), upon addition of 25 μL of mineral oil on the top of each well, followed by sealing of the microplate with a sealer (ampliseal, Greiner Bio-One, Germany) to avoid evaporation. Initially, the instrument was preheated to 65 °C for 3 h; a total of 120 cycles (30 s cycle time with 40 flashes) was set for each experiment, while fluorescence measurements were performed using the bottom optic reading option. Moreover, a gain adjustment (90%) was performed using a well containing 25 μL of undiluted GSH-AuNCs. The F_end_/F_0_ ratio of GSH-AuNCs/Mg_2_P_2_O_7_ complexes extracted from real-time measurements (Omega plate reader) showed a ∼2.5-fold increase, while the end-point fluorescence measurements using Fluoromax 3 showed a ∼4.7-fold increase, although in both cases the GSH-AuNCs used were 25-fold diluted. This difference may arise from the different instruments and experimental conditions used, such as subtraction of turbidity spectra in the Fluoromax 3 measurements, different temperature conditions during F_end_ extraction, and different sample volumes (25 μL in Omega, compared to 2 mL in Fluoromax). Furthermore, to test the effect of temperature on Mg_2_P_2_O_7_ formation in the presence of GSH-AuNCs, we repeated the above experiment by setting the microplate reader at 25 °C. Similar real-time measurements using the above setup were also performed during the incubation of Mg^2+^ (8 mM) with: (i) DTT (0.04 mM), P_2_O_7_^4-^ (1 mM) with either 1 or 3 μL GSH-AuNCs, (ii) varying amounts of (NH_4_)_2_SO_4_ (5, 10, 50 and 100 mM), P_2_O_7_^4-^ (1 mM) with GSH-AuNCs (1 μL), and (iii) varying amounts of dNTPs (3.2, 5.6 and 8 mM) and P_2_O_7_^4-^ (1-5 mM) with GSH-AuNCs (1 μL), all in Tris (20 mM, pH 8).

#### Turbidity estimation

Absorbance at 610 nm was used as a turbidity indicator at the end of the reactions (65 °C incubation for 60 min). The 25 μL reactions contained 8 mM Mg^2+^, 0-10 mM P_2_O_7_^4-^ and 1 μL GSH-AuNCs, or 8 mM Mg^2+^, 1 mM P_2_O_7_^4-^, 1 μL GSH-AuNCs and varying amounts of dNTPs (3.2, 5.6 and 8 mM), all in Tris (20 mM, pH 8). The measurements were performed in triplicate as described before (Nanodrop One).

### Preparation and evaluation of LAMP reactions

LAMP reactions (65 °C) were set up in 25 μL total volume containing 2.5 μL isothermo buffer, 2 μL Mg^2+^ (from 100 mM stock), 1 μL Bst DNA/RNA polymerase (8 U/μL), 3.5 μL dNTPs (from stock containing dATP, dCTP, dGTP, dTTP, 10 mM each), 2.5 μL of PM (containing 18 μM FIP and BIP, 2 μM F3 and B3 and 6 μM Loop-F and Loop-B), 3 μL of GSH-AuNCs fluorescent probe, 9.5 μL of nuclease-free water and 1 μL of the target, here Influenza A RNA; in the case of the negative control, the 1 μL of the target was replaced by nuclease-free water. For the calibration curve, a starting concentration of 10^8^ copies/reaction of Influenza A was used upon serial dilutions down to 10^2^ copies/reaction. For the crude saliva samples, 5 μL of lysed saliva (95 °C for 10 min) was added together with 4.5 μL of nuclease-free water, in the LAMP mix described above. To prevent solvent evaporation during heating, mineral oil (15-25 μL) was added over the LAMP mix. To study the effect of varying Bst DNA/RNA polymerase (0.25, 0.5 and 1 μL corresponding to 2, 4 and 8 U/reaction) and dNTPs amounts (2, 3.5 and 5 μL corresponding to 3.2, 5.6 and 8 mM/reaction) on the fluorescence signal, nuclease-free water volumes were appropriately adjusted to keep a total reaction volume of 25 μL. LAMP reactions containing the commercially available colorimetric HNB dye (120 μM final concentration) instead of 3 μL of GSH-AuNCs were also used to derive a calibration curve of Influenza A RNA within the range of 10^2^-10^6^ copies/reaction, using the Omega plate reader (absorbance reading at 650 nm) and a portable real-time colorimetric device for LAMP (Pebble, Biopix DNA Technology, Greece). For the heating and evaluation of the fluorescence in real LAMP reactions similar procedures were followed, as described above for the simulated LAMP reactions.

### Software

Plotting and statistical analyses were performed using Origin (OriginLab Corporation, USA). Standard deviations (SD) were calculated using the StdDevP function, which computes the population standard deviation of a specified database. Data are presented as mean±SD, while all experiments were performed in triplicate, unless otherwise stated. Image analysis for determination of Mg_2_P_2_O_7_ crystal sizes was performed using ImageJ software (National Institutes of Health, USA). SEM images were spatially calibrated to physical units using the embedded scale bar *via* the “Set Scale” function. Crystal size was determined by measuring the diameter of each crystal using the straight-line selection tool.

For each experimental condition, measurements were obtained from all individually resolvable crystals identified across at least three independent SEM images. Crystals overlapping with neighboring crystals or intersecting the image boundaries were excluded to avoid ambiguous or incomplete size measurements. Crystal sizes for each condition are reported as the mean±SD calculated from all individual crystals included in the analysis.

Equilibrium Mg^2+^ - P_2_O_7_^4-^ speciation and Mg_2_P_2_O_7_ precipitation trends were estimated using a custom Python script.

The calculations employed a simplified equilibrium model in which total [Mg^2+^] and [P_2_O_7_^4-^] were specified in mM, and aqueous speciation was described by the equilibria shown in Eq. (1). The mass balances were reduced to a single scalar equation in free Mg^2+^ and solved using Brent’s root-finding method. Mg_2_P_2_O_7(s)_ (solid) was expressed in Mg_2_P_2_O_7_ formula units, with each mole of solid consuming two Mg^2+^ equivalents and one P_2_O_7_^4-^ equivalent from the aqueous phase. Under supersaturated conditions, the equilibrium solid concentration was obtained by solving for the solid amount required to return the solution to Q=[Mg^2+^]^2^[P_2_O_7_^4-^]=K_sp_ (Eq. 2). The apparent concentration-form K_sp_ value for Mg_2_P_2_O_7_ ⋅ 3.5H_2_O (at 25 °C) and the thermodynamic parameters used to temperature-correct K_1_ and K_2_ to 65 °C using van’t Hoff relationships were taken from the literature.^16^ The model was used only as a reduced equilibrium trend calculation and did not include pH, P_2_O_7_^4-^ protonation, buffer interactions, or fluorescent probe-based binding effects. The Python script was developed with assistance from ChatGPT and manually reviewed, edited, and validated by the authors. Sourcecode, documentation, and software requirements are provided as Supplementary Data/Code.

## Supporting information

Supplementary Code (Python)

## Associated Content

### Supporting Information

Additional spectroscopic, electrokinetic, microscopic, diffraction, elemental, kinetic, simulated equilibrium modeling, additive-interference and LAMP-optimization data, as well as particle-size analyses, comparative LAMP colorimetric results and supporting references, can be found in the Supporting Information.

*Supplementary Figures*; **S1.** Spectroscopic, electrokinetic and morphological characterization of GSH-AuNCs. **S2.** ATR-FTIR, XRD and EDX measurements of neat Mg_2_P_2_O_7_ and GSH-AuNCs/Mg_2_P_2_O_7_ complexes. **S3.** Confocal microscopy images of GSH-AuNCs/Mg_2_P_2_O_7_ complexes, in which GSH-AuNCs were added either to preformed Mg_2_P_2_O_7_, or in situ with the Mg^2+^ and P_2_O_7_^4-^ reagents. **S4.** Real-time fluorescence monitoring of Mg_2_P_2_O_7_ formation under various [Mg^2+^]/[P_2_O_7_^4-^] ratios using GSH-AuNCs. **S5:** Relative fluorescence intensity of the maximum (F_max_/F_0_) and end-point fluorescence (F_end_/F_0_) signal against [Mg^2+^]/[P_2_O_7_^4-^] ratio. **S6:** SEM images of GSH-AuNCs/Mg_2_P_2_O_7_ complexes at different time intervals, formed using 12 mM Mg^2+^ and 3 mM P_2_O_7_^4-^. **S7:** Size of Mg_2_P_2_O_7_ crystals formed in the presence of GSH-AuNCs and various [P_2_O_7_^4-^], [(NH_4_)_2_SO_4_] or [dNTPs]. **S8.** Real-time fluorescence monitoring of Mg_2_P_2_O_7_ formation at 25 °C, using GSH-AuNCs. **S9:** Effect of different GSH-AuNCs amounts on the Mg_2_P_2_O_7_-induced fluorescence enhancement. **S10:** SEM images of Mg_2_P_2_O_7_ crystals formed in the absence or in the presence of 1 or 3 μL GSH-AuNCs. **S11:** Reduced equilibrium Mg^2+^-P_2_O_7_^4-^ speciation and supersaturation ratios (Python-simulated). **S12:** Turbidity measurements *via* absorbance (610 nm) of solutions containing a combination of 8 mM Mg^2+^, 1 mM P_2_O_7_^4-^ and various [dNTPs]. **S13:** Effect of [dNTPs] on the Mg_2_P_2_O_7_-induced fluorescence enhancement of GSH-AuNCs. **S14**: Real-time fluorescent monitoring of LAMP reactions using GSH-AuNCs, utilizing different [Bst] and [dNTPs] combinations. **S15:** Real-time quantitative colorimetric LAMP using HNB dye, utilizing either absorbance measurements (Omega plate reader) or image analysis (Pebble).

## Author Information

### Authors

**Stylianos Grammatikos** - Department of Biology, University of Crete, 70013 Voutes, Heraklion, Greece and Institute of Molecular Biology and Biotechnology, Foundation for Research and Technology-Hellas, 100 N. Plastira Str., 70013 Heraklion, Greece

**Konstantina Alexaki** - Institute of Molecular Biology and Biotechnology, Foundation for Research and Technology-Hellas, 100 N. Plastira Str., 70013 Heraklion, Greece

### Author Contributions (CRediT author statement)

1. **S. Grammatikos:** conceptualization, formal analysis, investigation, methodology, software, visualization, writing-original draft, writing-review & editing. **K. Alexaki:** data curation, formal analysis, investigation, methodology, visualization, writing-review & editing. **E. Gizeli:** investigation, methodology, project administration, supervision, validation, writing-review & editing.

All authors have given approval to the final version of the manuscript.

## Acknowledgments

The authors would like to acknowledge Prof. M. Stylianakis, Prof. D. Anglos, Dr. G. Kenanakis, Prof. S. Anastasiadis (IESL-FORTH) and Prof. C. Delidakis (IMBB-FORTH) for providing access to the ZP, FS, FTIR, XRD and Confocal instruments, respectively. Ms. K. Katsara, Dr. A. Philippidis, Ms. A. Manousaki (IESL-FORTH), Ms. S. Papadogiorgaki (Electron Microscopy Lab, University of Crete) and Dr. M. Stapountzi are also acknowledged for their assistance in taking FTIR measurements, FS measurements, SEM, TEM and Confocal images, respectively.

