## Supplementary Information File for "Mechanistic Insights into Magnesium Pyrophosphate Formation in the Presence of Gold Nanoclusters Enable Genetic Analysis via Co-Aggregation-Induced Fluorescence Enhancement"

### Supplementary Figures

**S1.** Spectroscopic, electrokinetic and morphological characterization of GSH-AuNCs

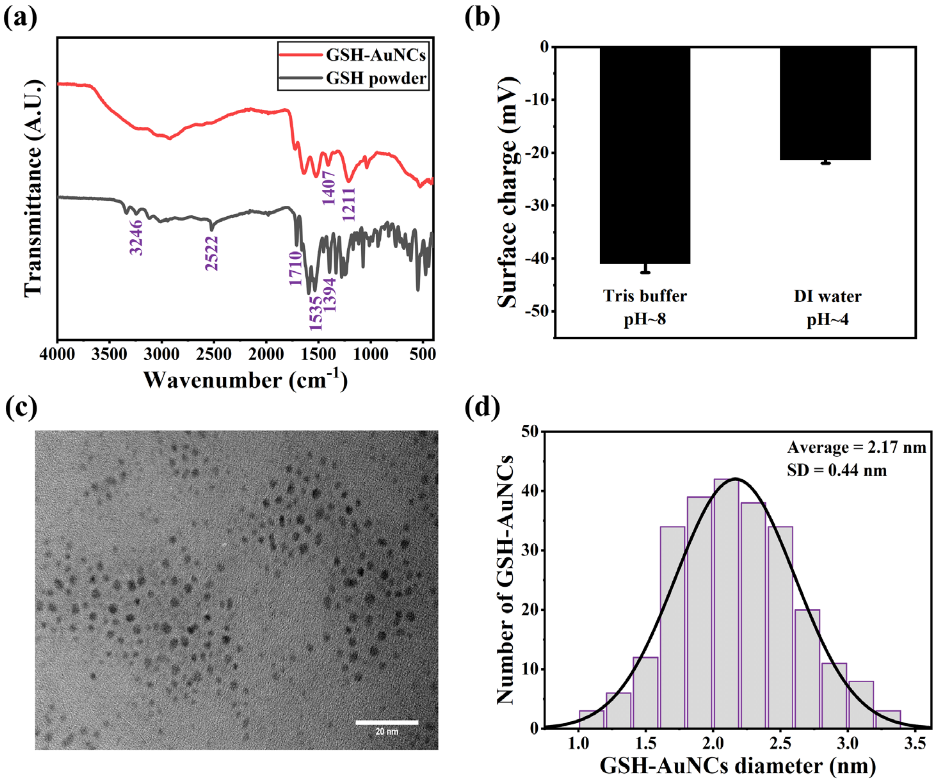

***Figure S1:*** *(a) ATR-FTIR spectra of GSH-AuNCs (dried film) and GSH (powder), revealing spectral changes due to GSH binding to the Au core. (b) Zeta potential values of GSH-AuNCs in the presence of either Tris buffer (pH 8) or LC-MS grade water (final pH~4). The GSH-AuNCs were 25-fold diluted in both solutions. (c) Representative TEM image of the spherical GSH-AuNCs (scale bar: 20 nm). (d) Size distribution histogram with a fitted distribution curve, obtained from measurements of approximately 250 GSH-AuNCs from different grid areas.*

Attenuated total reflectance-Fourier transform infrared (ATR-FTIR) spectroscopy provided further insights into the surface chemistry of the glutathione-capped gold nanoclusters (GSH-AuNCs). The spectrum of GSH exhibited characteristic vibrations, including a weak S-H stretch at 2522 $cm^{-1}$,[1] a sharp C=O stretch at 1710 $cm^{-1}$,[2] asymmetric and symmetric $-COO^-$ stretches at 1394 and 1535 $cm^{-1}$,[1] and a N-H vibration at 3246 $cm^{-1}$.[3] In the GSH-AuNCs spectrum, a significant decrease in the 2522 $cm^{-1}$ band implies thiol deprotonation and formation of an Au-S linkage.[3] Concurrently, the $-COO^-$ asymmetric stretch shifted to ~1211 $cm^{-1}$ and the symmetric stretch to ~1407 $cm^{-1}$, indicative of altered carboxylate electron density upon surface coordination (**Fig. S1a**).[1] These spectral changes demonstrate that GSH binds to the AuNCs through its thiolate sulfur, accompanied by deprotonation and reorganization of its carboxylate moieties.

Zeta potential (ZP) measurements performed in triplicate indicated that GSH-AuNCs in Tris buffer (pH 8) possess a highly negative surface charge (-41.0±1.7 mV), consistent with strong colloidal stability that minimizes aggregation through electrostatic repulsion. This behavior reflects the pH-dependent charge state of GSH, which is strongly negative at a pH above 7.[4] When dispersed in LC-MS grade water (final pH 4), the GSH-AuNCs exhibited a reduced ZP value of -21.3±0.7 mV, underscoring the pronounced influence of solution pH on the surface charge of AuNCs (**Fig. S1b**).

Transmission electron microscopy (TEM) images revealed that synthesized GSH-AuNCs have a spherical shape and a good overall uniformity (**Fig. S1c**), with a calculated mean diameter (D) of 2.2±0.4 nm (**Fig. S1d**). The solution concentration was calculated to be 6.3 µM, assuming that: (i) all gold ions ($Au^{3+}$) from $HAuCl_4$ were fully reduced to $Au^0$ atoms, (ii) the density (ρ) of each AuNC was taken as equal to that of bulk gold (19.3 $g/cm^3$), and (iii) the AuNCs are of spherical shape with a face-centered cubic (FCC) lattice.[5,6]

**S2.** ATR-FTIR, XRD and EDX measurements of neat $Mg_2P_2O_7$ and GSH-AuNCs/$Mg_2P_2O_7$ complexes

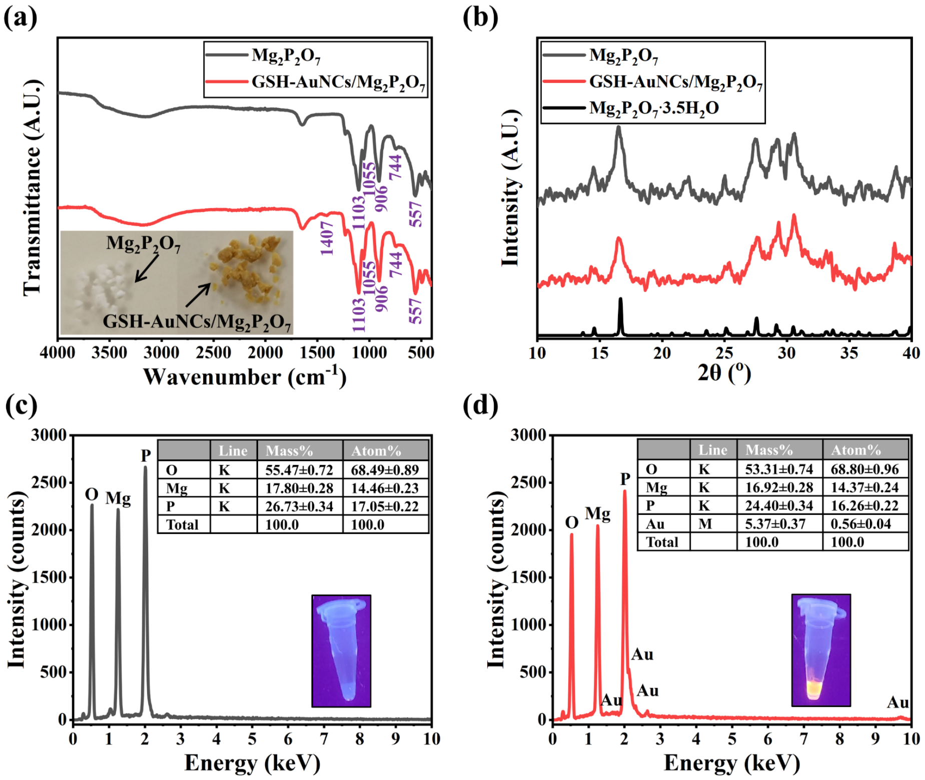

***Figure S2:*** *(a) ATR-FTIR spectra of neat $Mg_2P_2O_7$ and GSH-AuNCs/$Mg_2P_2O_7$ complexes. Inset: Photograph of the measured powders, highlighting the observed color difference. (b) XRD spectra of neat $Mg_2P_2O_7$ and GSH-AuNCs/$Mg_2P_2O_7$ complexes, along with the reference pattern of the $Mg_2P_2O_7 \cdot 3.5\ H_2O$ hydrate phase reported in the literature,[7] for comparison. (c, d) EDX analyses of neat $Mg_2P_2O_7$ and GSH-AuNCs/$Mg_2P_2O_7$ complexes, prepared using 8 mM $Mg^{2+}$ and 4 mM $P_2O_7^{4-}$. Inset: Photographs of the solutions under UV light. Note: to ensure good sensitivity in GSH-AuNCs detection, an excess of GSH-AuNCs was added after $Mg_2P_2O_7$ was formed using 8 mM $Mg^{2+}$, 4 mM $P_2O_7^{4-}$ and 1 μL GSH-AuNCs.*

ATR-FTIR spectroscopy (**Fig. S2a**) of the dried $Mg_2P_2O_7$ powders (neat and GSH-AuNC-containing) revealed all of the expected $P_2O_7^{4-}$ vibrations and, in the GSH-AuNCs-containing one, an additional band arising from the GSH ligand. More specifically, in both spectra, the terminal [$PO_3$] groups gave rise to an asymmetric stretching band at 1055 cm$^{-1}$ and a symmetric stretching band at 1103 cm$^{-1}$, the P-O-P bridge modes appeared at 906 cm$^{-1}$ (stretching) and 744 cm$^{-1}$ (bending), while the Mg-O lattice vibration was observed at 557 cm$^{-1}$.[8,9] Furthermore, both spectra exhibited a very weak, broad band between 3000 and 3720 cm$^{-1}$, attributable to O-H stretching vibrations of surface adsorbed hydroxyls or residual moisture.[9] Upon incorporation of GSH-AuNCs, the $P_2O_7^{4-}$ bands remained unchanged, and a new absorption band at 1407 cm$^{-1}$, assigned to the -COO$^-$ symmetric stretch, appeared, indicative of altered carboxylate electron density upon coordination to the Au surface.[1]

X-ray diffraction (XRD) spectroscopy was also employed to identify the crystalline phase of the $Mg_2P_2O_7$ precipitate (**Fig. S2b**). The Bragg reflections observed in the XRD patterns of both powders (neat and GSH-AuNC-containing) matched the reference pattern of the $Mg_2P_2O_7 \cdot 3.5H_2O$ hydrate phase as reported in the literature,[7] under the specific experimental conditions used. The GSH-AuNCs-containing powder did not present any other characteristic Au signals, implying the GSH-AuNCs quantum-sized dimensions, low content, uniform distribution and their physical and/or coordinative association within the $Mg_2P_2O_7$ crystals.[10]

The incorporation of GSH-AuNCs into the $Mg_2P_2O_7$ precipitate was further investigated by energy-dispersive X-ray spectroscopy (EDX) analysis of neat $Mg_2P_2O_7$ and GSH-AuNCs/$Mg_2P_2O_7$ complexes. Elemental analysis confirmed the presence of characteristic Mg, P, and O signals in both samples, while the presence of Au signals was additionally detected in the second complex, confirming the presence of GSH-AuNCs to the $Mg_2P_2O_7$ crystalline structures (**Fig. S2c, d**).

**S3.** Confocal microscopy images of GSH-AuNCs/$Mg_2P_2O_7$ complexes, in which GSH-AuNCs were added either to preformed $Mg_2P_2O_7$, or *in situ* with the $Mg^{2+}$ and $P_2O_7^{4-}$ reagents

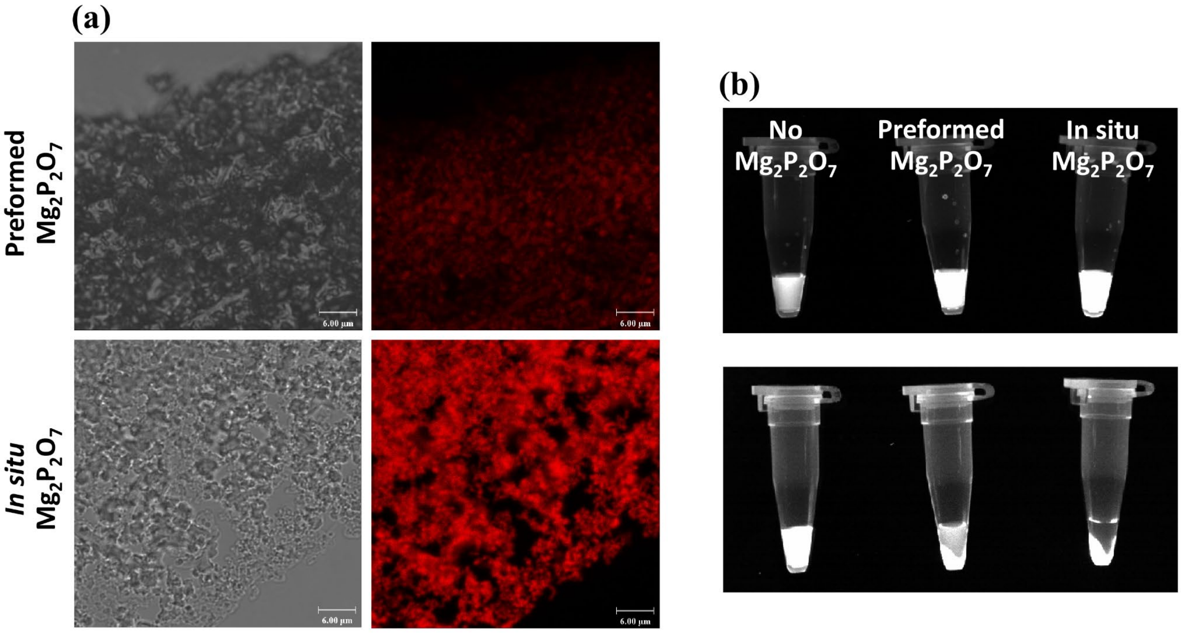

***Figure S3:*** *(a) Confocal images of GSH-AuNCs/$Mg_2P_2O_7$ complexes, comparing fluorescence of preformed (up) or in situ formed (down) $Mg_2P_2O_7$ crystals. Both the visible light channel (left; i.e., transmitted light/brightfield), and the fluorescence channel (right; i.e., red fluorescence) are depicted (scale bar: 6 μm). (b) Photographs of solutions containing GSH-AuNCs with no $Mg_2P_2O_7$ crystals (only $Mg^{2+}$) (left), preformed $Mg_2P_2O_7$ crystals (middle), or in situ formed $Mg_2P_2O_7$ crystals (right), before (up) or after (down) spin-down, under UV light. Note: in all cases, 3 μL GSH-AuNCs per 25 μL reaction were used, while $Mg_2P_2O_7$ crystals were formed using 8 mM $Mg^{2+}$ and 4 mM $P_2O_7^{4-}$ in Tris buffer (pH 8).*

Confocal microscopy imaging provided insights into the incorporation of GSH-AuNCs to the $Mg_2P_2O_7$ crystals. When they are formed in the presence of GSH-AuNCs, a higher fluorescence intensity can be observed, compared to when the same amount of GSH-AuNCs is added after $Mg_2P_2O_7$ is formed (**Fig. S3a**). In both cases, a fluorescence increase is observed when compared to the control ($Mg^{2+}$ only) under UV light (**Fig. S3b**). After spin-down, fluorescence is also apparent in the supernatant of the solution in the case of the preformed $Mg_2P_2O_7$, in contrast to the *in situ* formed $Mg_2P_2O_7$, indicating that more GSH-AuNCs are present in $Mg_2P_2O_7$ crystals in the latter. This implies that GSH-AuNCs may be embedded within *in situ* formed $Mg_2P_2O_7$ crystals, while they may be bound primarily to the surface of the preformed ones.[11]

**S4.** Real-time fluorescence monitoring of $Mg_2P_2O_7$ formation under various $[Mg^{2+}]/[P_2O_7^{4-}]$ ratios using GSH-AuNCs

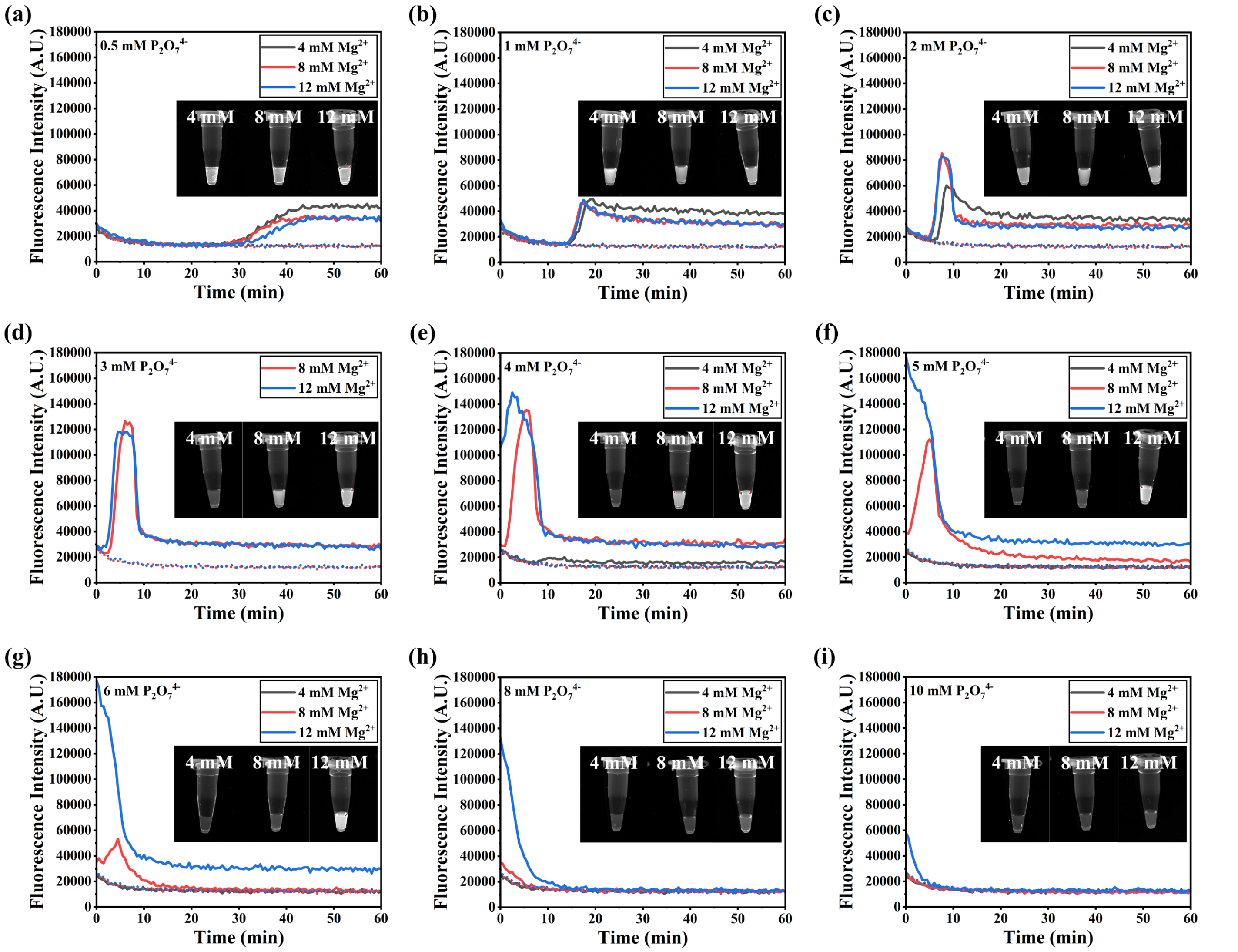

***Figure S4:*** *(a-i) Real-time fluorescence monitoring of $Mg_2P_2O_7$ formation using GSH-AuNCs (1 μL), in 25 μL reactions containing 4, 8 or 12 mM $Mg^{2+}$ and 0.5-10 mM $P_2O_7^{4-}$ (65 °C), with photographs of the solutions at the end of the reaction under UV light (insets). Note: dotted lines represent the fluorescence in the absence of $P_2O_7^{4-}$ (0 mM).*

Real-time fluorescence measurements were obtained during incubation of 4, 8 or 12 mM $Mg^{2+}$ with 0-10 mM $P_2O_7^{4-}$, in the presence of 1 μL GSH-AuNCs per 25 μL total reaction volume. Based on the results shown in **Fig. S4**, we observe: (i) increasing $[P_2O_7^{4-}]$ accelerates $Mg_2P_2O_7$ formation and shifts the fluorescence increase to earlier times, (ii) maximum fluorescence intensity when $[Mg^{2+}]/[P_2O_7^{4-}]$ ratio equals 2, (iii) "hat-shaped" kinetics for $[Mg^{2+}]/[P_2O_7^{4-}]>1$ (consisting of an initial fluorescence increase followed by a decrease and stabilization) and (iv) a signal overlapping with the 0 mM $P_2O_7^{4-}$ control when $[Mg^{2+}]/[P_2O_7^{4-}] <1$. Within the "hat-shaped" kinetics, when $[Mg^{2+}]/[P_2O_7^{4-}]\geq 2$, $Mg_2P_2O_7$ is expected to dominate, and fluorescence decreases and stabilizes at a value higher than the control (fluorescence enhancement). As the $[Mg^{2+}]/[P_2O_7^{4-}]$ ratio approaches 1, $MgP_2O_7^{2-}$ becomes predominant, and the final fluorescence decreases to a point where no fluorescence enhancement is observed (*i.e.*, release of GSH-AuNCs into the solution). Notably, in the case of 0.5 mM $P_2O_7^{4-}$, instead of the "hat-shaped" kinetics, a signal stabilization is observed, which could possibly be attributed to differences in the crystallization rate, as well as the size of final $Mg_2P_2O_7$ crystals, compared to those formed using higher $[P_2O_7^{4-}]$.

**S5.** Relative fluorescence intensity of the maximum ($F_{max}/F_0$) and end-point fluorescence ($F_{end}/F_0$) signal against $[Mg^{2+}]/[P_2O_7^{4-}]$ ratio

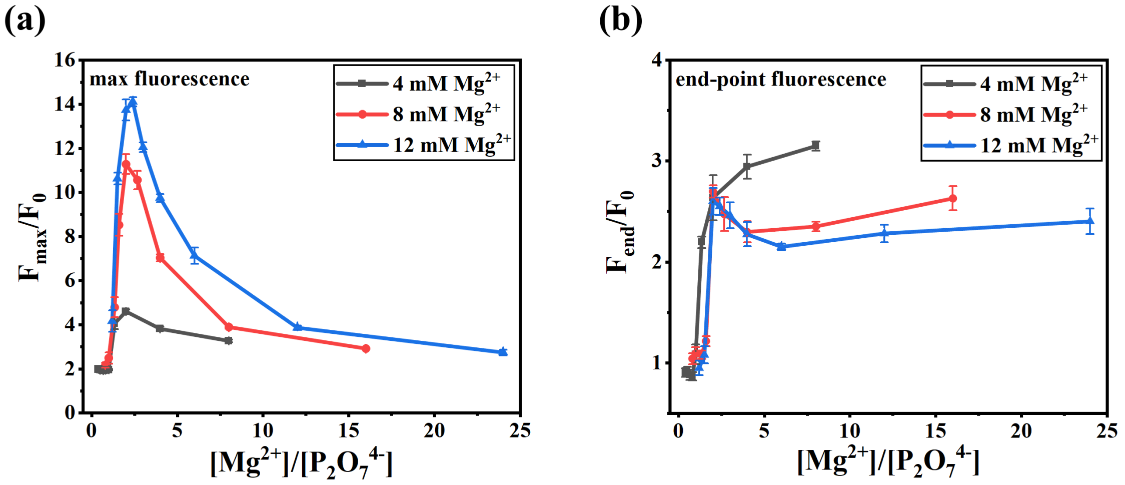

***Figure S5:*** *Relative fluorescence intensity of the (a) maximum ($F_{max}/F_0$) and (b) end-point fluorescence ($F_{end}/F_0$) against $[Mg^{2+}]/[P_2O_7^{4-}]$ ratio of the reactions containing 4, 8 or 12 mM $Mg^{2+}$, 0.5-10 mM $P_2O_7^{4-}$ and 1 µL GSH-AuNCs (65 °C).*

Based on **Fig. S5a**, it can be seen that the solutions exhibit maximum fluorescence when the $[Mg^{2+}]/[P_2O_7^{4-}]$ ratio is 2. However, end-point fluorescence values (**Fig. S5b**) do not follow a similar trend; instead, at high $[Mg^{2+}]/[P_2O_7^{4-}]$ ratios, *i.e.*, low $P_2O_7^{4-}$ compared to $Mg^{2+}$ concentrations, the signal increases, especially in the case of 4 mM $Mg^{2+}$.

**S6.** SEM images of GSH-AuNCs/$Mg_2P_2O_7$ complexes at different time intervals, formed using 12 mM $Mg^{2+}$ and 3 mM $P_2O_7^{4-}$

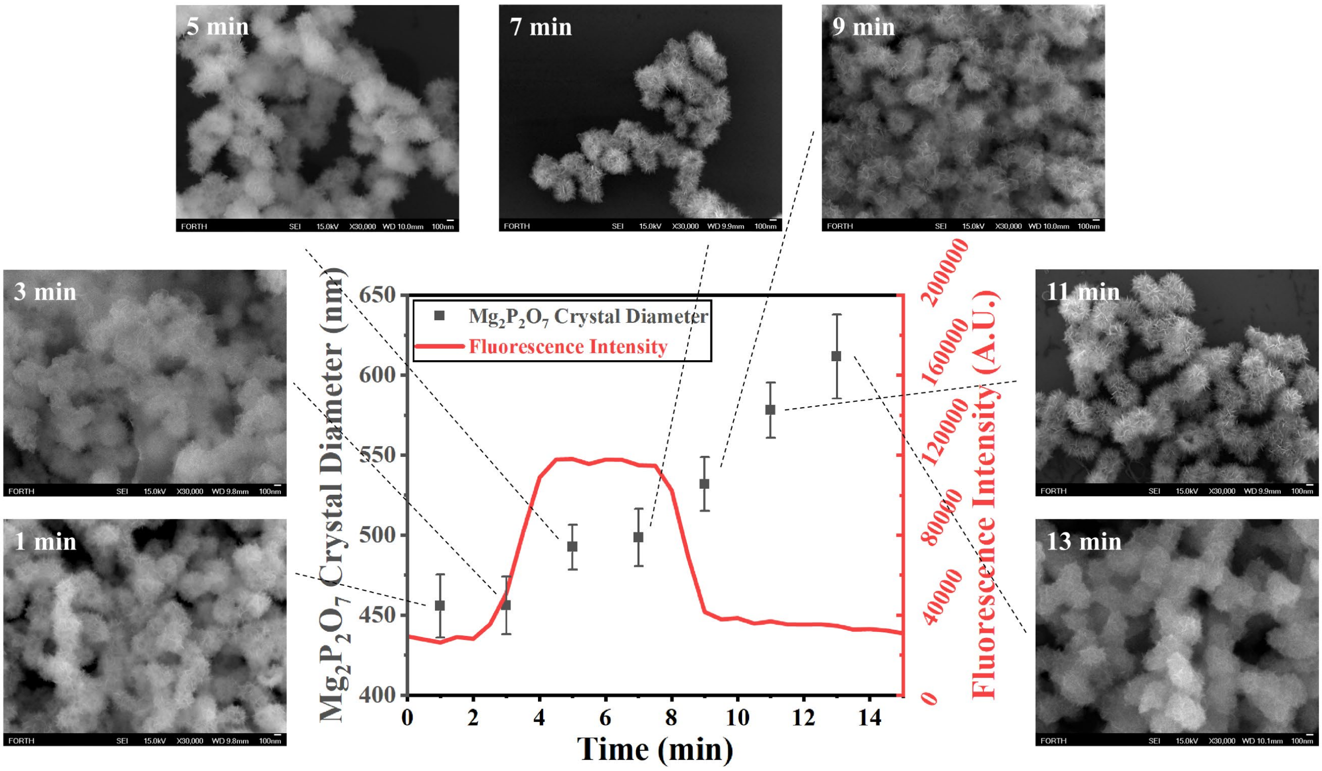

***Figure S6:*** *SEM images of $Mg_2P_2O_7$ crystals, obtained at different time intervals during the interaction of 12 mM $Mg^{2+}$ and 3 mM $P_2O_7^{4-}$, in the presence of 1 μL GSH-AuNCs (scale bar: 100 nm). A dual-y-axis graph depicting the real-time fluorescence graph, as well as the mean calculated size of the formed $Mg_2P_2O_7$ crystals at different time intervals, is also depicted. Note: drying of the SEM specimens was performed under vacuum, to minimize continued crystal growth and to ensure the size distribution at the time of collection is accurately reflected.*

As shown in **Fig. S6**, the $Mg_2P_2O_7$ crystals gradually increase in size over time, consistent with the expected progression from an initial nucleation stage under supersaturated conditions to subsequent crystal growth.[7,12,13] Quantification of crystal size at each time point revealed a progressive increase from an initial 455.77±19.73 nm and 456.13±18.07 nm at 1 and 3 min, respectively, to 492.65±13.96 nm at 5 min, 498.60±17.86 nm at 7 min, 531.90±16.82 nm at 9 min, 578.05±17.28 nm at 11 min, and 611.64±26.31 nm at 13 min. The relatively small change observed during the earliest time points and the more pronounced increase in crystal size at later stages are consistent with an initial period dominated by the formation of new crystal nuclei, followed by the preferential growth of the already-formed crystals. As the reaction proceeds, $Mg^{2+}$ and $P_2O_7^{4-}$ are progressively consumed, thereby making the formation of additional nuclei less favorable relative to the continued growth of existing crystals.[7,12,13] This interpretation is consistent with the temporal evolution observed, where the crystal population appears to become established at earlier time points, while the size of individual crystals continues to increase during the later time points. The relevant real-time fluorescence graph is provided to guide the eye, without necessarily implying that SEM time points overlap completely with those of the fluorescence graph.

**S7.** Size of $Mg_2P_2O_7$ crystals formed in the presence of GSH-AuNCs and various [$P_2O_7^{4-}$], [$(NH_4)_2SO_4$] or [dNTPs]

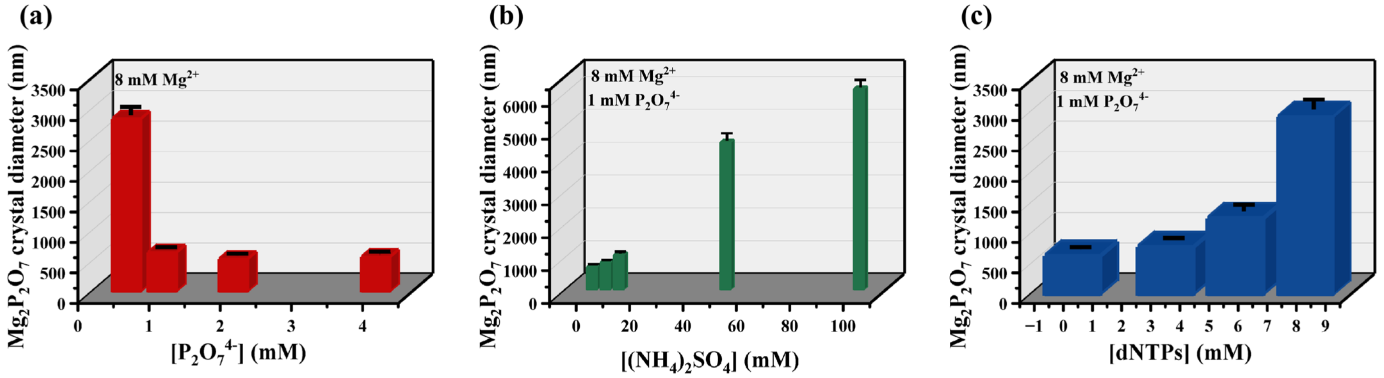

***Figure S7:*** *Bar charts representing the effect of varying (a) [$P_2O_7^{4-}$], (b) [$(NH_4)_2SO_4$] and (c) [dNTPs], on the size of $Mg_2P_2O_7$ crystals.*

SEM images were obtained to study the morphology of $Mg_2P_2O_7$ crystals formed when $Mg^{2+}$ (8 mM) and GSH-AuNCs (1 µL per 25 µL reaction) were incubated with various [$P_2O_7^{4-}$] (0.5, 1, 2 and 4 mM). Furthermore, a combination of 8 mM $Mg^{2+}$ with 1 mM $P_2O_7^{4-}$ and the same GSH-AuNCs volume (1 µL) was utilized to study the effect of various amounts of [$(NH_4)_2SO_4$] (5, 10, 50, 100 mM) or [dNTPs] (0, 3.2, 5.6 and 8 mM) on the morphology and size of the $Mg_2P_2O_7$ crystals.

In the case of varying [$P_2O_7^{4-}$] (**Fig. S7a**), larger crystals (2846.5±131.9 nm) were obtained in the presence of the lowest concentration (0.5 mM) compared with those formed using 1-4 mM $P_2O_7^{4-}$ (651.2±26.7 nm, 546.3±22.6 nm, and 580.7±25.0 nm, respectively). These findings are consistent with the concept that lower [$P_2O_7^{4-}$] favours the growth of fewer, larger crystals, whereas higher [$P_2O_7^{4-}$] promotes the formation of a greater number of smaller crystals.

The size-dependency of GSH-AuNCs/$Mg_2P_2O_7$ crystals as a function of [$(NH_4)_2SO_4$] and [dNTPs] is shown in **Fig. S7b** and **Fig. S7c**, respectively. In both cases, increasing the concentration of each reagent resulted in the formation of progressively larger crystalline structures. Specifically, the average crystal sizes (651.2±26.7 nm) formed using 8 mM $Mg^{2+}$ and 1 mM $P_2O_7^{4-}$ increased progressively in the presence of 5, 10, 50 and 100 mM $(NH_4)_2SO_4$ (840.9±26.3 nm, 1067.8±54.3 nm, 4498.7±225.1 nm and 6127.3±223.9 nm, respectively), implying crystal growth alterations. Similarly, formation of progressively larger crystals in the presence of 3.2, 5.6 and 8 mM dNTPs (799.6±30.8 nm, 1270.3±102.8 nm and 2938.4±157.7 nm, respectively) was also observed. This is attributed to interactions between dNTPs and $Mg^{2+}$,[14] which lowers the free $Mg^{2+}$ available to react with $P_2O_7^{4-}$, thereby shifting the [$Mg^{2+}$]/[$P_2O_7^{4-}$] ratio and altering crystal growth dynamics; incorporation of dNTPs in the formed $Mg_2P_2O_7$ crystals may also contribute.[15,16]

Overall, the above results confirm that [$Mg^{2+}$]/[$P_2O_7^{4-}$] ratio and the presence [$(NH_4)_2SO_4$] and [dNTPs] influence $Mg_2P_2O_7$ crystal size, in line with the real-time fluorescence data discussed in the main text.

**S8.** Real-time fluorescence monitoring of $Mg_2P_2O_7$ formation at 25 °C, using GSH-AuNCs

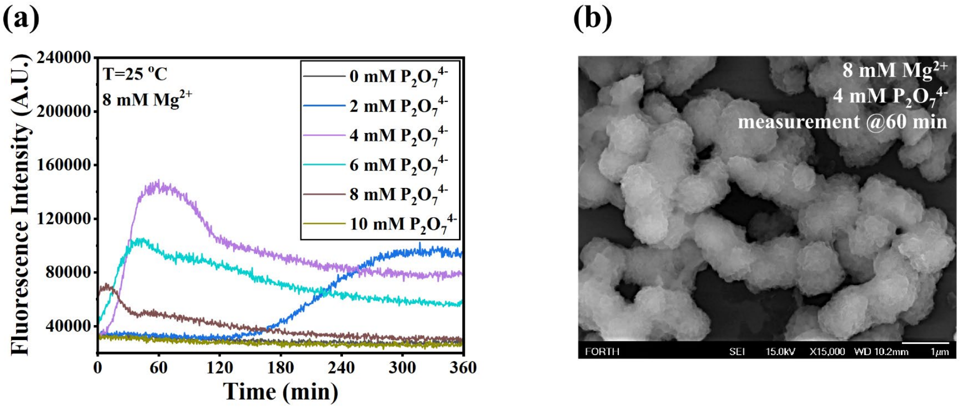

***Figure S8:*** *(a) Real-time fluorescence monitoring of $Mg_2P_2O_7$ formation using GSH-AuNCs (1 μL), in 25 μL reactions containing 8 mM $Mg^{2+}$ and 2-10 mM $P_2O_7^{4-}$ in the absence of heating (25 °C). (b) Representative SEM image of the formed $Mg_2P_2O_7$ crystals at 60 min (scale bar: 1 μm).*

Differences observed in the fluorescence response (*i.e.*, $Mg_2P_2O_7$ formation) at 25 °C compared to the experiment performed at 65 °C are consistent with the crystallization kinetics, where nucleation and growth parameters are temperature-dependent and $Mg_2P_2O_7$ formation proceeds through nucleation/precipitation processes.[13,17] Despite the slower kinetics observed at 25 °C, the "hat-shaped" behavior (initial fluorescence increase followed by decrease and stabilization) was retained overall (**Fig. S8a**). Finally, the fluorescence stabilization at higher $F_{end}$ values implies differences in the formed $Mg_2P_2O_7$ crystal size. We assume that larger crystalline structures may be formed in the absence of heating due to slower reaction kinetics. As shown in **Fig. S8b**, larger crystals (910±91.01 nm) are observed compared to crystals formed at 65 °C. Note that in this experiment, the size was measured at 60 min as a direct comparison to the 65 °C experiment (**Fig. S7a**), although here the crystal growth/reaction may continue afterwards, leading to the formation of even larger crystals at the end of the reaction.

**S9.** Effect of different GSH-AuNCs amounts on the $Mg_2P_2O_7$-induced fluorescence enhancement

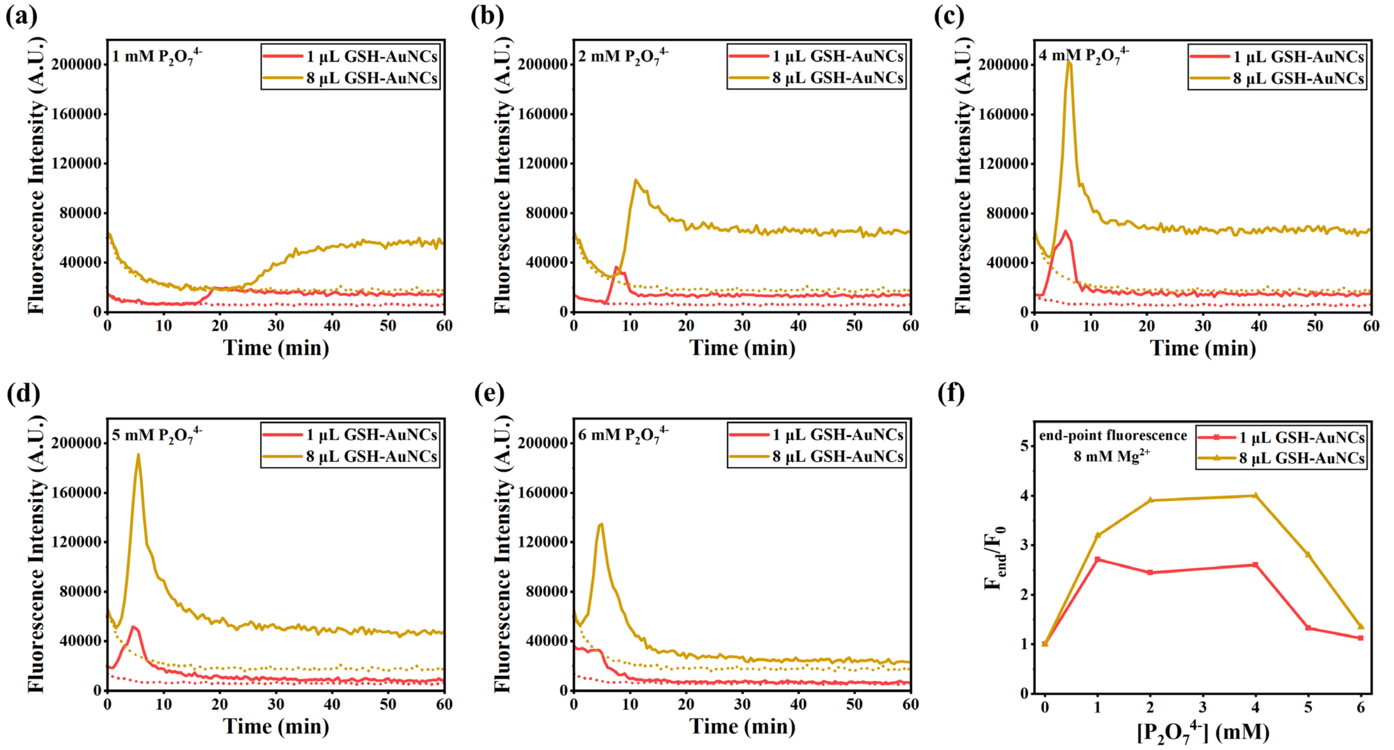

***Figure S9:*** *(a-e) Real-time fluorescence graphs of different GSH-AuNCs volumes (1 and 8 µL), in the presence of 8 mM* $Mg^{2+}$ *and 1-6 mM* $P_2O_7^{4-}$ *(65 °C) in 25 µL reactions. (f) Relative fluorescence intensity of the end-point fluorescence (*$F_{end}/F_0$*) against [*$P_2O_7^{4-}$*] of the reactions presented in (a-e). Note: to avoid signal overflow due to the* $Mg_2P_2O_7$*-induced fluorescence enhancement of GSH-AuNCs, gain adjustment was carried out using a 25 µL well containing 12 µL GSH-AuNCs with* $Mg_2P_2O_7$*, formed using 8 mM* $Mg^{2+}$ *and 4 mM* $P_2O_7^{4-}$*.*

A total of five different [$P_2O_7^{4-}$] combined with 8 mM $Mg^{2+}$ were used, in the presence of either 1 or 8 µL GSH-AuNCs. When a low [$P_2O_7^{4-}$] is used, the time-to-positivity (TTP) is longer with increasing GSH-AuNCs volume (**Fig. S9a**), indicating an effect of the GSH-AuNCs fluorescent probe on the formed $Mg_2P_2O_7$ crystals; however, the TTP is not significantly affected at higher [$P_2O_7^{4-}$] (**Fig. S9b-e**). Overall, the elevated volume of GSH-AuNCs (8 µL) provides an increased end-point fluorescent signal intensity, with the $F_{end}$ trend more closely matching the trend in solid $Mg_2P_2O_7$ formation monitoring (**Fig. 1f**), indicated by the end-point absorbance/turbidity measurements. In addition, the $F_{end}$ trend with 1 µL GSH-AuNCs does not closely follow the solid $Mg_2P_2O_7$ formation, suggesting that all GSH-AuNCs may already be confined within the lower $Mg_2P_2O_7$ concentration, formed with 8 mM $Mg^{2+}$ and 1 mM $P_2O_7^{4-}$ (**Fig. S9f**).

**S10.** SEM images of $Mg_2P_2O_7$ crystals formed in the absence or in the presence of 1 or 3 μL GSH-AuNCs

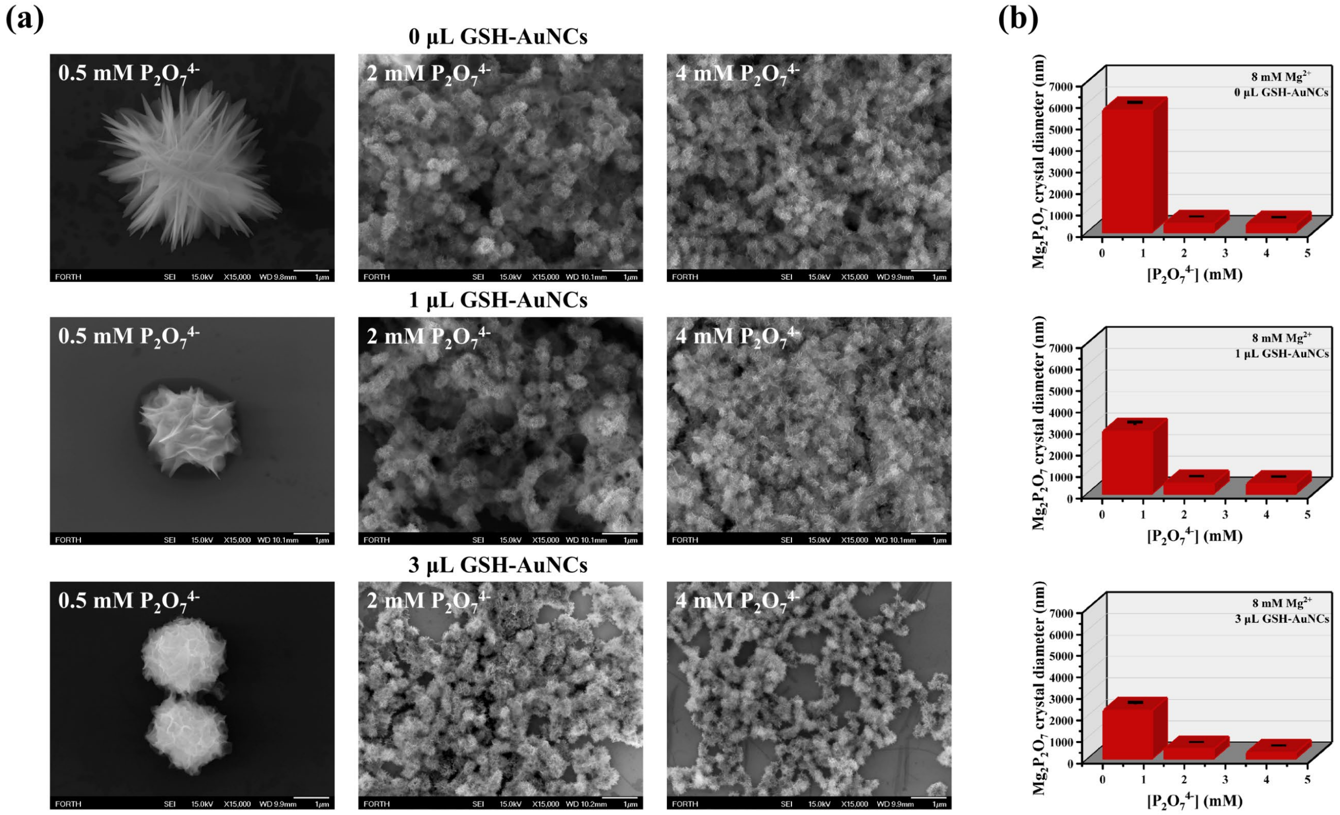

***Figure S10:*** *(a) SEM images of $Mg_2P_2O_7$ crystals formed after incubating 8 mM $Mg^{2+}$ with 0.5, 2 and 4 mM $P_2O_7^{4-}$ in the absence, or in the presence of either 1 or 3 μL GSH-AuNCs per 25 μL reaction (scale bar: 1 μm). (b) Bar charts comparing the size of the formed $Mg_2P_2O_7$ crystals.*

Based on the SEM images (**Fig. S10a**), it can be seen that for low [$P_2O_7^{4-}$] (0.5 mM), fewer and larger crystals are formed, while for 2 and 4 mM $P_2O_7^{4-}$, more $Mg_2P_2O_7$ crystals of smaller size are formed, in agreement with the results presented in **Fig. 2i**. However, in the case of 0.5 mM $P_2O_7^{4-}$, an effect of GSH-AuNCs on the $Mg_2P_2O_7$ crystal size can be observed. Specifically, in the absence of GSH-AuNCs, larger crystals of 5709±67.1 nm size are formed, while they are smaller at 2931.9±135.7 nm and 2274.9±77.6 nm in the presence of 1 and 3 μL GSH-AuNCs, respectively. This implies an effect of the GSH-AuNCs fluorescent probe on the $Mg_2P_2O_7$ crystal formation, altering the crystal growth dynamics and affecting their size and possibly their concentration. Notably, at higher [$P_2O_7^{4-}$], no significant size variations of $Mg_2P_2O_7$ crystals were observed by varying amounts of GSH-AuNCs (**Fig. S10b**), indicating that the fluorescent probe effect is more significant when fewer crystals are formed (*i.e.*, low $Mg^{2+}$ and/or $P_2O_7^{4-}$ concentrations).

#### **S11.** Reduced equilibrium $Mg^{2+}$-$P_2O_7^{4-}$ speciation and supersaturation ratios (Python-simulated)

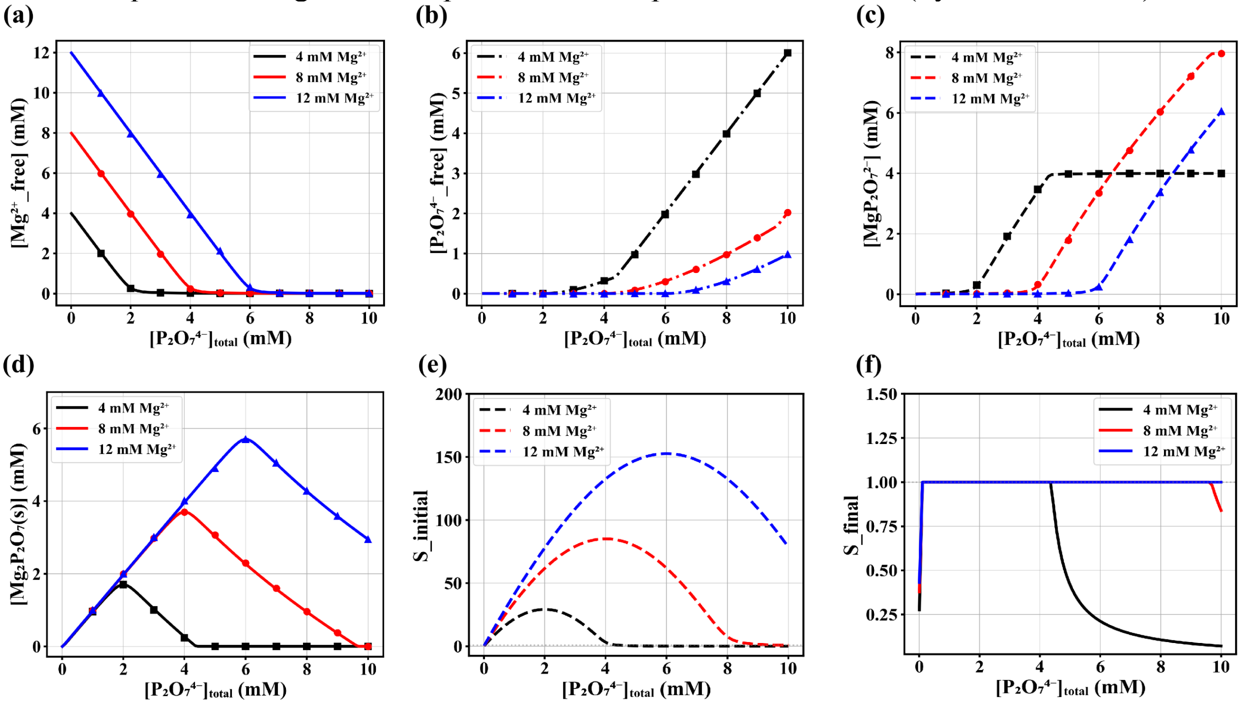

***Figure S11:*** *Theoretical calculations of the concentrations of: (a) free* $Mg^{2+}$, (b) free $P_2O_7^{4-}$, *(c)* $MgP_2O_7^{2-}$, *and (d) solid* $Mg_2P_2O_7$, *as well as (e)* $S_{initial}$ *and (f)* $S_{final}$ *at equilibrium, during the reaction of 4, 8 and 12 mM* $Mg^{2+}$ *with 0-10 mM* $P_2O_7^{4-}$. *Note: the script was written in Python.*

The dependence of $Mg_2P_2O_7$ precipitation on $[P_2O_7^{4-}]$ was modeled using a reduced equilibrium model (omitting pH-dependent $P_2O_7^{4-}$ protonation, ionic strength effects, activity coefficients, buffer effects, and other competing solution components). Free $Mg^{2+}$ and $P_2O_7^{4-}$ first form the 1:1 aqueous $MgP_2O_7^{2-}$ complex, followed by formation of the 2:1 aqueous $Mg_2P_2O_7$ species:

$$Mg^{2+} + P_2O_7{}^{4-} \underset{K_1}{\rightleftharpoons} MgP_2O_7{}^{2-} \underset{K_2}{\overset{Mg^{2+}}{\rightleftharpoons}} Mg_2P_2O_7 \qquad (1)$$

with reported stability constants $K_1=10^{5.41}$ $M^{-1}$ and $K_2=10^{2.34}$ $M^{-1}$, respectively, under alkaline conditions.[18,19] Precipitation of $Mg_2P_2O_7$ is then evaluated using the free-ion product:

$$Q = [Mg^{2+}]^2[P_2O_7{}^{4-}] \qquad (2)$$

where it is thermodynamically favored when $Q > K_{sp}(T)$. The apparent concentration-form solubility product for $Mg_2P_2O_7 \cdot 3.5H_2O$ was taken as $K_{sp} = 1.63\cdot10^{-13}$ $M^3$ at 25 °C,[7] while temperature corrections to 65 °C were applied to $K_1$ and $K_2$ values where appropriate, using van't Hoff relationships and reported enthalpy values.[7]

More specifically, the model calculates free $Mg^{2+}$, free $P_2O_7^{4-}$, $MgP_2O_7^{2-}$ (**Fig. S11a-c**), precipitated/solid $Mg_2P_2O_7$ (**Fig. S11d**) and saturation profiles (**Fig. S11e, f**), as a function of total $[P_2O_7^{4-}]$ (0-10 mM) at fixed total $[Mg^{2+}]$ (4, 8 and 12 mM), using Brent's root-finding method.[20] $S_{initial}=Q_{initial}/K_{sp}$ is the no solid/pre-precipitation supersaturation ratio, equivalently calculated from the initial precursor $Mg_2P_2O_{7(aq)}$, and can therefore represent the initial thermodynamic driving force for $Mg_2P_2O_{7(s)}$ precipitation (**Fig. S11e**). The $S_{initial}$ trend can be qualitatively associated with stronger initial supersaturation and precipitation tendency, following a pattern comparable to the $F_{max}$ trend obtained through our real-time fluorescence measurements (**Fig. 2e**). In contrast, $S_{final}=Q_{final}/K_{sp}$ is the residual post-precipitation supersaturation ratio after applying the equilibrium precipitation step (**Fig. S11f**). The $S_{final}$ trend reflects the saturation state of the remaining aqueous phase after the equilibrium precipitation step. Values near 1 indicate conditions where $Mg_2P_2O_{7(s)}$ is predicted and the remaining solution is returned to the solubility boundary, whereas values below 1 indicate undersaturated final aqueous conditions, where no $Mg_2P_2O_{7(s)}$ precipitation is thermodynamically favored. Finally, the predicted $Mg_2P_2O_{7(s)}$ trend can be qualitatively compared with absorbance/turbidity measurements (**Fig. 1f**), as well as the $F_{end}$ trend when sufficient GSH-AuNCs are present in solution (**Fig. S9f**). Deviations between simulated precipitation trends and experimental responses are expected, since turbidity and fluorescence depend not only on the total precipitated amount but also on particle size, aggregation state, and crystal morphology.[21] For example, agreement with $F_{end}$ may be expected mainly under conditions where the formed $Mg_2P_2O_7$ crystals have comparable sizes; deviations may occur at high $[Mg^{2+}]/[P_2O_7^{4-}]$ ratios, where larger crystals can form under slower crystallization conditions, which may alter AIE-like fluorescence enhancement. These values should therefore be interpreted as reduced equilibrium trends rather than kinetic predictions of nucleation rate, crystal number, particle size, or crystal morphology.

**S12.** Turbidity measurements *via* absorbance (610 nm) of solutions containing a combination of 8 mM $Mg^{2+}$, 1 mM $P_2O_7^{4-}$ and various [dNTPs]

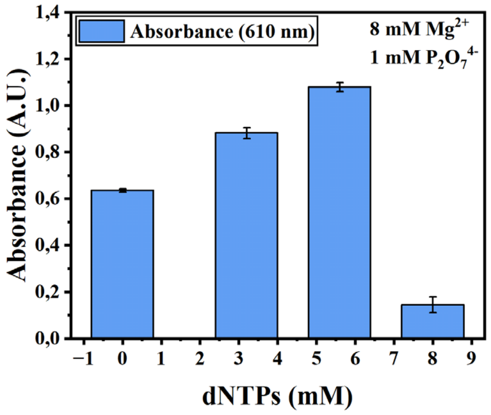

***Figure S12:*** *Absorbance measurements at 610 nm for the solutions containing 1 µL GSH-AuNCs, and $Mg_2P_2O_7$ formed using 8 mM $Mg^{2+}$, 1 mM $P_2O_7^{4-}$, and either 0, 3.2, 5.6 or 8 mM dNTPs.*

At a high dNTP concentration (8 mM), the turbidity/absorbance of the solution is significantly decreased compared to the absence of dNTPs (0 mM). This is probably due to insufficient remaining free $Mg^{2+}$ to interact with $P_2O_7^{4-}$,[14] resulting in less $Mg_2P_2O_7$ formation, while the soluble $MgP_2O_7^{2-}$ form dominates as the $[Mg^{2+}]/[P_2O_7^{4-}]$ ratio is close to 1 (**Fig. S12**). However, at intermediate dNTP concentrations (3.2 and 5.6 mM), an increase in turbidity is observed, likely reflecting differences in the formed $Mg_2P_2O_7$ crystals (**Fig. S7c**, see also **Fig. 3f**).

**S13.** Effect of [dNTPs] on the $Mg_2P_2O_7$-induced fluorescence enhancement of GSH-AuNCs

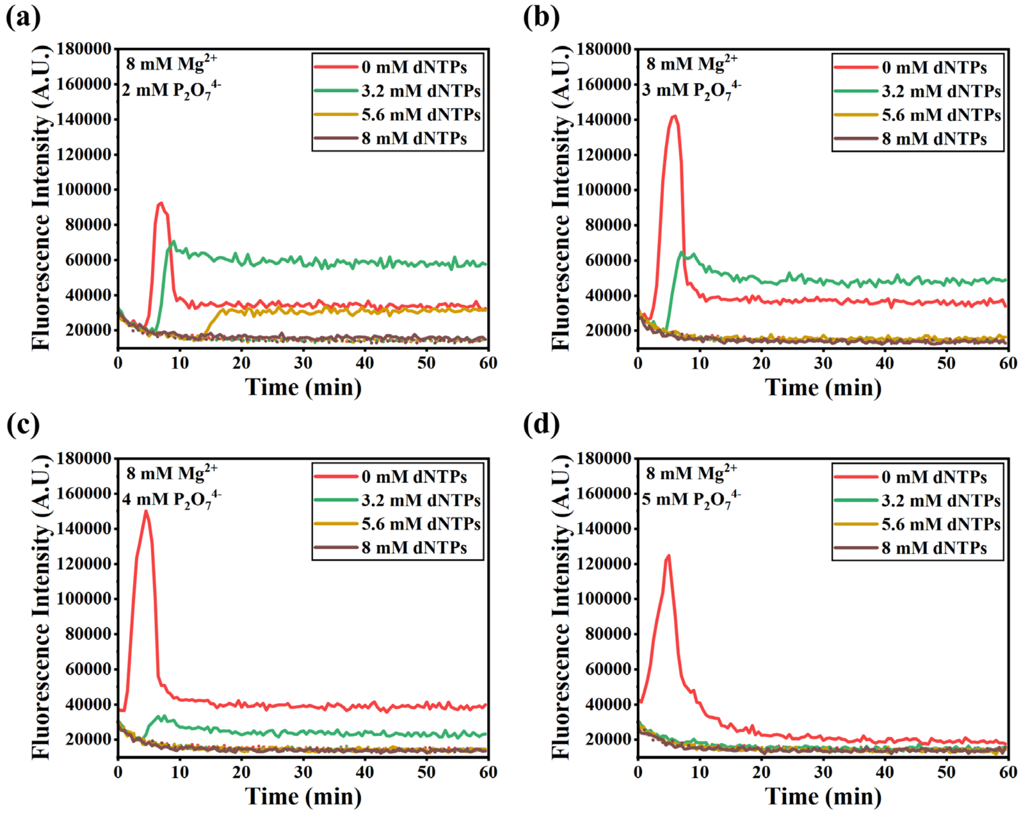

***Figure S13:*** *(a-d) Effect of varying [dNTPs] (3.2, 5.6 and 8 mM) on the $Mg_2P_2O_7$-induced fluorescence enhancement of GSH-AuNCs (1 µL in 25 µL reactions), formed using 8 mM $Mg^{2+}$ and 2, 3, 4 or 5 mM $P_2O_7^{4-}$ (65 °C).*

According to **Fig. S13a-d**, using high [dNTPs] (8 mM) results in no fluorescence enhancement for any of the [$P_2O_7^{4-}$] used in this experiment. This observation can be rationalized by the near 1:1 binding stoichiometry[14] of dNTPs with $Mg^{2+}$, which may reduce the pool of free $Mg^{2+}$ available to react with $P_2O_7^{4-}$. As a result, either $Mg_2P_2O_7$ formation is suppressed, or the effective [$Mg^{2+}$]/[$P_2O_7^{4-}$] ratio falls below 1, favoring soluble $MgP_2O_7^{2-}$ species instead of the insoluble $Mg_2P_2O_7$ phase. However, at lower final dNTP concentrations, *i.e.*, 3.2 and 5.6 mM, there is sufficient free $Mg^{2+}$ to produce $Mg_2P_2O_7$ as the dominant species. The above observations, including results from **Fig. 3e, f** and **Fig. S7c**, confirm that the presence of dNTPs can modulate $Mg_2P_2O_7$ formation and thereby the fluorescence response of GSH-AuNCs.

**S14.** Real-time fluorescent monitoring of LAMP reactions using GSH-AuNCs, utilizing different [Bst] and [dNTPs] combinations

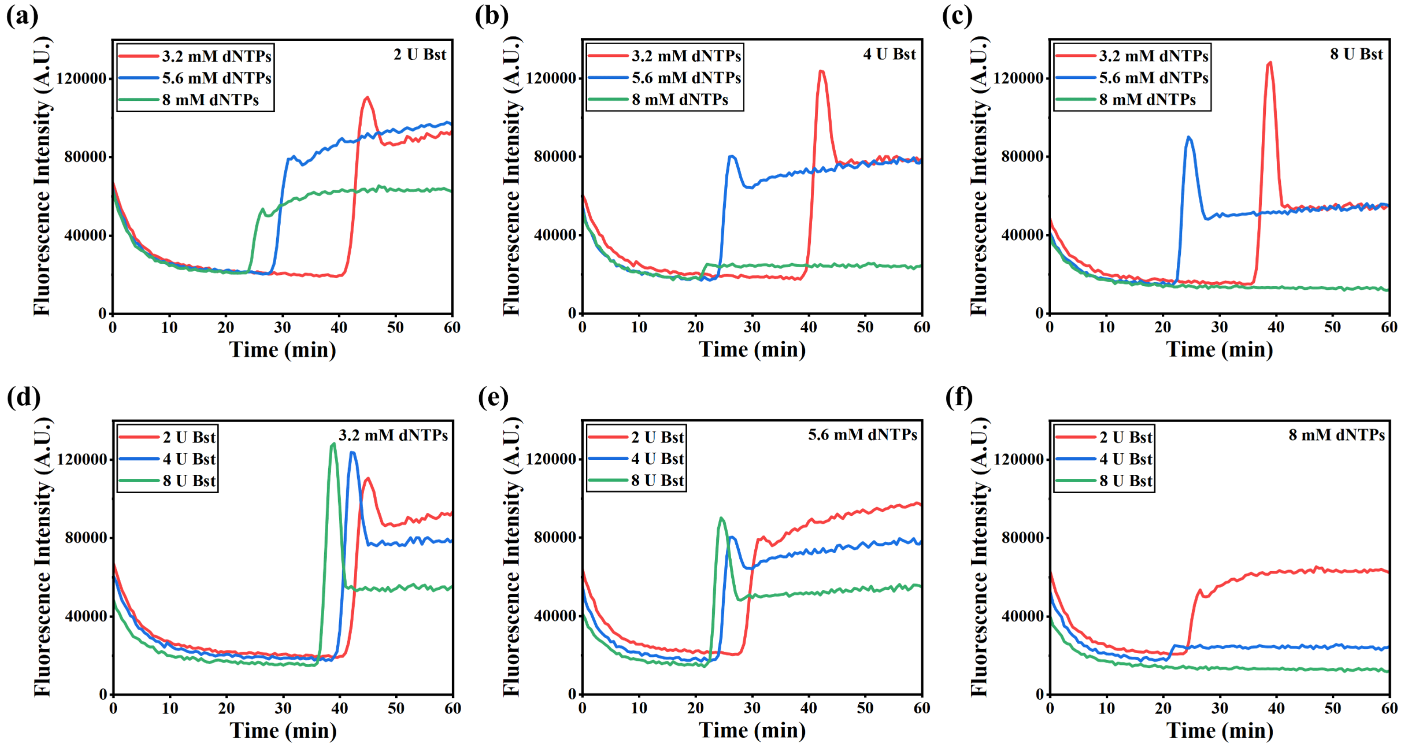

***Figure S14:*** *Comparative graphs depicting the effect of different [dNTPs] (3.2, 5.6 and 8 mM) with (a) 2, (b) 4, or (c) 8 U Bst, and different [Bst] (2, 4, 8 U) with (d) 3.2, (e) 5.6, or (f) 8 mM dNTPs, on the GSH-AuNCs fluorescence enhancement, induced by* $Mg_2P_2O_7$ *formation. Note: All 25 µL LAMP reactions (65 °C) contained* $10^5$ *Influenza A RNA copies/reaction, along with 3 µL GSH-AuNCs.*

Based on the data presented in **Fig. S14a-f**, a longer TTP is generally observed when using low Bst or low dNTP concentrations. With low [dNTPs], there are fewer “building blocks” available, which the Bst polymerase enzyme uses to synthesize new DNA strands, even if plenty of Bst is present, leading to a slower reaction.[22] In all cases (2, 4, 8 U Bst), using 3.2 mM dNTPs results in a significantly longer TTP compared to the use of 5.6 mM dNTPs. However, excessively high dNTP concentrations can also inhibit the reaction by interfering with Bst activity. Similarly, with low Bst concentrations, fewer active Bst polymerase molecules are available to extend the DNA primers and thus, synthesize new strands of DNA, which also leads to a slower reaction. Interestingly, the use of low [Bst] (2 U) combined with high [dNTPs] (8 mM) provided a distinct fluorescent signal with an early TTP.

Regarding our GSH-AuNCs detection method, the slower kinetics of the reaction (*i.e.*, dNTPs to $P_2O_7^{4-}$ conversion) when using low [Bst] result in the fluorescent signal being stabilized, especially in combination with high dNTP concentrations relative to the Bst concentration. This is probably due to altered crystal growth dynamics, possible incorporation of dNTPs into the formed $Mg_2P_2O_7$ crystals due to slower dNTPs to $P_2O_7^{4-}$ conversion, and/or the subsequent formation of larger crystals (see also **Fig. 3f** and **Fig. S7c**).

**S15.** Real-time quantitative colorimetric LAMP using HNB dye, with either absorbance measurements (Omega plate reader) or image analysis (Pebble)

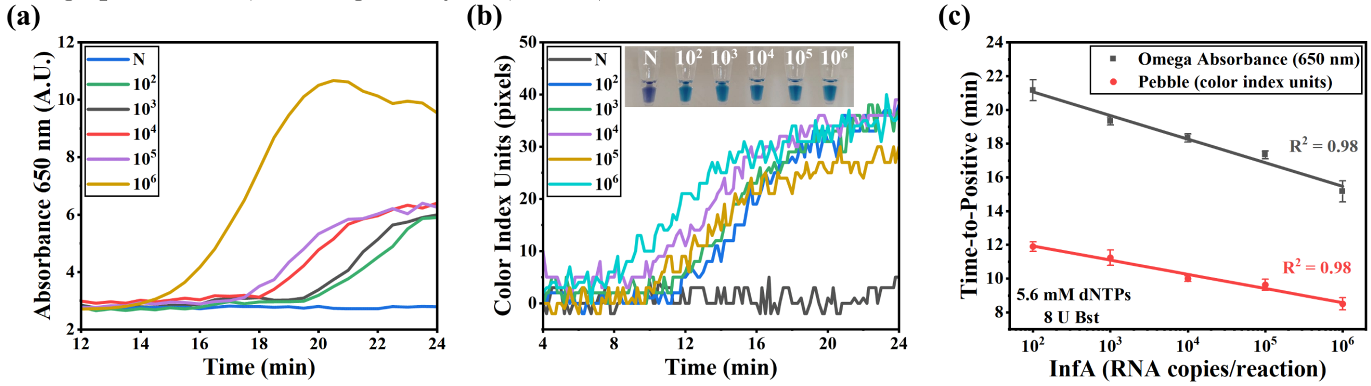

***Figure S15:*** *Real-time colorimetric LAMP (qcLAMP) signals obtained with: (a) absorbance measurements at 650 nm using the Omega plate reader, and (b) image analysis using the Pebble device. Inset: photograph of the solutions, revealing the HNB color change from violet (negative) to sky blue (positive). (c) Calibration curves derived from the above real-time colorimetric measurements. Note: LAMP reactions contained* $10^2$*-*$10^6$ *Influenza A copies/reaction, while each reaction contained a final 120 μM HNB.*

Real-time quantitative colorimetric reactions employing the commercially available hydroxynaphthol blue (HNB) dye were performed in the Omega plate reader (**Fig. S15a**) and the commercially available Pebble device[23] (**Fig. S15b**), in triplicate. In the first case, colorimetric monitoring was performed using absorbance measurements at 650 nm, while colorimetric image analysis was used for the results obtained *via* Pebble. Influenza A RNA target within a concentration range of $10^2$-$10^6$ copies/reaction was used in both cases, to enable comparison with the performance of the GSH-AuNCs fluorescent probe.

According to **Fig. S15c**, a linear correlation ($R^2$=0.98) was obtained between the TTP and the number of RNA copies in the initial sample, for both instruments. Using the Omega plate reader, we obtained response times in the range of 21±0.6 and 16±0.6 min for the lowest and highest concentrations, respectively, a range similar to that of the GSH-AuNC assay in the same instrument. However, the overall assay using the Pebble device was significantly faster, a result that could be attributed to more efficient heating at 65 °C in this case.

It is finally noted that the HNB dye was preferred over the calcein dye, which, although it binds to $Mg^{2+}$ and is fluorescent, requires the use of manganese ions ($Mn^{2+}$), making it less suitable for comparison with our assay.[24]
