## Supplementary Code (Python) for "Mechanistic Insights into Magnesium Pyrophosphate Formation in the Presence of Gold Nanoclusters Enable Genetic Analysis via Co-Aggregation-Induced Fluorescence Enhancement": mg_p2o7_equilibrium.py

```
# -*- coding: utf-8 -*-
"""Mg2+-P2O7(4-) equilibrium speciation and precipitation model.

The aqueous network contains free Mg2+, free P2O7(4-), MgP2O7(2-), and
Mg2P2O7(aq). ``solid_model="equilibrium"`` adds one Mg2P2O7(s) pool and
solves the long-time saturation endpoint; ``solid_model="none"`` reports the
no-solid aqueous upper bound.

This is not a complete reaction-mixture model: pH-dependent P2O7
protonation, activities, ionic-strength iteration, buffers, competing ligands,
and alternative solid phases are outside scope. Edit the configuration objects
below, then run the file. Importing the module does not calculate, plot, or
write files.
"""

from __future__ import annotations

import csv
import math
from dataclasses import asdict, dataclass
from pathlib import Path
from typing import Iterable, Mapping, Sequence

import matplotlib as mpl
import matplotlib.pyplot as plt
import numpy as np
from scipy.optimize import brentq


# Parameter/source summary (equations and citations: README.md):
# - K1/K2: conditional Mg-pyrophosphate association constants from Kern-Davis.
# - dH_K1: +12.552 kJ/mol is the association-direction equivalent of the
#   -12.552 kJ/mol MgPPi dissociation parameter used in the Ahmadi-Stover
#   implementation; dH_K2=0 disables K2 temperature transport here.
# - fixed Ksp: 1.63e-13 M^3 is the 25 °C Mg2P2O7·3.5H2O concentration-form
#   value fitted in the Ahmadi-Stover implementation. This compact model does
#   not apply their Ksp temperature transport.

# -----------------------------------------------------------------------------
# USER CONFIGURATION
# -----------------------------------------------------------------------------
# The dataclasses are immutable so one calculation retains a reproducible set
# of inputs. For a standalone run, edit these defaults. When importing, create a
# new config or use dataclasses.replace(), as shown in README.md.
@dataclass(frozen=True)
class ModelConfig:
    # "equilibrium" forms solid until S=1; "none" suppresses solid while
    # still reporting the counterfactual equilibrium amount s_eq_mM.
    solid_model: str = "equilibrium"

    # Calculation temperature. The reference K1/K2 values below are defined
    # at 25 °C, the baseline temperature of the source equilibrium calculations.
    # Set temperature_C=25.0 to use that reference temperature; other values
    # are applied to K1/K2 only when the correction toggle is True. Ksp is fixed.
    temperature_C: float = 65.0
    use_K1K2_temperature_correction: bool = True

    # Conditional association constants at 25 °C, in M^-1.
    # K1: Mg2+ + P2O7(4-) <-> MgP2O7(2-)
    # K2: Mg2+ + MgP2O7(2-) <-> Mg2P2O7(aq)
    log10_K1_ref: float = 5.42
    log10_K2_ref: float = 2.33

    # Enthalpies follow the association directions above. +12.552 kJ/mol is
    # the association-direction form used for K1 temperature transport. A zero
    # dH_K2 disables temperature transport of K2 in this parameterization.
    dH_K1_kJ_mol: float = 12.552
    dH_K2_kJ_mol: float = 0.0

    # Fixed concentration-form threshold, Ksp=[Mg2+]^2[P2O7(4-)] in M^3.
    # Ksp temperature correction and alternative threshold-basis selection are
    # not implemented in this script.
    Ksp_M3: float = 1.63e-13

    # Main sweep: one P2O7 grid is calculated for each Mg total.
    mg_totals_mM: tuple[float, ...] = (4.0, 8.0, 12.0)
    p2o7_min_mM: float = 0.01
    p2o7_max_mM: float = 10.0
    p2o7_points: int = 100


@dataclass(frozen=True)
class PlotConfig:
    # Preview enabled outputs in 2×2 pages (four parameters per window).
    # Figure export remains one standalone file per enabled parameter.
    enabled: bool = True

    # Plot appearance. Series begin with black/red/blue colors and s/o/^
    # markers. Any number of Mg totals is supported. Marker styles are
    # cycled safely when there are more Mg series than marker entries.
    show_lines: bool = True
    line_width: float = 3.2

    # Series styling. Colors and markers are assigned by series order.
    # The first three defaults reproduce the black/red/blue and s/o/^
    # sequence. If more series are plotted than colors are listed,
    # additional colors are generated deterministically from the seed.
    series_colors: tuple[str, ...] = ("black", "red", "blue")
    extra_color_seed: int = 42
    show_markers: bool = True
    marker_styles: tuple[str, ...] = ("s", "o", "^", "D", "v", "<", ">", "p", "*", "H")
    marker_every: int = 10
    marker_size: float = 7.0

    # Variable-specific line styles. These apply to both preview and export.
    linestyle_Mg2_free: str = "-"
    linestyle_P2O7_free: str = "-."
    linestyle_MgP2O7: str = "--"
    linestyle_Mg2P2O7_aq: str = ":"
    linestyle_Mg2P2O7_initial: str = "--"
    linestyle_solid: str = "-"
    linestyle_S_initial: str = "--"
    linestyle_S_final: str = "-"
    # Size of one panel/standalone export. Preview pages are 2× this width
    # and 2× this height because they contain a 2×2 matrix.
    figure_width: float = 7.5
    figure_height: float = 5.5
    dpi: int = 300

    # Shared axes/frame emphasis for both preview and exports.
    spine_width: float = 2.5
    tick_width: float = 2.0
    tick_length: float = 6.5
    grid_line_width: float = 1.0
    grid_alpha: float = 0.55

    # Shared typography for preview and exported figures. Plot labels use the
    # original Unicode chemistry notation. Times New Roman is preferred when
    # available; DejaVu Serif is the portable fallback.
    use_times_new_roman: bool = True
    axis_label_size: float = 22.0
    tick_label_size: float = 19.0
    legend_size: float = 16.0
    legend_title_size: float = 16.0
    bold_text: bool = True

    # Toggle individual species/diagnostic figures.
    show_Mg2_free: bool = True
    show_P2O7_free: bool = True
    show_MgP2O7: bool = True
    show_Mg2P2O7_aq: bool = True
    show_Mg2P2O7_initial: bool = True
    show_solid: bool = True
    show_S_initial: bool = True
    show_S_final: bool = True


@dataclass(frozen=True)
class ExportConfig:
    directory: str = "outputs"
    export_table: bool = False
    table_format: str = "csv"  # "csv", "tsv", or "xlsx"
    table_stem: str = "mg_p2o7_equilibrium"

    # Each enabled output is saved as its own figure.
    export_figures: bool = False
    figure_format: str = "png"  # any Matplotlib-supported extension
    figure_stem: str = "mg_p2o7_equilibrium"


MODEL = ModelConfig()
PLOT = PlotConfig()
EXPORT = ExportConfig()

R_GAS_J_MOL_K = 8.31446261815324
T_REF_K = 298.15


# -----------------------------------------------------------------------------
# EQUILIBRIUM CHEMISTRY
# -----------------------------------------------------------------------------
@dataclass(frozen=True)
class Constants:
    K1_M_inv: float
    K2_M_inv: float
    K1_mM_inv: float
    beta_mM_inv2: float
    precursor_threshold_mM: float


@dataclass(frozen=True)
class PointResult:
    Mg_total_mM: float
    P2O7_total_mM: float
    Mg2_free_mM: float
    P2O7_free_mM: float
    MgP2O7_mM: float
    Mg2P2O7_aq_mM: float
    Mg2P2O7_initial_mM: float
    Mg2P2O7_s_mM: float
    S_initial: float
    S_final: float
    s_eq_mM: float


def _van_t_hoff(K_ref: float, dH_kJ_mol: float, temperature_C: float) -> float:
    """Transport an equilibrium constant from 25 °C with van 't Hoff."""
    T_K = temperature_C + 273.15
    dH_J_mol = dH_kJ_mol * 1000.0
    return K_ref * math.exp(
        -dH_J_mol / R_GAS_J_MOL_K * (1.0 / T_K - 1.0 / T_REF_K)
    )


def effective_constants(config: ModelConfig = MODEL) -> Constants:
    """Return molar constants, mM-scaled solver constants, and c2_eq.

    The aqueous equations use mM, so each M^-1 stepwise constant is multiplied
    by 1e-3. The equivalent precursor threshold is derived in molar units as
    [Mg2P2O7(aq)]eq=K1*K2*Ksp and converted to mM.
    """
    if not math.isfinite(config.Ksp_M3) or config.Ksp_M3 <= 0.0:
        raise ValueError("Ksp_M3 must be finite and positive.")

    K1_M_inv = 10.0 ** config.log10_K1_ref
    K2_M_inv = 10.0 ** config.log10_K2_ref

    if config.use_K1K2_temperature_correction:
        K1_M_inv = _van_t_hoff(K1_M_inv, config.dH_K1_kJ_mol, config.temperature_C)
        K2_M_inv = _van_t_hoff(K2_M_inv, config.dH_K2_kJ_mol, config.temperature_C)

    if not math.isfinite(K1_M_inv) or K1_M_inv <= 0.0:
        raise ValueError("Effective K1 must be finite and positive.")
    if not math.isfinite(K2_M_inv) or K2_M_inv <= 0.0:
        raise ValueError("Effective K2 must be finite and positive.")

    K1_mM_inv = K1_M_inv * 1e-3
    K2_mM_inv = K2_M_inv * 1e-3
    return Constants(
        K1_M_inv=K1_M_inv,
        K2_M_inv=K2_M_inv,
        K1_mM_inv=K1_mM_inv,
        beta_mM_inv2=K1_mM_inv * K2_mM_inv,
        precursor_threshold_mM=K1_M_inv * K2_M_inv * config.Ksp_M3 * 1e3,
    )


def _validate_total(name: str, value: float) -> float:
    value = float(value)
    if not math.isfinite(value) or value < 0.0:
        raise ValueError(f"{name} must be finite and non-negative.")
    return value


def solve_aqueous_speciation(
    Mg_total_mM: float,
    P2O7_total_mM: float,
    constants: Constants,
) -> tuple[float, float, float, float]:
    """Solve free Mg2+, free P2O7(4-), MgP2O7(2-), and Mg2P2O7(aq).

    All totals and returned concentrations are in mM. The P2O7 balance gives

        p = P_total / (1 + K1*m + K1*K2*m^2),

    and substitution into the Mg balance leaves one scalar root in free Mg m.
    """
    Mg_total_mM = _validate_total("Mg_total_mM", Mg_total_mM)
    P2O7_total_mM = _validate_total("P2O7_total_mM", P2O7_total_mM)

    if Mg_total_mM == 0.0 or P2O7_total_mM == 0.0:
        return Mg_total_mM, P2O7_total_mM, 0.0, 0.0

    K1 = constants.K1_mM_inv
    beta = constants.beta_mM_inv2

    def mg_balance_residual(m: float) -> float:
        p = P2O7_total_mM / (1.0 + K1 * m + beta * m * m)
        return m + K1 * m * p + 2.0 * beta * m * m * p - Mg_total_mM

    # A conservative non-negative upper bound contains the physical free-Mg
    # root while preserving a simple bracketed solve.
    upper = Mg_total_mM + 2.0 * P2O7_total_mM + 10.0
    m = brentq(mg_balance_residual, 0.0, upper, maxiter=1000)
    p = P2O7_total_mM / (1.0 + K1 * m + beta * m * m)
    c1 = K1 * m * p
    c2 = beta * m * m * p
    return max(m, 0.0), max(p, 0.0), max(c1, 0.0), max(c2, 0.0)


def speciation_after_solid(
    Mg_total_mM: float,
    P2O7_total_mM: float,
    solid_mM: float,
    constants: Constants,
) -> tuple[float, float, float, float]:
    """Re-solve aqueous speciation after forming Mg2P2O7(s)."""
    Mg_total_mM = _validate_total("Mg_total_mM", Mg_total_mM)
    P2O7_total_mM = _validate_total("P2O7_total_mM", P2O7_total_mM)
    solid_mM = _validate_total("solid_mM", solid_mM)
    maximum = min(P2O7_total_mM, Mg_total_mM / 2.0)
    if solid_mM > maximum + 1e-10:
        raise ValueError("solid_mM exceeds the stoichiometric maximum.")
    solid_mM = min(solid_mM, maximum)
    return solve_aqueous_speciation(
        Mg_total_mM - 2.0 * solid_mM,
        P2O7_total_mM - solid_mM,
        constants,
    )


def ion_product_M3(Mg2_free_mM: float, P2O7_free_mM: float) -> float:
    """Return Q = [Mg2+]^2[P2O7(4-)] in M^3."""
    return (Mg2_free_mM * 1e-3) ** 2 * (P2O7_free_mM * 1e-3)


def supersaturation(Mg2_free_mM: float, P2O7_free_mM: float, Ksp_M3: float) -> float:
    """Return the concentration-form supersaturation ratio Q/Ksp."""
    if not math.isfinite(Ksp_M3) or Ksp_M3 <= 0.0:
        raise ValueError("Ksp_M3 must be finite and positive.")
    return ion_product_M3(Mg2_free_mM, P2O7_free_mM) / Ksp_M3


def solve_equilibrium_solid(
    Mg_total_mM: float,
    P2O7_total_mM: float,
    constants: Constants,
    Ksp_M3: float,
) -> float:
    """Return equilibrium Mg2P2O7(s) in mM formula units."""
    m0, p0, _, _ = solve_aqueous_speciation(Mg_total_mM, P2O7_total_mM, constants)
    if supersaturation(m0, p0, Ksp_M3) <= 1.0:
        return 0.0

    maximum = min(P2O7_total_mM, Mg_total_mM / 2.0)

    def saturation_residual(solid_mM: float) -> float:
        Mg_aq = Mg_total_mM - 2.0 * solid_mM
        P2O7_aq = P2O7_total_mM - solid_mM
        if Mg_aq <= 0.0 or P2O7_aq <= 0.0:
            return -Ksp_M3
        m, p, _, _ = solve_aqueous_speciation(Mg_aq, P2O7_aq, constants)
        return ion_product_M3(m, p) - Ksp_M3

    # At the stoichiometric maximum at least one aqueous reactant is exhausted,
    # so the residual is negative and the supersaturated no-solid state provides
    # the opposite sign at s=0.
    return brentq(saturation_residual, 0.0, maximum, maxiter=300)


def run_point(
    Mg_total_mM: float,
    P2O7_total_mM: float,
    config: ModelConfig = MODEL,
    constants: Constants | None = None,
) -> PointResult:
    """Calculate one no-solid or equilibrium point."""
    constants = constants or effective_constants(config)
    m0, p0, c10, c20 = solve_aqueous_speciation(Mg_total_mM, P2O7_total_mM, constants)
    S_initial = supersaturation(m0, p0, config.Ksp_M3)
    s_eq = solve_equilibrium_solid(
        Mg_total_mM, P2O7_total_mM, constants, config.Ksp_M3
    )

    if config.solid_model == "none":
        solid = 0.0
        m, p, c1, c2 = m0, p0, c10, c20
    elif config.solid_model == "equilibrium":
        solid = s_eq
        m, p, c1, c2 = speciation_after_solid(
            Mg_total_mM, P2O7_total_mM, solid, constants
        )
    else:
        raise ValueError("solid_model must be 'equilibrium' or 'none'.")

    return PointResult(
        Mg_total_mM=float(Mg_total_mM),
        P2O7_total_mM=float(P2O7_total_mM),
        Mg2_free_mM=m,
        P2O7_free_mM=p,
        MgP2O7_mM=c1,
        Mg2P2O7_aq_mM=c2,
        Mg2P2O7_initial_mM=c20,
        Mg2P2O7_s_mM=solid,
        S_initial=S_initial,
        S_final=supersaturation(m, p, config.Ksp_M3),
        s_eq_mM=s_eq,
    )


def p2o7_grid(config: ModelConfig = MODEL) -> np.ndarray:
    if config.p2o7_points < 2:
        raise ValueError("p2o7_points must be at least 2.")
    if config.p2o7_min_mM < 0.0 or config.p2o7_max_mM <= config.p2o7_min_mM:
        raise ValueError("P2O7 sweep limits are invalid.")
    return np.linspace(config.p2o7_min_mM, config.p2o7_max_mM, config.p2o7_points)


def compute_sweep(config: ModelConfig = MODEL) -> list[PointResult]:
    """Calculate the configured P2O7 sweep for every Mg total."""
    validate_config(config)
    constants = effective_constants(config)
    return [
        run_point(mg, p2o7, config, constants)
        for mg in config.mg_totals_mM
        for p2o7 in p2o7_grid(config)
    ]


# -----------------------------------------------------------------------------
# SCIENTIFIC CONSISTENCY CHECKS
# -----------------------------------------------------------------------------
def validate_config(config: ModelConfig = MODEL) -> None:
    """Reject invalid settings before calculation begins."""
    if config.solid_model not in {"none", "equilibrium"}:
        raise ValueError("solid_model must be 'equilibrium' or 'none'.")
    if not math.isfinite(config.temperature_C) or config.temperature_C <= -273.15:
        raise ValueError("temperature_C must be above absolute zero.")
    for name in (
        "log10_K1_ref",
        "log10_K2_ref",
        "dH_K1_kJ_mol",
        "dH_K2_kJ_mol",
        "Ksp_M3",
        "p2o7_min_mM",
        "p2o7_max_mM",
    ):
        if not math.isfinite(float(getattr(config, name))):
            raise ValueError(f"{name} must be finite.")
    if config.Ksp_M3 <= 0.0:
        raise ValueError("Ksp_M3 must be positive.")
    if not config.mg_totals_mM or any(
        (not math.isfinite(value) or value < 0.0)
        for value in config.mg_totals_mM
    ):
        raise ValueError("mg_totals_mM must contain finite non-negative values.")
    p2o7_grid(config)


def check_results(
    rows: Iterable[PointResult],
    config: ModelConfig = MODEL,
    tol: float = 1e-7,
) -> None:
    """Raise if mass balance, bounds, or equilibrium endpoints are inconsistent."""
    for row in rows:
        Mg_balance = (
            row.Mg2_free_mM
            + row.MgP2O7_mM
            + 2.0 * row.Mg2P2O7_aq_mM
            + 2.0 * row.Mg2P2O7_s_mM
        )
        P_balance = (
            row.P2O7_free_mM
            + row.MgP2O7_mM
            + row.Mg2P2O7_aq_mM
            + row.Mg2P2O7_s_mM
        )
        if abs(Mg_balance - row.Mg_total_mM) > tol:
            raise RuntimeError("Mg mass-balance check failed.")
        if abs(P_balance - row.P2O7_total_mM) > tol:
            raise RuntimeError("P2O7 mass-balance check failed.")
        maximum_solid = min(row.P2O7_total_mM, row.Mg_total_mM / 2.0)
        if row.Mg2P2O7_s_mM < -tol or row.Mg2P2O7_s_mM > maximum_solid + tol:
            raise RuntimeError("Solid amount is outside stoichiometric bounds.")
        if (
            config.solid_model == "equilibrium"
            and row.Mg2P2O7_s_mM > tol
            and abs(row.S_final - 1.0) > 5.0 * tol
        ):
            raise RuntimeError("A precipitating equilibrium point did not end at S=1.")


# -----------------------------------------------------------------------------
# PLOTTING
# -----------------------------------------------------------------------------
SERIES = {
    "Mg2_free_mM": "[Mg²⁺_free] (mM)",
    "P2O7_free_mM": "[P₂O₇⁴⁻_free] (mM)",
    "MgP2O7_mM": "[MgP₂O₇²⁻] (mM)",
    "Mg2P2O7_aq_mM": "[Mg₂P₂O₇(aq), final] (mM)",
    "Mg2P2O7_initial_mM": "[Mg₂P₂O₇(aq), initial] (mM)",
    "Mg2P2O7_s_mM": "[Mg₂P₂O₇(s)] (mM)",
    "S_initial": "S_initial",
    "S_final": "S_final",
}
SWEEP_XLABEL = "[P₂O₇⁴⁻]ₜₒₜₐₗ (mM)"


def _configure_font(plot: PlotConfig) -> None:
    """Apply shared serif typography for preview and figure export."""
    serif_families = (
        ["Times New Roman", "DejaVu Serif"]
        if plot.use_times_new_roman
        else ["DejaVu Serif"]
    )
    mpl.rcParams.update({
        "font.family": "serif",
        "font.serif": serif_families,
        "pdf.fonttype": 42,
        "ps.fonttype": 42,
        "svg.fonttype": "path",
    })


def _text_weight(plot: PlotConfig) -> str:
    return "bold" if plot.bold_text else "normal"


def _style_axis(ax: plt.Axes, plot: PlotConfig) -> None:
    """Apply shared axis-label and tick typography to preview/export axes."""
    weight = _text_weight(plot)
    ax.xaxis.label.set_fontsize(plot.axis_label_size)
    ax.yaxis.label.set_fontsize(plot.axis_label_size)
    ax.xaxis.label.set_fontweight(weight)
    ax.yaxis.label.set_fontweight(weight)
    ax.tick_params(
        axis="both",
        labelsize=plot.tick_label_size,
        width=plot.tick_width,
        length=plot.tick_length,
    )
    for spine in ax.spines.values():
        spine.set_linewidth(plot.spine_width)
    for label in (*ax.get_xticklabels(), *ax.get_yticklabels()):
        label.set_fontweight(weight)


def _add_legend(ax: plt.Axes, plot: PlotConfig, **kwargs) -> None:
    """Add a legend using the shared plot typography."""
    legend = ax.legend(
        prop={
            "family": "serif",
            "size": plot.legend_size,
            "weight": _text_weight(plot),
        },
        **kwargs,
    )
    if legend is not None and legend.get_title() is not None:
        legend.get_title().set_fontsize(plot.legend_title_size)
        legend.get_title().set_fontweight(_text_weight(plot))


def selected_series(plot: PlotConfig = PLOT) -> list[str]:
    """Return enabled sweep outputs in a stable display order."""
    flags = {
        "Mg2_free_mM": plot.show_Mg2_free,
        "P2O7_free_mM": plot.show_P2O7_free,
        "MgP2O7_mM": plot.show_MgP2O7,
        "Mg2P2O7_aq_mM": plot.show_Mg2P2O7_aq,
        "Mg2P2O7_initial_mM": plot.show_Mg2P2O7_initial,
        "Mg2P2O7_s_mM": plot.show_solid,
        "S_initial": plot.show_S_initial,
        "S_final": plot.show_S_final,
    }
    return [name for name, enabled in flags.items() if enabled]


def _rows_by_mg(rows: Sequence[PointResult]) -> dict[float, list[PointResult]]:
    grouped: dict[float, list[PointResult]] = {}
    for row in rows:
        grouped.setdefault(row.Mg_total_mM, []).append(row)
    return grouped


def _validate_plot_config(plot: PlotConfig) -> None:
    if not plot.show_lines and not plot.show_markers:
        raise ValueError("Enable show_lines, show_markers, or both.")
    if not plot.series_colors:
        raise ValueError("series_colors must contain at least one color.")
    if plot.show_markers and not plot.marker_styles:
        raise ValueError("marker_styles must contain at least one marker when markers are enabled.")
    if plot.marker_every < 1:
        raise ValueError("marker_every must be a positive integer.")
    if plot.marker_size <= 0.0 or plot.line_width <= 0.0:
        raise ValueError("marker_size and line_width must be positive.")
    if plot.figure_width <= 0.0 or plot.figure_height <= 0.0 or plot.dpi <= 0:
        raise ValueError("Figure dimensions and dpi must be positive.")
    text_sizes = (
        plot.axis_label_size,
        plot.tick_label_size,
        plot.legend_size,
        plot.legend_title_size,
    )
    if min(text_sizes) <= 0.0:
        raise ValueError("Text sizes must be positive.")
    frame_sizes = (
        plot.spine_width,
        plot.tick_width,
        plot.tick_length,
        plot.grid_line_width,
    )
    if min(frame_sizes) <= 0.0:
        raise ValueError("Spine, tick, and grid dimensions must be positive.")
    if not (0.0 <= plot.grid_alpha <= 1.0):
        raise ValueError("grid_alpha must be between 0 and 1.")


def _series_color(
    plot: PlotConfig, series_index: int
) -> str | tuple[float, float, float]:
    """Return the configured series-color sequence for any number of curves."""
    if series_index < len(plot.series_colors):
        return plot.series_colors[series_index]
    rng = np.random.RandomState(plot.extra_color_seed)
    extras = rng.rand(series_index - len(plot.series_colors) + 1, 3)
    return tuple(float(value) for value in extras[-1])


def _series_marker(plot: PlotConfig, series_index: int) -> str:
    """Return the configured marker sequence, cycling additional markers safely."""
    markers = plot.marker_styles
    if series_index < min(3, len(markers)):
        return markers[series_index]
    extras = markers[3:] or markers
    return extras[(series_index - 3) % len(extras)]


def _variable_linestyle(plot: PlotConfig, variable: str) -> str:
    """Return the user-configured line style for a plotted variable."""
    style_map = {
        "Mg2_free_mM": plot.linestyle_Mg2_free,
        "P2O7_free_mM": plot.linestyle_P2O7_free,
        "MgP2O7_mM": plot.linestyle_MgP2O7,
        "Mg2P2O7_aq_mM": plot.linestyle_Mg2P2O7_aq,
        "Mg2P2O7_initial_mM": plot.linestyle_Mg2P2O7_initial,
        "Mg2P2O7_s_mM": plot.linestyle_solid,
        "S_initial": plot.linestyle_S_initial,
        "S_final": plot.linestyle_S_final,
    }
    return style_map.get(variable, "-")


def _line_kwargs(
    plot: PlotConfig,
    series_index: int,
    variable: str | None = None,
    *,
    allow_markers: bool = True,
) -> dict[str, object]:
    """Return shared line/marker settings for one plotted series."""
    marker = None
    markevery = None
    if plot.show_markers and allow_markers:
        marker = _series_marker(plot, series_index)
        markevery = plot.marker_every
    linestyle = _variable_linestyle(plot, variable) if plot.show_lines else "None"
    return {
        "color": _series_color(plot, series_index),
        "linewidth": plot.line_width,
        "linestyle": linestyle,
        "marker": marker,
        "markevery": markevery,
        "markersize": plot.marker_size,
    }


def _draw_sweep_variable(
    ax: plt.Axes,
    name: str,
    grouped: Mapping[float, Sequence[PointResult]],
    plot: PlotConfig,
) -> None:
    for index, (mg, group) in enumerate(grouped.items()):
        ax.plot(
            [row.P2O7_total_mM for row in group],
            [getattr(row, name) for row in group],
            label=f"{mg:g} mM Mg²⁺",
            **_line_kwargs(
                plot,
                index,
                name,
                allow_markers=(
                    name not in {"S_initial", "S_final"} or not plot.show_lines
                ),
            ),
        )
    if name in {"S_initial", "S_final"}:
        ax.axhline(1.0, color="gray", linestyle=":", alpha=0.5)
    ax.set_xlabel(SWEEP_XLABEL)
    ax.set_ylabel(SERIES[name])
    ax.grid(True, linewidth=plot.grid_line_width, alpha=plot.grid_alpha)
    _style_axis(ax, plot)
    _add_legend(ax, plot)


def plot_sweep_preview(
    rows: Sequence[PointResult],
    config: ModelConfig = MODEL,
    plot: PlotConfig = PLOT,
) -> list[plt.Figure]:
    """Create 2×2 preview pages; unused cells on the last page are hidden."""
    del config  # presentation is intentionally title-free
    _validate_plot_config(plot)
    _configure_font(plot)
    names = selected_series(plot)
    if not names:
        raise ValueError("At least one plotted output must be enabled.")
    grouped = _rows_by_mg(rows)
    pages: list[plt.Figure] = []
    for offset in range(0, len(names), 4):
        fig, axes = plt.subplots(
            2, 2,
            figsize=(2.0 * plot.figure_width, 2.0 * plot.figure_height),
            squeeze=False,
        )
        flat = axes.ravel()
        chunk = names[offset:offset + 4]
        for ax, name in zip(flat, chunk):
            _draw_sweep_variable(ax, name, grouped, plot)
        for ax in flat[len(chunk):]:
            ax.set_visible(False)
        fig.tight_layout()
        pages.append(fig)
    return pages


def plot_sweep(
    rows: Sequence[PointResult],
    config: ModelConfig = MODEL,
    plot: PlotConfig = PLOT,
) -> dict[str, plt.Figure]:
    """Create one title-free standalone figure per enabled output for export."""
    del config
    _validate_plot_config(plot)
    _configure_font(plot)
    names = selected_series(plot)
    if not names:
        raise ValueError("At least one plotted output must be enabled.")
    grouped = _rows_by_mg(rows)
    figures: dict[str, plt.Figure] = {}
    for name in names:
        fig, ax = plt.subplots(figsize=(plot.figure_width, plot.figure_height))
        _draw_sweep_variable(ax, name, grouped, plot)
        fig.tight_layout()
        figures[name] = fig
    return figures


def save_sweep_figures(
    figures: Mapping[str, plt.Figure],
    plot: PlotConfig = PLOT,
    export: ExportConfig = EXPORT,
) -> list[Path]:
    """Save each standalone sweep figure to an independent file."""
    output_dir = Path(export.directory)
    output_dir.mkdir(parents=True, exist_ok=True)
    paths: list[Path] = []
    for name, fig in figures.items():
        path = output_dir / f"{export.figure_stem}_{name}.{export.figure_format.lower()}"
        fig.savefig(path, dpi=plot.dpi, bbox_inches="tight")
        paths.append(path)
    return paths


# -----------------------------------------------------------------------------
# EXPORTS
# -----------------------------------------------------------------------------
def _metadata(config: ModelConfig, constants: Constants) -> Mapping[str, object]:
    return {
        "solid_model": config.solid_model,
        "temperature_C": config.temperature_C,
        "use_K1K2_temperature_correction": config.use_K1K2_temperature_correction,
        "log10_K1_effective_M_inv": math.log10(constants.K1_M_inv),
        "log10_K2_effective_M_inv": math.log10(constants.K2_M_inv),
        "Ksp_M3": config.Ksp_M3,
        "log10_Ksp": math.log10(config.Ksp_M3),
        "precursor_threshold_mM": constants.precursor_threshold_mM,
    }


def export_table(
    rows: Sequence[PointResult],
    config: ModelConfig = MODEL,
    export: ExportConfig = EXPORT,
) -> Path:
    """Write CSV, TSV, or XLSX with run metadata repeated on every row."""
    constants = effective_constants(config)
    records = [{**_metadata(config, constants), **asdict(row)} for row in rows]
    if not records:
        raise ValueError("rows must contain at least one result for table export.")
    output_dir = Path(export.directory)
    output_dir.mkdir(parents=True, exist_ok=True)
    fmt = export.table_format.lower()
    path = output_dir / f"{export.table_stem}.{fmt}"

    if fmt in {"csv", "tsv"}:
        delimiter = "," if fmt == "csv" else "\t"
        with path.open("w", newline="", encoding="utf-8") as handle:
            writer = csv.DictWriter(handle, fieldnames=list(records[0]), delimiter=delimiter)
            writer.writeheader()
            writer.writerows(records)
        return path

    if fmt == "xlsx":
        try:
            from openpyxl import Workbook
        except ImportError as exc:
            raise RuntimeError("XLSX export requires openpyxl.") from exc
        workbook = Workbook()
        sheet = workbook.active
        sheet.title = "sweep"
        sheet.append(list(records[0]))
        for record in records:
            sheet.append([record[key] for key in records[0]])
        workbook.save(path)
        return path

    raise ValueError("table_format must be 'csv', 'tsv', or 'xlsx'.")


# -----------------------------------------------------------------------------
# MAIN
# -----------------------------------------------------------------------------
def main() -> None:
    rows = compute_sweep(MODEL)
    check_results(rows, MODEL)
    constants = effective_constants(MODEL)

    print("Mg–P₂O₇ equilibrium model")
    print(f"  solid_model = {MODEL.solid_model}")
    print(f"  temperature = {MODEL.temperature_C:g} °C")
    print(f"  log10 K1 effective = {math.log10(constants.K1_M_inv):.6f}")
    print(f"  log10 K2 effective = {math.log10(constants.K2_M_inv):.6f}")
    print(f"  log10 Ksp = {math.log10(MODEL.Ksp_M3):.6f}")
    print(f"  precursor threshold = {constants.precursor_threshold_mM:.6g} mM")

    if PLOT.enabled:
        plot_sweep_preview(rows, MODEL, PLOT)

    if EXPORT.export_figures:
        export_figures = plot_sweep(rows, MODEL, PLOT)
        for path in save_sweep_figures(export_figures, PLOT, EXPORT):
            print(f"  figure: {path}")
        for figure in export_figures.values():
            plt.close(figure)

    if EXPORT.export_table:
        print(f"  table: {export_table(rows, MODEL, EXPORT)}")

    if PLOT.enabled:
        plt.show()


if __name__ == "__main__":
    main()
```
