## Supplementary Code (Python) for "Mechanistic Insights into Magnesium Pyrophosphate Formation in the Presence of Gold Nanoclusters Enable Genetic Analysis via Co-Aggregation-Induced Fluorescence Enhancement": README.md

# Mg²⁺–P₂O₇⁴⁻ Equilibrium Speciation and Precipitation Model

`mg_p2o7_equilibrium.py` calculates concentration-based Mg²⁺–P₂O₇⁴⁻ aqueous speciation and the equilibrium amount of Mg₂P₂O₇(s). It supports concentration sweeps, publication-style plotting, and numerical export while keeping the thermodynamic model explicit and compact.

## 1. Scope and assumptions

The aqueous network contains:

- free Mg²⁺
- free P₂O₇⁴⁻
- MgP₂O₇²⁻
- Mg₂P₂O₇(aq)

The equilibrium solid treatment adds Mg₂P₂O₇(s), reported in mM formula units.

Two solid modes are available:

- `solid_model="equilibrium"`: calculate the equilibrium Mg₂P₂O₇(s) amount.
- `solid_model="none"`: suppress solid formation while still reporting the counterfactual equilibrium amount as `s_eq_mM`.

The model does **not** explicitly include pH-dependent pyrophosphate protonation, activity coefficients, ionic-strength iteration, buffers, competing ligands, alternative solid phases, precipitation kinetics, crystal-size/population balances, aggregation, or morphology prediction. The P₂O₇ input is therefore the P₂O₇⁴⁻ pool represented by this model, not an internally calculated analytical-total pyrophosphate distribution across protonation states.

## 2. Requirements

- Python 3.10+
- NumPy `>=1.24,<3`
- SciPy `>=1.10,<2`
- Matplotlib `>=3.7,<4`
- openpyxl `>=3.1,<4` for XLSX export

Install dependencies with:

```bash
python -m pip install -r requirements.txt
```

## 3. Running the model

```bash
python mg_p2o7_equilibrium.py
```

Importing the module does not automatically calculate, plot, or write files. In a standalone run, preview plotting is enabled by default, while file exports are disabled (`export_table=False`, `export_figures=False`) until explicitly enabled in `EXPORT`. On a headless system, set `PLOT.enabled=False` when no interactive preview window is available, or enable figure export instead.

## 4. User configuration

The main settings are grouped near the beginning of the script in `ModelConfig`, `PlotConfig`, and `ExportConfig`. For a standalone run, edit the instantiated `MODEL`, `PLOT`, and `EXPORT` objects. For programmatic use, create new dataclass instances or use `dataclasses.replace()`.

### Model settings

| Parameter | Default | Meaning |
|---|---:|---|
| `solid_model` | `"equilibrium"` | `"equilibrium"` or `"none"` |
| `temperature_C` | `65.0` | calculation temperature; 25 °C is the reference temperature of the source K₁/K₂ values |
| `use_K1K2_temperature_correction` | `True` | apply van ’t Hoff transport to K₁/K₂ |
| `log10_K1_ref` | `5.42` | log₁₀K₁ at 25 °C, K₁ in M⁻¹ |
| `log10_K2_ref` | `2.33` | log₁₀K₂ at 25 °C, K₂ in M⁻¹ |
| `dH_K1_kJ_mol` | `12.552` | association-direction enthalpy used for K₁ transport |
| `dH_K2_kJ_mol` | `0.0` | zero disables K₂ temperature transport in this parameterization |
| `Ksp_M3` | `1.63e-13` | fixed concentration-form Ksp in M³ |
| `mg_totals_mM` | `(4, 8, 12)` | Mg-total values used in the sweep |
| `p2o7_min_mM` | `0.01` | lower P₂O₇ sweep value |
| `p2o7_max_mM` | `10.0` | upper P₂O₇ sweep value |
| `p2o7_points` | `100` | number of P₂O₇ sweep points |

`mg_totals_mM` can contain any non-empty set of finite non-negative Mg²⁺ totals, for example:

```python
mg_totals_mM=(2.0, 4.0, 6.0, 8.0, 10.0, 12.0)
```

The P₂O₇ range and numerical resolution are controlled independently by `p2o7_min_mM`, `p2o7_max_mM`, and `p2o7_points`.

### Plot settings

Preview figures are arranged in **2×2 pages**, with four parameters per window and unused cells hidden on the final page. Different parameters do not share an axis. Exported figures are **standalone**, one file per enabled parameter. Plot titles are intentionally omitted; each y-axis identifies the plotted quantity directly.

Important shared controls include:

- `show_lines`, `line_width`
- `series_colors`, `extra_color_seed`
- `show_markers`, `marker_styles`, `marker_every`, `marker_size`
- the variable-specific `linestyle_*` settings
- `figure_width`, `figure_height`, `dpi`
- `spine_width`, `tick_width`, `tick_length`
- `grid_line_width`, `grid_alpha`
- `use_times_new_roman`
- `axis_label_size`, `tick_label_size`, `legend_size`, `legend_title_size`
- `bold_text`

The default series order starts with black/red/blue and square/circle/triangle-up markers. Additional colors are generated deterministically when needed, and additional markers cycle safely, so the number of Mg²⁺ series is not limited by the default style arrays.

Line-only, symbol-only, and line+symbol plotting are supported through `show_lines` and `show_markers`. Supersaturation sweeps omit markers during ordinary line plotting, but remain visible in marker-only mode.

The `show_*` flags select which sweep outputs are previewed/exported:

- `show_Mg2_free`
- `show_P2O7_free`
- `show_MgP2O7`
- `show_Mg2P2O7_aq`
- `show_Mg2P2O7_initial`
- `show_solid`
- `show_S_initial`
- `show_S_final`

### Export settings

`ExportConfig` controls the output directory, numerical-table format (`csv`, `tsv`, or `xlsx`), figure format (for example `png`, `pdf`, or `svg`), and filename stems. The output directory is created automatically when an export is requested.

## 5. Model equations and numerical solution

### Aqueous association and temperature transport

The represented association equilibria are:

```text
Mg²⁺ + P₂O₇⁴⁻ ⇌ MgP₂O₇²⁻             K₁
MgP₂O₇²⁻ + Mg²⁺ ⇌ Mg₂P₂O₇(aq)         K₂
```

At the 25 °C reference temperature:

```text
log10(K₁ / M⁻¹) = 5.42
log10(K₂ / M⁻¹) = 2.33
```

Using

```text
m  = [Mg²⁺]
p  = [P₂O₇⁴⁻]
c₁ = [MgP₂O₇²⁻]
c₂ = [Mg₂P₂O₇(aq)]
```

gives:

```text
c₁ = K₁mp
c₂ = K₁K₂m²p
```

The source constants are in M⁻¹; the solver works in mM, so each stepwise association constant is multiplied internally by `1e-3` before use in the mM equations.

When temperature correction is enabled, K₁ and K₂ are transported from 25 °C with:

```text
K(T) = Kref × exp[-(ΔH/R)(1/T - 1/Tref)]
```

where `Tref = 298.15 K`, `R = 8.31446261815324 J mol⁻¹ K⁻¹`, `ΔH₁ = 12.552 kJ mol⁻¹`, and `ΔH₂ = 0`. The enthalpy signs follow the association reactions written above. Ksp is kept fixed.

### Mass balances and Brent solve

Before solid formation:

```text
Mtot = m + c₁ + 2c₂
Ptot = p + c₁ + c₂
```

The P₂O₇ balance gives:

```text
p = Ptot / (1 + K₁m + K₁K₂m²)
```

which reduces the aqueous problem to one scalar Mg²⁺ mass-balance equation:

```text
f(m) = m + K₁mp + 2K₁K₂m²p - Mtot = 0
```

`solve_aqueous_speciation()` solves this equation with SciPy's `scipy.optimize.brentq` implementation of **Brent's bracketed root-finding method**. The physical solution satisfies `0 ≤ m ≤ Mtot`; the implementation uses a conservative non-negative upper bound that contains this interval. Once `m` is found, `p`, `c₁`, and `c₂` follow analytically.

### Supersaturation and equilibrium solid

The concentration-form solubility boundary is:

```text
Ksp = [Mg²⁺]²[P₂O₇⁴⁻]
Ksp = 1.63 × 10⁻13 M³
```

The free-ion product and supersaturation ratio are:

```text
Q = [Mg²⁺]²[P₂O₇⁴⁻]
S = Q / Ksp
```

Thus `S < 1`, `S = 1`, and `S > 1` denote under-, exactly-, and supersaturated states within this concentration-based model.

Because `c₂ = K₁K₂Q`, the corresponding aqueous Mg₂P₂O₇ threshold is derived as:

```text
c₂,eq = K₁K₂Ksp
```

and is reported as `precursor_threshold_mM`.

If the no-solid state is supersaturated and `solid_model="equilibrium"`, formation of `s` mM Mg₂P₂O₇(s) leaves:

```text
Maq = Mtot - 2s
Paq = Ptot - s
0 ≤ s ≤ min(Ptot, Mtot/2)
```

A second Brent solve finds the solid amount for which the re-equilibrated aqueous phase satisfies `Q = Ksp`. In `solid_model="none"`, no solid is applied to the returned state, although the same equilibrium amount is still reported as `s_eq_mM`.

## 6. Result fields

Each `PointResult` contains:

| Field | Meaning |
|---|---|
| `Mg_total_mM` | input Mg total |
| `P2O7_total_mM` | input P₂O₇ total represented by the model |
| `Mg2_free_mM` | final free Mg²⁺ |
| `P2O7_free_mM` | final free P₂O₇⁴⁻ |
| `MgP2O7_mM` | final MgP₂O₇²⁻ |
| `Mg2P2O7_aq_mM` | final Mg₂P₂O₇(aq) |
| `Mg2P2O7_initial_mM` | no-solid Mg₂P₂O₇(aq) value |
| `Mg2P2O7_s_mM` | selected solid amount |
| `S_initial` | no-solid supersaturation ratio |
| `S_final` | final supersaturation ratio |
| `s_eq_mM` | independently calculated equilibrium solid amount |

## 7. Sweeps, plots, and exports

`compute_sweep()` evaluates the configured P₂O₇ grid for every Mg total in `mg_totals_mM`.

`plot_sweep_preview()` creates title-free 2×2 preview pages. `plot_sweep()` creates the corresponding standalone figures used for export or programmatic access. Preview and export use the same drawing/styling logic.

Axis labels and legends use direct Unicode chemistry notation (for example, `Mg²⁺`, `P₂O₇⁴⁻`, and `MgP₂O₇²⁻`) for compact, natural-looking typesetting. Times New Roman is preferred when enabled and available; DejaVu Serif is the portable fallback. The `bold_text` option controls the shared text weight. SVG exports convert text to vector paths for portable appearance, and PDF exports use TrueType embedding.

Numerical data can be exported as CSV, TSV, or XLSX. Figures can be exported in any Matplotlib-supported format. Exported table rows include run metadata such as the effective K₁/K₂ values, Ksp, and the derived precursor threshold.

## 8. Validation and consistency checks

The script rejects invalid modes, non-finite or negative totals, invalid sweep limits, invalid grid sizes, non-positive Ksp, and non-positive/non-finite effective association constants.

`check_results()` verifies Mg and P₂O₇ mass balance, solid stoichiometric bounds, and the expected `S ≈ 1` endpoint for precipitating equilibrium states. Small residuals at floating-point/root-solver precision are expected.

## 9. Programmatic use

```python
from dataclasses import replace
import mg_p2o7_equilibrium as model

config = replace(
    model.MODEL,
    temperature_C=65.0,
    mg_totals_mM=(2.0, 4.0, 6.0, 8.0, 10.0, 12.0),
    p2o7_min_mM=0.1,
    p2o7_max_mM=8.0,
    p2o7_points=81,
)

rows = model.compute_sweep(config)
model.check_results(rows, config)

constants = model.effective_constants(config)
point = model.run_point(8.0, 4.0, config, constants)
print(point)
```

For programmatic plotting, `plot_sweep_preview()` returns the 2×2 preview pages and `plot_sweep()` returns a dictionary of standalone figures keyed by result-field name.

## 10. References and parameter roles

1. **Kern, J. A.; Davis, R. H.** Application of Solution Equilibrium Analysis to In Vitro RNA Transcription. *Biotechnology Progress* **1997**, *13*(6), 747–756. **DOI:** `10.1021/bp970094p`. Role: conditional Mg–pyrophosphate association constants used for K₁ and K₂.

2. **Ahmadi, S.; Stover, N. M.; Ganko, K.; Kayser, F. B.; Braatz, R. D.; Myerson, A. S.** DNA Nanoflowers Assemble via Template-Induced Crystallization during mRNA Synthesis. *Crystal Growth & Design* **2026**, *26*(5), 2108–2117. **DOI:** `10.1021/acs.cgd.5c01710`. Role: public implementation of Mg–P₂O₇ temperature transport and the fitted Mg₂P₂O₇·3.5H₂O solubility parameterization. That implementation stores the MgPPi enthalpy in the dissociation direction; this script uses the equivalent association-direction value. It also fits temperature dependence for Ksp, whereas this compact equilibrium script intentionally keeps `Ksp_M3` fixed at its 25 °C value.

The references support specific parameter choices and modeling elements; they should not be interpreted as validating model components explicitly outside the scope stated in Section 1. The code is distributed under the MIT License; see `LICENSE`.
